# Multi-omic spatial mapping of the human habenula defines the molecular and anatomical organization of medial and lateral subregions

**DOI:** 10.64898/2026.09.30.755667

**Authors:** Kelsey D. Montgomery, Jonathan M. Werner, Nicholas J. Eagles, Chunyu Liu, Cynthia S. Cardinault, Svitlana V. Bach, Atharv Chandra, Ruth Zhang, Sarah E. Maguire, Heena R. Divecha, Amy Deep-Soboslay, Ryan A. Miller, Louise A. Huuki-Myers, Rahul A. Bharadwaj, Joel E. Kleinman, Thomas M. Hyde, Brion Maher, Leonardo Collado-Torres, Kristen R. Maynard

**Affiliations:** Lieber Institute for Brain Development, Johns Hopkins Medical Campus, Baltimore, MD, 21205, USA; Department of Mental Health, Bloomberg School of Public Health, Johns Hopkins University, Baltimore, MD, 21205, USA; Department of Psychiatry and Behavioral Sciences, Johns Hopkins School of Medicine, Baltimore, MD, 21205, USA; UK Dementia Research Institute at The University of Cambridge, Cambridge, CB2 0AH, UK; Department of Clinical Neurosciences, School of Clinical Medicine, The University of Cambridge, Cambridge, CB2 0QQ, UK; Department of Neurology, Johns Hopkins School of Medicine, Baltimore, MD, 21205, USA; Department of Biostatistics, Johns Hopkins Bloomberg School of Public Health, Baltimore, MD, 21205, USA; Center for Computational Biology, Johns Hopkins University, Baltimore, MD, 21205, USA; The Solomon H. Snyder Department of Neuroscience, Johns Hopkins School of Medicine, Baltimore, MD, 21205, USA

**Author notes:** equal contributors (Kelsey D. Montgomery, Jonathan M. Werner, Nicholas J. Eagles).

## Abstract

The habenula (Hb) is a small epithalamic midline structure critical for reward processing and mood regulation that is implicated in many complex brain disorders, including depression and addiction. The molecular anatomy of the Hb has been well-characterized in model organisms, but little is known about its organization in the human brain, including the transcriptomic and epigenetic signatures of spatially-organized lateral (LHb) and medial (MHb) cell types. Here, we generated a high resolution spatiomolecular map of the human Hb using both multiomic single nucleus sequencing (snMultiome) and spatially-resolved transcriptomics. We measured gene expression and chromatin accessibility across 4 MHb and 5 LHb neuronal cell types as defined by transcriptomic conservation with rodents and zebrafish. We identified cell type-specific molecular specializations related to GABAergic and glutamatergic signaling as well as regulatory relationships between open chromatin regions, transcription factors, and target genes associated with depression and addiction. High resolution spatial transcriptomics identified the anatomical location of heterogeneous LHb subpopulations, including two putative inhibitory populations in the LHb. We further characterized cell-type relationships between Hb neurons and glial subtypes and predicted ligand-receptor interactions between LHb and MHb neurons. Finally, we delineated Hb cell types and spatial domains enriched in genes associated with genetic risk for psychiatric disorders, identifying the strongest enrichment in the MHb. Collectively, the generated data and analyses resolve the spatiomolecular organization of the human Hb and identify molecular-genetic associations with neuropsychiatric disease.

## 1 Introduction

The habenula (Hb) is a pea-sized epithalamic structure that acts as a critical reward circuit hub regulating motivated behaviors and affective states (Namboodiri et al. 2016). In mammals, the Hb is divided into medial (MHb) and lateral (LHb) subregions that contain functionally distinct cell types and topographically organized projections (Ables et al. 2023; Hu et al. 2020). The MHb primarily projects to the interpeduncular nucleus (IPN) and MHb cell types play a role in nicotine addiction and withdrawal-associated negative affect (Baldwin et al. 2011). The LHb is connected to monoaminergic and catecholaminergic hubs, such as the ventral tegmental area (VTA), locus coeruleus (LC), and dorsal raphe nucleus (DRN) (Herkenham and Nauta 1979; Nakamura et al. 2024), and is activated in response to aversive stimuli and absence of an anticipated reward (Matsumoto and Hikosaka 2007; Stamatakis and Stuber 2012).

Hb dysregulation is implicated in several neuropsychiatric disorders, including schizophrenia (SCZ), major depressive disorder (MDD), and substance use disorders (SUD) (Hu et al. 2020; Velasquez et al. 2014; F. Lin et al. 2026; Yalcinbas et al. 2025; Savitz et al. 2011). The Hb is particularly relevant in the context of opioid use disorder (OUD) given high expression of *OPRM1,* which encodes the mu opioid receptor (MOR) (Popat et al. 2026; Gardon et al. 2014). Indeed, Hb neurons expressing MOR mediate negative affect and anxiety through separate projections to the IPN and DRN (Bailly et al. 2023). In postmortem human Hb, reduced Hb volume and neuron numbers have been reported in donors with heroin addiction (Müller et al. 2021). Altered Hb activity and volume have also been linked to MDD through neuroimaging studies (Savitz et al. 2011; F. Lin et al. 2026; Fortin et al. 2025; R. P. Lawson et al. 2017). Consistent with these findings, aberrant LHb neuron burst firing is associated with depressive-like symptoms in mice (Y. Cui et al. 2018; Yang, Cui, et al. 2018; Bray 2018). Modulation of Hb activity is emerging as a potential therapy for neuropsychiatric disorders (Hu et al. 2020). In humans, deep brain stimulation of Hb reduces depressive symptoms in MDD (Z. Wang et al. 2024) and in mice the NMDA receptor antagonist ketamine disrupts LHb burst firing to alleviate depressive-like symptoms (Yang, Cui, et al. 2018).

The anatomical organization and cellular heterogeneity of the Hb has been appreciated for decades with early studies resolving MHb and LHb subdivisions using classic histological techniques (Marburg 1944; Andres et al. 1999; Díaz et al. 2011; Aizawa et al. 2012). Advancements in single-cell (sc) and single-nucleus (sn) RNA sequencing (sc/snRNA-seq) technologies further revealed transcriptionally distinct and spatially organized MHb and LHb cell types in mice and zebrafish, many of which show unique connectivity, electrophysiology, and functional engagement (Pandey et al. 2018; Hashikawa et al. 2020; Wallace et al. 2020). While the Hb is primarily glutamatergic, some studies have reported expression of GABAergic markers (L. Zhang et al. 2018; Flanigan et al. 2020). However, whether these cells can release GABA and function as inhibitory neurons remains unclear. The MHb contains Substance P and cholinergic neurons, and the LHb contains a heterogeneous set of glutamatergic cell types that express a diversity of neuropeptides (Ables et al. 2023; Zahm and Root 2017; Andres et al. 1999; Geisler et al. 2003; Aizawa et al. 2012). Astrocytes also show diversity across the Hb (Gampe et al. 2012) and play important roles in supporting Hb neuron function. For example, neuronal-astrocyte interactions regulate LHb neuron burst firing, and disruption of neuronal-astrocyte signaling mediates depressive- and anxiety-like behaviors in mice (W. Cui et al. 2014; Y. Cui et al. 2018; Xin et al. 2025; Song et al. 2025; Tan et al. 2024; Aizawa et al. 2025). Altered Hb neuron-glia dynamics are also implicated in alcohol withdrawal (Bosque-Cordero et al. 2026).

While snRNA-seq resolved major cell types in human Hb (Yalcinbas et al. 2025), their anatomical organization is not defined and how epigenetic regulatory mechanisms impact LHb and MHb cell type specialization is not known. Rodent studies highlight the functional importance of Hb cellular topography by linking spatial patterning of functionally distinct LHb and MHb cell types with discrete projections to downstream target regions (Ables et al. 2023; Hu et al. 2020). Furthermore, given challenges in resolving transcriptomically similar glutamatergic LHb neuron subtypes, epigenetic characterization of cell type-specific gene regulatory networks, including open chromatin regions, transcription factors, and their downstream targets, may clarify putative functions or disease associations for spatially organized LHb neuron subtypes projecting to aminergic centers.

Here, we generated a high resolution multi-omic spatiomolecular map of the human Hb. We measured gene expression and chromatin accessibility across evolutionarily conserved LHb and MHb cell types and identified cell type-specific relationships between transcription factors, target genes, and candidate cis-regulatory elements (cCREs). We also generated high resolution transcriptome-wide SRT data, which we used to define cell-type proximity relationships and putative ligand-receptor signaling between LHb, MHb, and glial subpopulations. We integrated these single cell and SRT datasets with genome-wide association studies (GWAS) of brain disorders, revealing the MHb as particularly enriched for genes associated with risk for neuropsychiatric disorders and addiction. Collectively, this study defines the molecular and cellular organization of the human habenula and its relevance to neuropsychiatric disease.

## 2 Results

### 2.1 Molecularly-defined neuronal and glial cell types in the human habenula

To generate a spatiomolecular map of the human Hb (**Fig. 1A-D**, **Fig. S1**), we first needed to confirm consistent collection of this small midline structure (∼3-9 mm^3^) (**Fig. 1E**) from postmortem human brain tissue. Tissue blocks putatively containing the Hb were dissected from fresh frozen brain slabs using neuroanatomical landmarks, including the segmented putamen (SG), lateral geniculate nucleus (LGN) and substantia nigra (SN) (**Fig. S2**). Because of its small size, confirming full inclusion of the Hb on the tissue blocks using neuroanatomical landmarks alone is insufficient. To anatomically validate the presence of MHb and LHb, we performed RNAscope single molecule fluorescence in situ hybridization (RNAscope/smFISH) on tissue sections from the blocks with probes against Hb marker genes, including *TAC3* (MHb), *GPR151* (MHb, LHb), and *POU4F1* (MHb, LHb; **Fig. 1F**, **Fig. S1, Fig. S2**) prior to data collection. After validating Hb capture, we next performed single nucleus multiome sequencing (snMultiome: RNA-seq and ATAC-seq) and Visium High Definition (VisiumHD) SRT of neurotypical control brain donors (n=10 and 5 donors, capturing a total of 54,193 and 287,325 cells, respectively; **Fig. 1A-D**, **Fig. S1, Table S1**, **Table S2**, **Methods 4.1-4.3**). snMultiome data was collected across the anterior-posterior (A-P) axis, whereas SRT data were collected at an intermediate level of the Hb AP axis to capture both MHb and LHb (**Fig. S1**), which was confirmed by robust expression of Hb marker genes (**Fig. 1G**). VisiumHD enables 2µm bin resolution SRT quantifications that can be assigned to individual cells by segmenting the H&E images, while snMultiome allows quantifying both open chromatin and RNA expression on the same nuclei. We integrated SRT and snMultiome data to characterize the molecular and spatial organization of LHb and MHb cell populations conserved across human (Yalcinbas et al. 2025), mouse (Hashikawa et al. 2020; Wallace et al. 2020), and zebrafish (Pandey et al. 2018). We leveraged this integrated cross-species consensus map to assess Hb cell type-specific gene regulation, cellular communication, and enrichment of genes associated with risk for neuropsychiatric and substance use disorders (SUDs) (**Fig. 1D**).

**Fig. 1:**
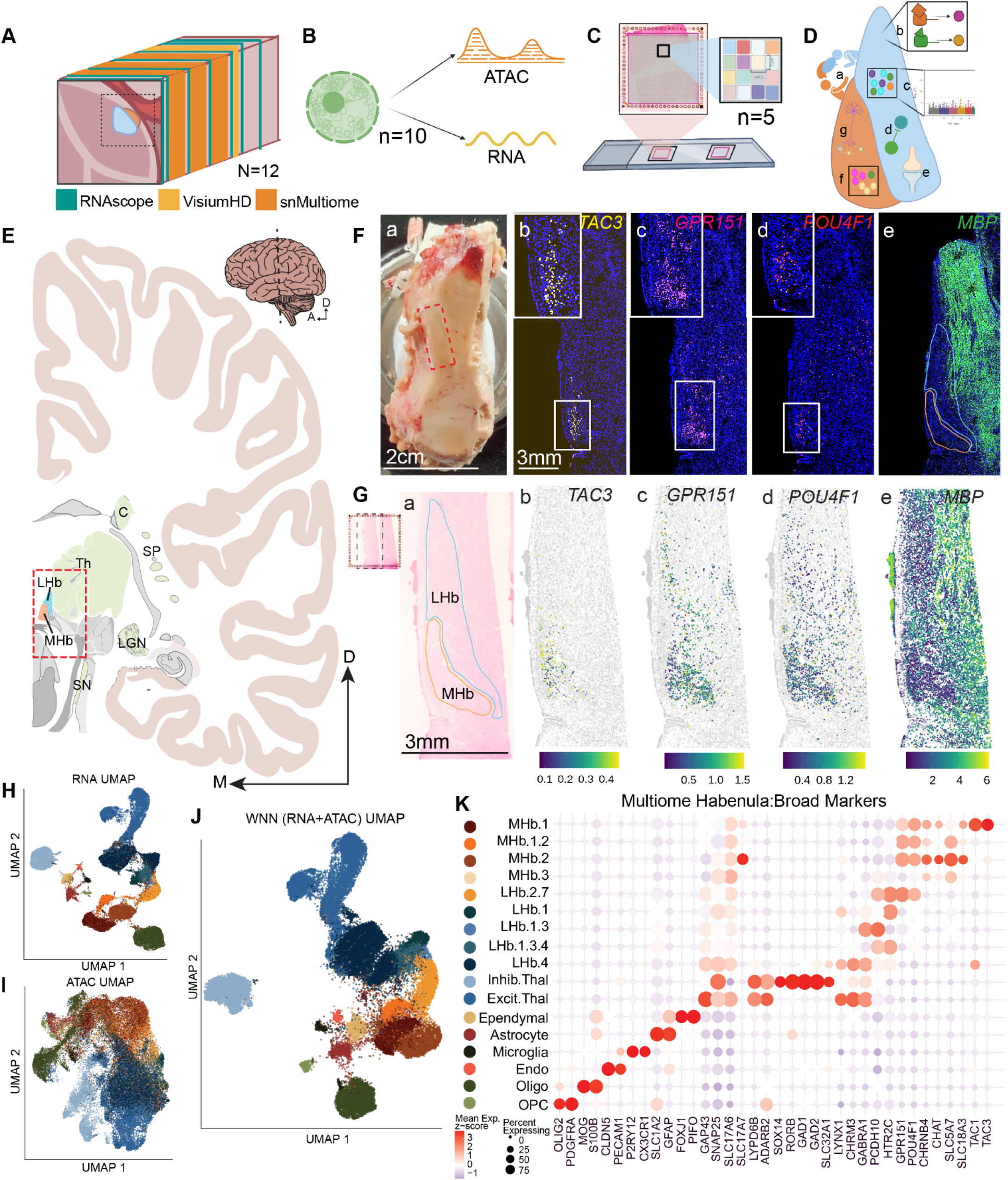
Multi-omic mapping of the human Hb. (**A**) Schematic of cutting scheme for smFISH/RNAscope, VisiumHD, and snMultiome. The Hb was scored to minimize thalamic contamination (n=12). (**B**) snMultiome schematic showing profiling of both the transcriptome and epigenome from the same nucleus (n=10 donors). (**C**) VisiumHD schematic showing subcellular spatial resolution (n=5 donors). (**D**) Visual representation of biological questions under investigation: **a.**) Cross talk between MHb and LHb. **b.**) Transcription factors regulating gene expression. **c.**) Cell types enriched in risk for disease states. **d.**) Identification of GABAergic neurons within the Hb. **e.**) Cell-to-cell communication between Hb neuron subtypes. **f.**) Composition and spatial organization of Hb cell types. **g.**) Neuron-glia interactions. (**E**) Coronal brain slab illustration depicting landmark structures used to guide Hb dissection (C - caudate, SP - segmented putamen, LGN - lateral geniculate nucleus, SN - substantial nigra, Th - thalamus). Red box demarcates approximate dissection of tissue blocks containing LHb and MHb. Medial (M), dorsal (D), and anterior (A) directions are shown by the respective arrows. (**F**) Anatomical validation of Hb dissections using RNAscope: **a.**) Tissue block with the Hb demarcated by red box. **b-e.**) Fluorescent images of Hb marker genes *TAC3* (MHb)*, GPR151* (LHb)*, POU4F1* (Hb), and *MBP* (WM). Insets are higher magnification images of the white boxes, and boundaries of the MHb and LHb are drawn in **e**. (**G**) VisiumHD from the same tissue block shown in E: **a.**) Small box to the left shows the entire 6x6 mm array containing two adjacent tissue sections, with the dashed line showing the representative section depicted at larger scale. Eosin staining completed during VisiumHD with boundaries of the MHb and LHb also in (F.e.). **b-e.**) Spatial expression of same marker genes in F. (**H**) UMAP of the gene expression/RNA data from snMultiome. (**I**) UMAP of the ATAC data from snMultiome. (**J**) Weighted Nearest Neighbors (WNN) UMAP of the integrated RNA and ATAC clusters. (**K**) Broad marker genes for the RNA WNN clusters, where the intensity of the color and size of the spot denote the expression of the respective gene.

From the snMultiome data, we recovered 54,193 high-quality nuclei across 10 donors following quality control (**Methods 4.4-4.7**, **Fig. S3**, **Fig. S4**, **Table S3**) with which we identified 40 fine-resolution clusters (**Fig. S5A**, **Methods 4.6**) using a Weighted Nearest Neighbor (WNN) graph incorporating both RNA and ATAC data. We mapped these clusters to our previous human epithalamic snRNA-seq dataset (Hb cell types were annotated as LHb.1-LHb.7 and MHb.1-MHb.3, **Methods 4.7**, **Fig. S5B-C**) (Yalcinbas et al. 2025) to establish initial cluster annotations. Following doublet filtering (**Fig. S5D-E**) and annotation refinement based on marker gene exploration (**Fig. S5F-G**), we retained 35 fine-resolution clusters. Because individual snMultiome clusters could correspond to more than one reference population from Yalcinbas et al., we retained these composite identities in the initial snMultiome cluster labels (e.g., LHb.2.7 for a snMultiome cluster mapping to both LHb.2 and LHb.7 in Yalcinbas et al.). We then consolidated fine-resolution clusters sharing the same cluster labels into 17 mid-resolution populations (**Fig. 1H-K**, **Fig. S5H**) representing distinct MHb and LHb populations as well as non-neuronal and thalamic populations (**Fig. 1K**).

Among mid-resolution clusters, MHb neurons separate into cholinergic populations defined by *CHAT*, *SLC5A7*, and *SLC18A3* (clusters MHb.1.2, MHb.2, MHb.3) and a Substance P population (MHb.1) marked by *TAC1*, encoding the precursor protein for Substance P (**Fig. 1K**). LHb neuronal populations express mouse LHb marker genes, including *CHRM3, GABRA1, PCDH10*, and *HTR2C* (Hashikawa et al. 2020; Wallace et al. 2020)(**Fig. 1K**). Intriguingly, LHb.4 expresses low levels of *SLC32A1* (VGAT) and *GAD1/2* in combination with glutamatergic markers, suggesting either a potential glutamate/GABA co-releasing phenotype or a putative GABAergic subcluster (**Fig. 1K**). Non-neuronal populations reflect expression profiles for canonical cell types, including OPCs (*OLIG2, PDGFRA*), oligodendrocytes (*MOG, S100B*), endothelial cells (*CLDN5, PECAM1*), microglia (*P2RY12, CX3CR1*), astrocytes (*SLC1A2, GFAP*), and ependymal cells (*FOXJ1, PIFO*) (**Fig. 1K**). We also identified excitatory (*SLC17A6*) and inhibitory (*SLC32A1*, *GAD1/2* ) neuronal populations expressing the thalamic markers *LYPD6B* and *ADARB2*, reflecting capture of adjacent thalamic tissue (**Fig. 1K**). Altogether, we identified 4 MHb and 5 LHb mid-resolution clusters, along with expected glial populations and neighboring thalamic neurons.

### 2.2 Conserved cross-species medial and lateral habenula cell types

Next, we sought to identify cross-species consensus Hb cell types to leverage evolutionary conservation as a framework for identifying stable Hb cellular identities (Zeng 2022; Shafer 2019; Arendt et al. 2016; Zhu et al. 2026). To derive consensus cell-type definitions across species, we first refined cluster annotations for Hb datasets in mouse (Hashikawa et al. 2020; Wallace et al. 2020) and zebrafish (Pandey et al. 2018) (**Fig. S6**, **Methods 4.8**). We then evaluated transcriptomic cluster replicability across all species using *MetaNeighbor* (Crow et al. 2018; Fischer et al. 2021)(**Fig. 2A**), where highly replicable clusters provide evidence for evolutionarily conserved cell types.

**Fig. 2:**
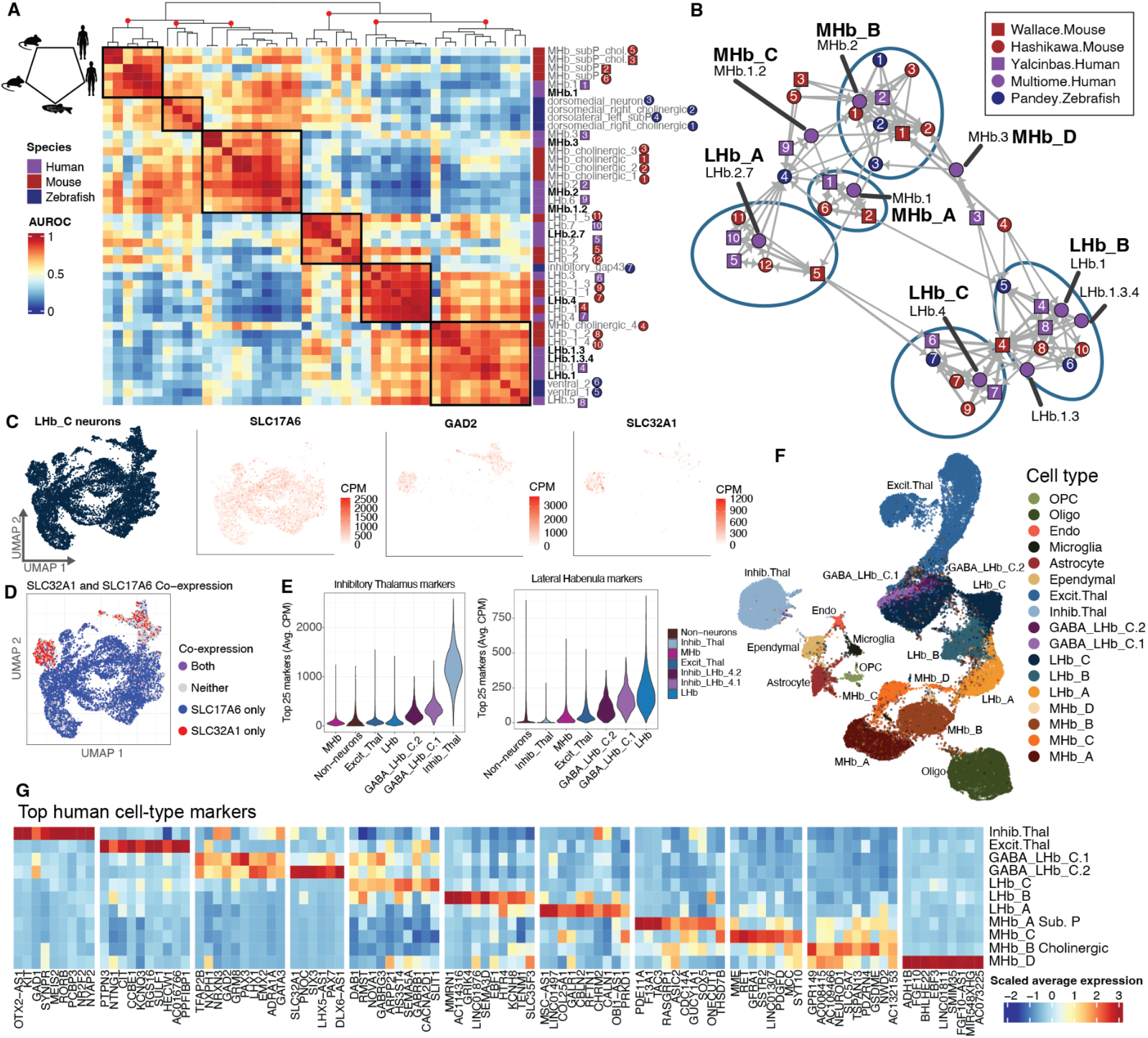
Consensus cross-species Hb cell types. (**A**) Hierarchically clustered heatmap of pairwise *MetaNeighbor* Area Under the Receiver Operating Characteristic curve (AUROC) scores, quantifying cross-cluster transcriptomic replicability across multiple studies (Hashikawa et al. 2020; Pandey et al. 2018; Wallace et al. 2020; Yalcinbas et al. 2025). Human snMultiome clusters are in bold. The numbered and colored shapes next to the cluster annotations correspond to (B). (**B**) Annotated cluster graph of BestVsNext *MetaNeighbor* AUROCs, quantifying strict per cluster matches. The numbered and colored shapes match the cluster annotations in (A). (**C**) UMAP of the snMultiome LHb_C neurons, colored by CPM expression of either *SLC17A6*, *GAD2*, or *SLC32A1*. (**D**) Same UMAP as in (C) with points colored based on detected transcripts of either *SLC17A6* (blue), *SLC32A1* (red), both (purple), or neither (light grey). (**E**) Violin plots of the average expression per cell of the top 25 inhibitory thalamus (left) or LHb (right) markers in CPMs. (**F**) UMAP of the human snMultiome RNA-assay depicting the final cross-species consensus cluster annotations used in all downstream analysis. (**G**) Top ten 1-vs-All markers per thalamus and Hb neuronal consensus cell type (F), with *Z*-scored average expression (normalized counts) per cell type. Duplicate marker genes were assigned to the cell type with the higher DE fold-change.

Broadly, the MHb and LHb are each characterized by three distinct cross-species cell-type groups (**Fig. 2A**). Within the MHb, the first cell-type group contains human and mouse Substance P populations, the second consists of zebrafish dorsal Hb clusters, and the third contains cholinergic populations from mouse and human (**Fig. 2A**). These findings support conserved MHb Substance P and cholinergic populations in mammals, while also capturing homology between zebrafish dorsal Hb and mammalian MHb despite greater transcriptional divergence between fish and mammals. Within the three LHb cell-type groups, one contains only human and mouse clusters and includes the snMultiome LHb.2.7 population. The two additional groups contain matches across all three species and separate snMultiome LHb.4 and LHb.1/1.3/1.3.4 populations, respectively (**Fig. 2A**). These analyses suggest that the LHb contains two transcriptionally distinct cell types that are conserved between zebrafish, rodents, and humans and one putative mammal-specific cell type.

To resolve one-to-one correspondence between individual clusters, we next applied the more stringent BestVsNext *MetaNeighbor* framework, which evaluates how well each reference cluster distinguishes its best-matching cluster from the next most similar population (**Fig. 2B**, **Methods 4.8**) (Crow et al. 2018; Fischer et al. 2021). This analysis largely reinforces the broader relationships identified with the base *MetaNeighbor* analysis (**Fig. 2A**), while clarifying several important annotations. Most notably, a conserved cholinergic group containing snMultiome MHb.2 now resolves across all three species (**Fig. 2B**). In contrast, snMultiome MHb.1.2 and MHb.3 show comparatively weak cross-species correspondence despite their broader clustering to cholinergic clusters (**Fig. 2A**), suggesting they may represent human-specialized cholinergic subtypes. The three LHb groupings identified in the base *MetaNeighbor* analysis (**Fig. 2A**) are preserved in this more stringent framework, with snMultiome LHb.1/1.3/1.3.4 exhibiting particularly strong mutual correspondence across datasets (**Fig. 2B**).

To place these cross-species relationships in the context of transcriptional variation within the human Hb, we next examined transcriptomic similarity exclusively among snMultiome clusters (**Fig. S7A**). Consistent with cross-species analyses, LHb.1, LHb.1.3, and LHb.1.3.4 form a cohesive transcriptional group, suggesting they represent a single conserved cell type partitioned into multiple fine-resolution clusters during unsupervised clustering. Conversely, MHb.1.2 occupies an intermediate transcriptional position between the conserved Substance P (MHb.1) and cholinergic (MHb.2) populations, while the small MHb.3 cluster (n=145 nuclei) remains transcriptionally distinct, indicating a potentially rare human cholinergic subtype.

Taken together, we establish a cross-species consensus annotation (**Fig. 2B**), where within MHb, MHb_A represents a conserved Substance P population (previously MHb.1), while MHb_B represents a conserved cholinergic population (previously MHb.2). MHb_C (previously MHb.1.2) and MHb_D (previously MHb.3) are transcriptionally distinct human cholinergic subtypes with comparatively weak cross-species correspondence. Within LHb, LHb_A represents the mammal-specific population (previously LHb2.7), whereas LHb_B combines the transcriptionally similar LHb.1, LHb.1.3, and LHb.1.3.4 clusters into a single consensus cell type, and LHb_C corresponds to the conserved cell-type group containing LHb.4. We provide mappings between all prior cluster labels and consensus cell-type annotations in non-human (**Table S4, Table S5, Table S6**) and human datasets (**Table S7**, **Table S8**). As an orthogonal assessment of these cross-species relationships, we quantified the aggregate expression of the top 50 marker genes from each human consensus cell type (**Results 2.4, Methods 4.8**) within individual mouse cells (**Fig. S8**). Human cell-type gene programs are enriched in the expected mouse populations, further supporting the transferability of consensus annotations at single-cell resolution. Notably, the mouse SubP-Chol populations (co-expressing cholinergic and Substance P markers (**Fig. S6**)) do not show preferential enrichment for the human MHb_C marker program, supporting MHb_C as a human-enriched cholinergic subtype. Altogether, we generate a robust cross-species consensus annotation of Hb cell types (**Fig. 2B**), which we employ for all subsequent analyses.

### 2.3 Identification of GABAergic neurons in human lateral habenula

The Hb is primarily composed of glutamatergic neurons, but some evidence suggests existence of GABAergic neurons. For example, rare populations of GABAergic neurons have been reported in Hb of mice (L. Zhang et al. 2018; Flanigan et al. 2020) and zebrafish (Pandey et al. 2018). However, functional studies in mice questioned whether these populations act as conventional inhibitory neurons, or rather co-release glutamate and GABA, while primarily driving excitatory responses (Quina et al. 2020; Green et al. 2023; Nguyen et al. 2024). To investigate the putative presence of these neurons in the human brain, we examined expression of genes required for the production and packaging of glutamate and GABA. Among all cross-species consensus Hb cell types, only LHb_C consistently expresses the canonical GABAergic markers *SLC32A1* (VGAT), *GAD1*, and *GAD2* (**Fig. S7B**). Within human snMultiome LHb_C, *SLC32A1* and *SLC17A6* (VGLUT2) are primarily expressed in separate nuclei, arguing against glutamate/GABA co-release from individual cells (**Fig. 2C-D**). Subclustering LHb_C reveals two transcriptionally distinct GABAergic subpopulations (**Fig. 2C**, **Fig. S9N**, **Methods 4.9**), which we termed GABA_LHb_C.1 (n=1,123 nuclei) and GABA_LHb_C.2 (n=932 nuclei). Given the proximity of the Hb to thalamic regions containing inhibitory neurons, we evaluated whether the LHb GABAergic neurons could be inhibitory thalamic neurons. GABA_LHb_C.1 and GABA_LHb_C.2 show strong expression of LHb gene markers and lower expression of inhibitory thalamic marker genes compared to the inhibitory thalamus cluster (**Fig. 2E**), supporting their identity as GABAergic LHb neurons. Furthermore, GABA_LHb_C.2 exhibits higher *SLC32A1* expression than GABA_LHb_C.1, suggesting GABA_LHb_C.2 as the more likely candidate for synaptic GABA release (**Fig. 2G**). As an orthogonal validation, RNAscope smFISH targeting *POU4F1*, *SLC32A1*, and *MBP* detects a sparse population of *SLC32A1*+ cells within the human Hb (∼0.4% of Hb cells), with signal localized predominantly in the LHb (**Fig. S9**, **Methods 4.10**). Together, these findings provide complementary evidence for a rare GABAergic neuronal population within the human LHb. We therefore retained these populations as distinct annotations for all subsequent snMultiome and SRT analyses (**Fig. 2F**).

### 2.4 Lateral and medial habenula subtypes exhibit unique molecular signatures

We next identified data-driven cell-type marker genes to establish unique molecular signatures for consensus Hb cell types (top 10 markers in **Fig. 2G**, top 50 markers in **Table S9**, **Methods 4.8 and 4.11**). In the LHb, LHb_C is distinguished by prominent expression of *SEMA5A*, whereas LHb_B is characterized by *SEMA3D* expression. Both genes are semaphorin (SEMA) family members, which have established roles in Hb axon fasciculation (Roberson and Halpern 2018), suggesting distinct molecular programs governing axonal connectivity. LHb_B is also characterized by expression of *GRIK4*, a kainate receptor subunit associated with treatment-resistant depression (Paddock et al. 2007; MILANESI et al. 2015). LHb_A preferentially expresses the inhibitory G-protein-coupled receptors *GALR1* and *CHRM2*, previously implicated in antidepressant responsiveness (García-Durán et al. 2021) and drug-seeking behavior (Wolfe et al. 2022).

MHb subpopulations show similarly specialized molecular signatures. The MHb_A Substance P population expresses *PDE11A* and *GUCY1A1*, enzymes with complementary roles in nitric oxide-cGMP signaling, a pathway that regulates Substance P neuron activity in mice (Ciscato et al. 2025). The acid-sensing ion channel *ASIC2* is broadly enriched across multiple MHb populations (MHb_A/B/D), suggesting acid sensing may represent a shared feature of MHb neuron physiology. Cholinergic populations (MHb_B/C/D) broadly share MHb_B markers, in addition to more distinct molecular specializations. For example, MHb_B shows high expression of *GSDME*, a pore-forming protein implicated in inflammation and cell death that has been linked to mitochondrial damage and neurodegeneration in neurons (Neel et al. 2023). MHb_C is distinguished by *MME*, encoding an enzyme that degrades neuropeptides, including Substance P. Finally, *MIR548XHG* is among the top markers of the rare MHb_D population and it is also highly expressed in human MHb.3 in the Yalcinbas et al. reference dataset (Yalcinbas et al. 2025). *MIR548XHG* is the host gene for the primate-specific miR-548 family (Liang et al. 2012), providing additional molecular support for the putative human specialization of the MHb_D cholinergic subtype. Together, data-driven marker analysis reveals diverse molecular programs across consensus human Hb cell types, highlighting candidate pathways regulating neuronal connectivity, intrinsic physiology, and neuromodulatory signaling.

### 2.5 Molecular evidence for functional specialization of human habenula cell types

To infer putative functional specializations of LHb and MHb subpopulations, we identified Gene Ontology (GO) terms that were both enriched among the top marker genes of Hb cell types and also associated with cell-type identity in a complementary *MetaNeighbor* analysis (**Fig. S7E**, **Methods 4.12**). Of the eight GO terms supported by both analyses, six were related to glutamatergic/GABAergic signaling or synapse assembly (**Fig. S7E**), highlighting neurotransmission as a prominent axis of molecular specialization. To further understand the heterogeneity across Hb cell types, we characterized the gene expression profiles of these enriched GO terms.

Distinct electrophysiological profiles have been observed within the LHb, including tonic and burst firing, which is altered in models relevant for neuropsychiatric disease (Li et al. 2011; Yang, Wang, et al. 2018; Wagner et al. 2017). Consistent with previous physiological studies, the human LHb is enriched in genes implicated in burst firing, including those encoding NMDA receptor subunits and T-type voltage-sensitive calcium channels (T-VSCCs) (Yang, Cui, et al. 2018) (**Fig. 3A-C**). However, receptor composition differs substantially between LHb cell types. While *GRIN2B* is broadly expressed throughout the Hb, *GRIN1*, *GRIN2D*, and *GRIN3A* are enriched in all three LHb_C cell types, whereas *GRIN2A* is predominantly expressed in LHb_A and LHb_B. Likewise, the T-VSCC genes *CACNA1G* and *CACNA1I* are preferentially LHb-enriched (**Fig. 3C**). Tonic firing in the LHb has been linked to hyperpolarization-activated cyclic nucleotide-gated (HCN) channels (Poller et al. 2011; Notomi and Shigemoto 2004). Correspondingly, *HCN1*, *HCN2*, and *HCN3* are enriched across LHb cell types, with all three highest in GABA_LHb_C.1 (**Fig. 3D**). *HCN1* shows the greatest cell-type specificity, with high expression in GABA_LHb_C.1 and LHb_B but low expression in LHb_A. (**Fig. 3D**). Together, these results indicate that human LHb neuronal subtypes possess distinct combinations of receptors that likely influence their electrophysiological properties.

**Fig. 3:**
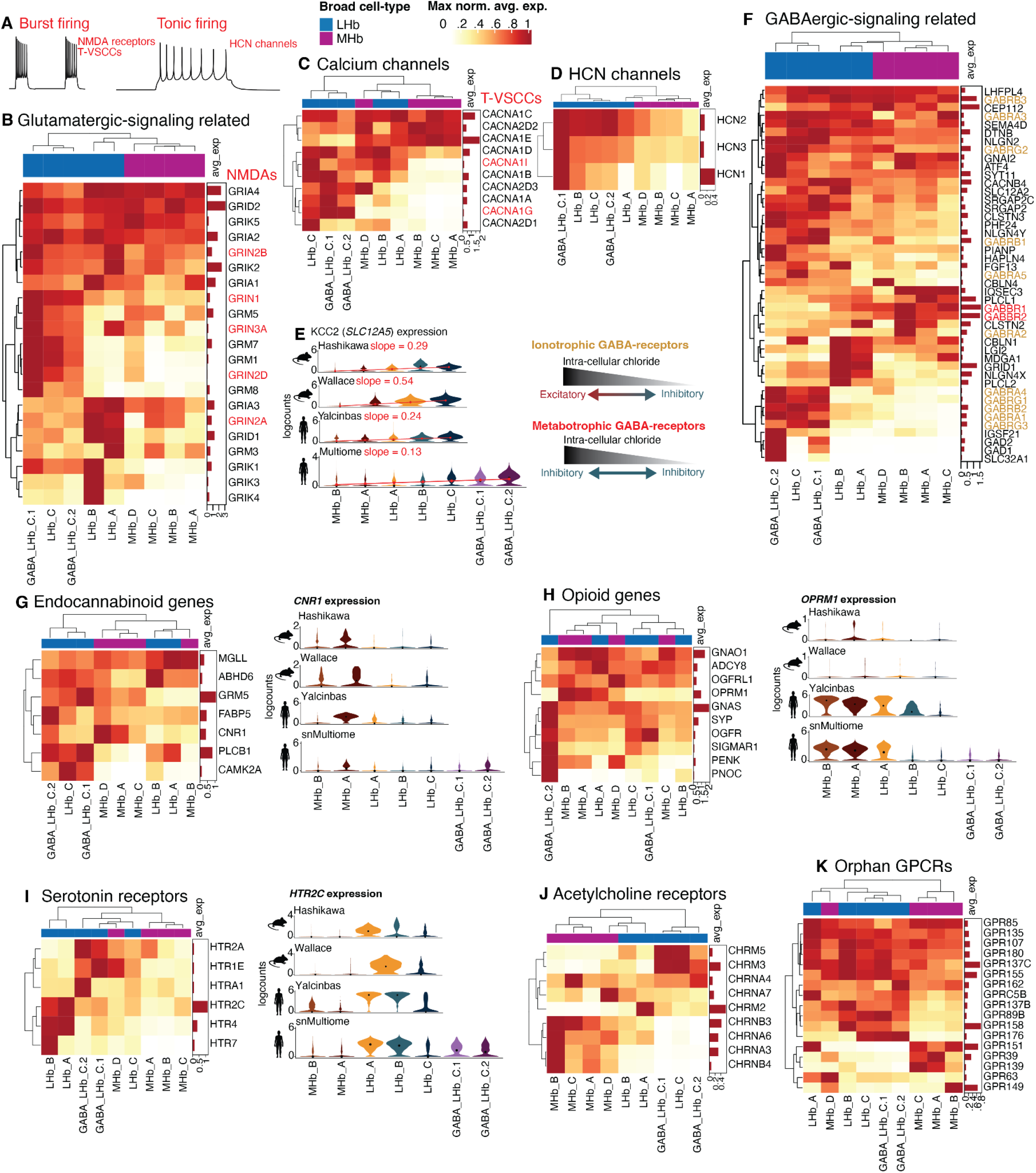
Transcriptional landscape of human Hb cell types. (**A**) Schematics of burst and tonic firing profiles. (**B**, **C**, **D**, **F**, **G**, **H**, **I**, **J**, **K**) Hierarchically clustered heatmaps of gene expression across Hb cell types. For each heatmap, gene expression (logcounts) is averaged across cells per cluster (columns) and then max normalized across cell types (rows). Columns are annotated by broad cell type: LHb in blue and MHb in purple. Genes shown are: (**B**) Genes from glutamatergic-signaling GO terms derived in **Fig. S7E**, (**C**) calcium channels, (**D**) HCN channels, (**F**) Genes from GABAergic-signaling GO terms derived in **Fig. S7E**, (**G**) endocannabinoid genes sourced from GO, (**H**) opioid genes sourced from GO, (**I**) serotonin receptors, (**J**) acetylcholine receptors, or (**K**) orphan GPCRs. (**E**) Violin plots of *SLC12A5* expression ordered by median expression across cell types for the Hashikawa et al., Wallace et al., Yalcinbas et al., and snMultiome consensus Hb cell types. A fitted linear regression line is shown in red with the slope reported for each dataset (all p-values < 2.2e-16). The adjacent schematic highlights the excitatory vs inhibitory response of activated ionotrophic vs. metabotrophic GABA-receptors depending on intra-cellular chloride concentration. (**G**, **H**, **I**) Violin plots for *CNR1*, *OPRM1*, and *HTR2C*, respectively.

The Hb is also a locus of non-canonical GABAergic signaling; reduced expression of the chloride transporter KCC2 (*SLC12A5*) in rodent Hb can produce depolarizing (excitatory) responses to GABA through elevated intracellular chloride concentrations (Wagner et al. 2016; U. Kim and Chung 2007). *SLC12A5* follows a conserved expression gradient across cell types in both mouse and human (**Fig. 3E**), with the lowest expression observed in cholinergic MHb neurons and progressively higher expression across LHb_A, LHb_B, and LHb_C. This conserved pattern suggests that non-canonical GABAergic signaling is likely most pronounced within cholinergic MHb neurons, while also predicting graded differences in GABA responsiveness across LHb cell types. Intriguingly, the metabotropic GABA receptors *GABBR1* and *GABBR2*, which inhibit neuronal activity through G protein-coupled signaling independent of chloride gradients, are broadly elevated in expression in MHb compared to LHb, with the cholinergic MHb_B cell type showing highest expression (**Fig. 3F**). Together, these findings implicate the cholinergic MHb as uniquely specialized for integrating GABA-signaling through both chloride-dependent ionotropic and chloride-independent metabotropic mechanisms.

Beyond glutamatergic and GABAergic signaling, the Hb integrates diverse neuromodulatory inputs involved in reward processing, aversion, and motivated behavior (Namboodiri et al. 2016). To better understand which cell types integrate these neuromodulatory signals, we mapped genes associated with endocannabinoid, opioid, serotonergic, and cholinergic signaling across cell types (**Fig. 3G-J**). The endocannabinoid receptor CB1 (*CNR1*) is enriched within GABA_LHb_C.2, MHb_A Substance P, and MHb_D populations, suggesting existence of distinct cannabinoid-sensitive LHb and MHb cell types. Conserved *CNR1* expression between humans and mice is particularly apparent for Substance P neurons (**Fig. 3G**). Consistent with *OPRM1* expression in rodent Hb (Popat et al. 2026), *OPRM1* is enriched in the human MHb, and the mammal-specific LHb_A population. Notably, the human Hb expresses *OPRM1* at much higher levels than mice in our sampled datasets (**Fig. 3H**). Genes encoding serotonergic and cholinergic receptors are selectively expressed across Hb cell types with *HTR2C* and *HTR4* enriched in LHb_A and LHb_B, and *HTR7* largely restricted to LHb_B. *HTR2C*, the most highly expressed serotonergic receptor in the human Hb, also displays a highly conserved cell-type expression profile in the mouse Hb (**Fig. 3I**). As expected, expression of nicotinic acetylcholine receptor subunits is concentrated almost exclusively within the cholinergic MHb, particularly in MHb_B (**Fig. 3J**). Among these, *CHRNA3* and *CHRNB4* are especially enriched. Given the well-established genetic association between the *CHRNA3-CHRNA5-CHRNB4* locus and nicotine dependence (Improgo et al. 2010), these results support cholinergic MHb neurons as the principal cellular substrate through which this genetic risk locus influences human Hb function.

G-protein-coupled receptors (GPCRs) are pharmacologically accessible modulators of neuronal function (Sriram and Insel 2018; Civelli 2012) and many orphan GPCRs, which often lack identified endogenous ligands, have been identified in the rodent Hb and implicated in metabolic, neuropsychiatric, and substance-use phenotypes (Roy and Parhar 2022). In the human Hb, *GPR149* is enriched in MHb_B cholinergic neurons, whereas *GPR158* is LHb-enriched (**Fig. 3K**); both have been implicated in energy metabolism (Wyler et al. 2024; J. Lin et al. 2022), with *GPR158* additionally functioning as a glycine receptor that can modulate neuronal excitability (Laboute et al. 2023). *GPR176*, a constitutively active GPCR implicated in GABAergic activity regulation (Tian et al. 2025), is LHb_C-enriched. Consistent with rodents, *GPR151* and *GPR139*, implicated in nicotine intake (Antolin-Fontes et al. 2020) and compulsive alcohol consumption (Kononoff et al. 2018), respectively, show MHb expression. Altogether, these analyses reveal extensive cell type-specific specialization of functionally relevant neurotransmitter receptors, ion channels, and neuromodulatory pathways across the human Hb, defining molecular features that may shape the distinct physiological properties and signaling responses of individual Hb neuronal subtypes.

### 2.6 Disease-associated genes show cell type-specific regulation in the human habenula

To better understand the regulatory mechanisms underlying Hb cell type-specific transcriptional specialization, we leveraged the paired RNA-seq and ATAC-seq data from snMultiome. Using *TRIPOD* (Jiang et al. 2022), we integrated chromatin accessibility and gene expression to identify regulatory trios linking transcription factors (TFs), cCREs, and their putative target genes (**Fig. 4A**, **Fig. S10**, **Table S10**, **Methods 4.13-4.14**). *TRIPOD* identifies conditional associations among TF expression, accessibility of a peak containing the TF motif, and expression of putative target genes (**Fig. 4A**). Level 1 trios are supported by conditional associations among these features, whereas Level 2 trios additionally exhibit interaction effects in which the strength of the peak-gene relationship depends on TF expression, signifying potentially stronger regulatory relationships (Jiang et al. 2022). Across Hb cell types, *TRIPOD* identified thousands to tens of thousands of Level 1 and Level 2 trios (**Fig. S11A-B**). Level 1 and Level 2 trios show similar distributions around transcription start sites (TSSs), but Level 2 trios exhibit greater density than Level 1 trios at distances of 100-500 kb from TSSs, suggesting an increased contribution of distal regulatory elements to these stronger regulatory relationships (**Fig. S11C**).

**Fig. 4:**
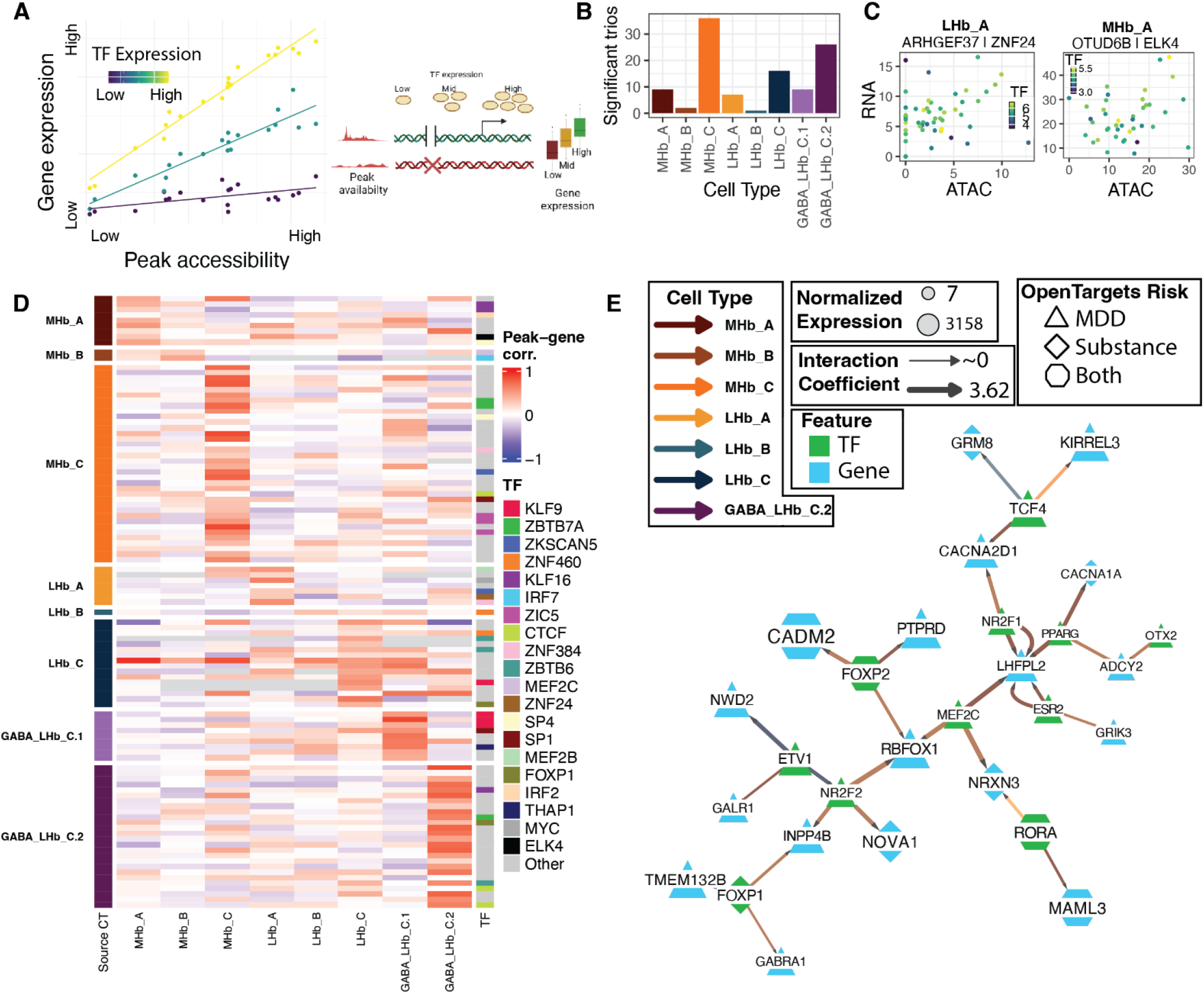
Identification of Hb cell-type-specific gene regulatory networks associated with MDD and addiction. (**A**) Cartoon with example data demonstrating the mathematical and biological properties of trios. When transcription factor (TF) expression is held constant, peak accessibility and gene expression are correlated (“level 1 trios”). As TF expression increases, higher peak accessibility induces larger increases in gene expression (“level 2 trios”). (**B**) Number of significant intersecting trios (i.e. both level 1 and level 2) for Hb neuronal cell types. (**C**) Normalized gene expression plotted against normalized accessibility for an intersection trio found in LHb_A, with peak chr5-149217776-149218031, and MHb_A, with peak chr8-91040035-91041213, respectively. Points are metacells, colored by log-normalized TF expression. (**D**) Correlation across each cell type’s metacells between gene expression and peak accessibility plotted for the top-20 intersection trios per Hb cell type. Each row is a trio, and 20 of the most common TFscolor these rows. Row groups indicate the cell type where the trio was found, while columns indicate the cell type in which expression and accessibility are measured. (**E**) Subsets of the level 2 trio networks (**Table S10**) among Hb cell types and *OpenTargets* MDD or SUD-associated genes (**Methods 4.26**). TFs and regulated genes are shown as green or blue nodes, respectively. Node size is determined by the gene expression (logcounts). Edge arrows show which genes are regulated by each TF. Edge width scales with the *TRIPOD* level 2 coefficient, indicating the degree to which TF expression interacts with peak accessibility to produce increases in gene expression. Edge color represents the cell type. Genes overlapping *OpenTargets* MDD or SUD-associated genes are shown as triangle or diamond nodes, respectively. Genes overlapping both MDD and SUD lists are shown as hexagons.

To focus on the most stringent regulatory relationships, we retained the intersection of Level 1 and Level 2 trios, yielding tens of trios per Hb cell type (**Fig. 4B**). For example, we identified the TF *ZNF24* regulating *ARHGEF37* (guanyl-nucleotide-exchange factor involved in endocytosis) in LHb_A and *ELK4* regulating *OTUD6B* (protein deubiquitinase) in MHb_A. (**Fig. 4C**). Trios were highly cell type-specific, with individual trios exhibiting strong peak-gene correlations in the cell type in which they were identified and substantially weaker correlations across other cell types (**Fig. 4D**, **Fig. S11B**).

Finally, we used *RCy3* (Gustavsen et al. 2019; Shannon et al. 2003) to predict networks among TFs and their putative target genes (**Fig. 4E, Methods 4.15**). Using the significant level 2 trios (**Table S10**), we built graphical networks differentiating TFs from their target genes with cell type-specific edges. We intersected these networks with genes implicated in MDD and SUD as defined by *OpenTargets* (Buniello et al. 2025), a database of candidate drug target genes for multiple diseases curated through gene-disease associations, GWAS, and other publicly-available data. Given the putative role of the Hb in depression and addiction, we filtered to TFs and putative targets associated with MDD or SUD and refined network complexity to focus on cell type-specific regulation associated with disease. As one example, despite relatively low expression in MHb_A and MHb_B (**Fig. 3**), calcium channel genes *CACNA1A* and *CACNA2D1* are specifically regulated in these cell types (**Fig. 4E**), highlighting MHb-specific pathways related to calcium signaling associated with MDD and SUD. *LHFPL2* shows the greatest convergence of putative regulatory relationships among disease-associated target genes, with multiple TFs (*MEF2C*, *ESR2*, *NR2F1*, *PPARG*) linked to the gene specifically in MHb_A Substance P neurons. For two of these TFs, independent trios linked the same TF to *LHFPL2* through two distinct accessible chromatin regions, providing multiple putative regulatory connections to the same target. Together, these analyses identify candidate TF-cCRE-gene relationships implicated in disease and reveal cell type-specific regulatory architecture across human Hb cell types.

### 2.7 Distinct neuro-glial microenvironments in the medial and lateral habenula

Immunohistochemical studies determined that the human Hb is organized into discrete MHb and LHb subregions containing distinct cell populations (Marburg 1944; Díaz et al. 2011). To characterize the spatiomolecular anatomy of MHb and LHb subregions, we initially performed spot-based SRT (Visium) across 4 different Hb A-P levels in 3 neurotypical donors (**Fig. S12**, **Methods 4.16-4.18**). Due to the Hb’s small size and the limited resolution of the Visium platform, we could not resolve LHb and MHb. To resolve the anatomical organization and local microenvironments of LHb and MHb cell types, we also generated SRT data with the VisiumHD platform in 5 neurotypical control donors, which provides transcriptome-wide gene expression at subcellular spatial resolution (**Methods 4.19**, **Fig. 1**).

Following VisiumHD quality control (**Fig. S13**) and nuclei segmentation (**Fig. S14**), transcriptomic features were assigned to cells (**Methods 4.20-23**). Cells were then clustered using *BANKSY* (Singhal et al. 2024) and registered to snMultiome cell types to annotate spatially-defined VisiumHD cell types (**Fig. 5A**, **Fig. S15**, **Fig. S16**, **Methods 4.20-4.23)**. Given that assigning transcriptomic features to segmented cells utilizes only ∼20% of VisiumHD bins, we also leveraged *FICTURE* (Si et al. 2024), an algorithm which clusters agnostic to cell segmentation and uses both cellular and extracellular bins (**Fig. S17**). All *BANKSY* and *FICTURE* clusters were represented in all donors. Donor-biased *BANKSY* clusters were restricted to thalamus and endothelial clusters, likely reflecting variable sampling of thalamic tissue and vasculature across sections (**Fig. S18**). *FICTURE* identified glia-rich white matter tracts, but did not capture distinct Hb cell types (**Fig. S19**). Therefore, to analyze cell type-specific interactions, we moved forward with *BANKSY* clusters for downstream analyses, which we refer to as VisiumHD clusters (**Fig. 5B-C**).

**Fig. 5:**
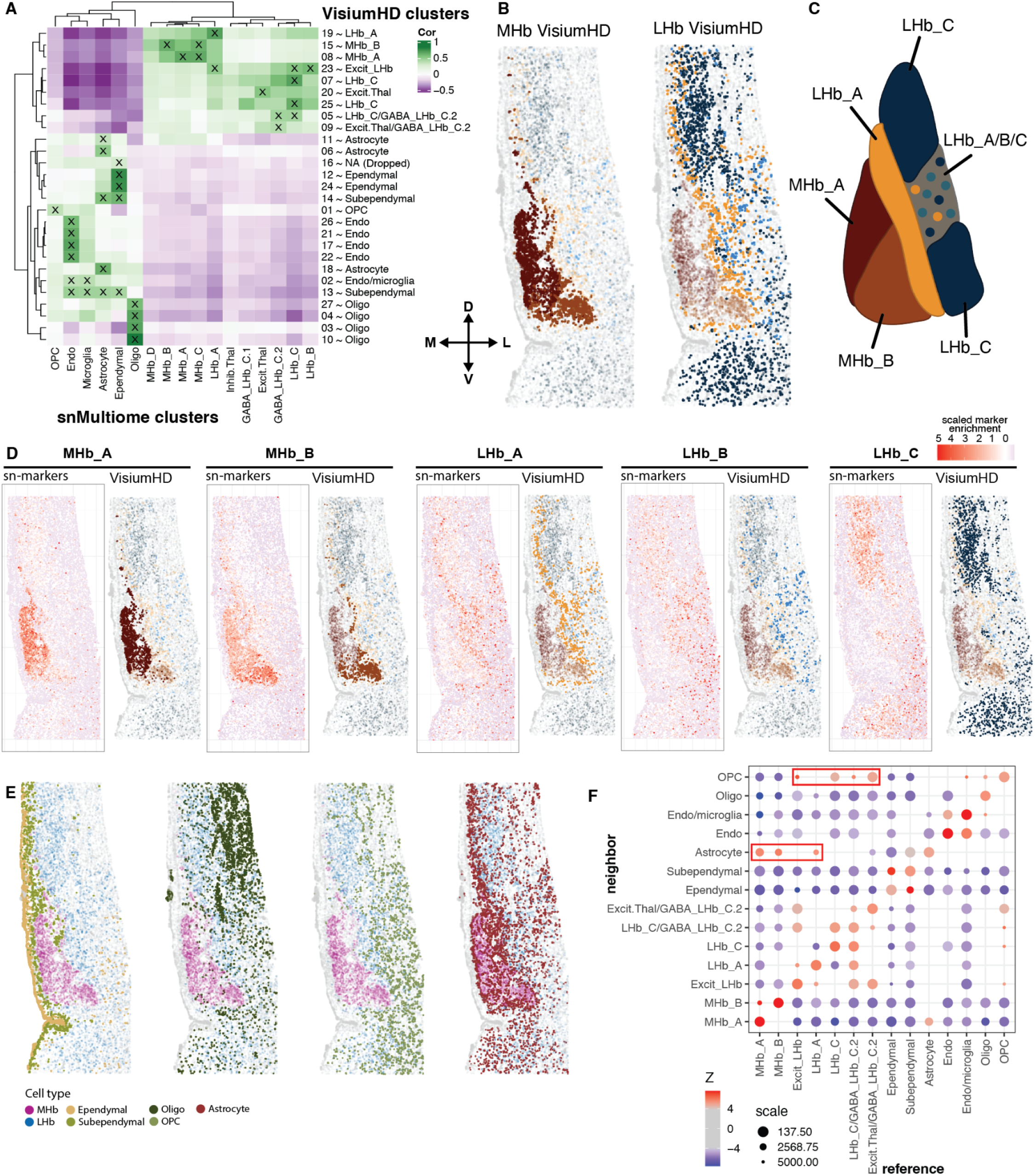
Topographical organization of molecularly-defined MHb and LHb cell types. (**A**) Transcriptional cluster registration between Hb snMultiome clusters (columns) and *BANKSY* spatial VisiumHD clusters (rows). Correlations are computed across transcriptome-wide 1-vs-Alll DE *t*-statistics to identify transcriptionally similar clusters, with the ‘x’ representing high confidence matches (**Methods 4.20**). (**B**) Representative donor (Br9090) showing the spatial organization of the annotated VisiumHD Hb clusters, for MHb (left) and LHb (right) VisiumHD clusters collapsed by consensus Hb cell type (**Fig. S16**, **Methods 4.20**). Arrows denote the anatomical orientation of VisiumHD sections: medial (M), lateral (L), dorsal (D), ventral (V). (**C**) Schematic of MHb and LHb cell type spatial organization based on B.(**D**) Spatial plots for Br9090 comparing aggregate snMultiome marker expression (left) to the associated VisiumHD cluster(s) (right) for the consensus Hb cell types. (**E**) Spatial plots for Br9090 highlighting non-neuronal VisiumHD clusters with MHb and LHb neuronal subpopulations grouped together. Left to right: ependymal and sub-ependymal, oligodendrocytes, OPCs, and astrocytes. (**F**) Bubble heatmap depicting *CRAWDAD* quantification of significant spatial cell-type colocalization (**Methods 4.21**). Red boxes highlight Astrocyte and OPC colocalization as neighbors to MHb_A/B/LHb_A and Excit_LHb/LHb_C/GABA_LHb_C.1/2, respectively.

To validate VisiumHD cell-type annotations, we examined the spatial expression of snMultiome cell-type marker profiles within VisiumHD tissue sections (**Fig. 5D**, **Methods 4.8**). Across neuronal populations, aggregate expression of snMultiome cell-type profiles closely recapitulates the boundaries of their corresponding VisiumHD clusters, strongly supporting VisiumHD cell-type annotations (**Fig. 5B-C**). In MHb, MHb_A Substance P neurons are located dorsomedially and MHb_B cholinergic neurons are located ventrolaterally, although the precise boundary shifts across donors, likely reflecting differences in AP sampling or coronal plane (**Fig. S20**, **Fig. S21**). In LHb, there is greater spatial heterogeneity, with LHb_A forming a narrow domain adjacent to the MHb and excitatory LHb_C occupying distinct dorsal and ventral territories. Although a single VisiumHD cluster mapped to snMultiome LHb_A, LHb_B, and LHb_C (VisiumHD Excit_LHb, **Fig. 5A**), marker projection reveals clear enrichment of the LHb_B gene program within the Excit-LHb spatial cluster, located in the medial portion of the LHb (**Fig. 5C)**. MHb_C, MHb_D, and GABAergic LHb_C populations were not resolved as distinct spatial clusters in VisiumHD. MHb_C marker expression identifies a localized population in only one donor, likely reflecting AP sampling variability (**Fig. S22**), whereas MHb_D lacks discrete spatial organization (**Fig. S23**), consistent with its rarity in the snMultiome data. GABAergic LHb_C populations similarly lack restricted spatial positioning and largely recapitulate the broader spatial organization of LHb_C (**Fig. S24**).

Among non-neuronal populations, we find similarly strong correspondence between snMultiome and VisiumHD cell types, with each major non-neuronal snMultiome cell type represented by at least one uniquely matched VisiumHD cluster with the exception of microglia (**Fig. 5A**). Ependymal and subependymal clusters line the medial border of the Hb adjacent to the third ventricle, whereas oligodendrocyte and OPC clusters define white matter tracts along the LHb that separate the Hb from adjacent thalamic regions (**Fig. 5E**, **Fig. S25**). LHb neurons are intermingled with these white matter domains, suggestive of a gradual transition between the LHb and surrounding thalamus.

Intriguingly, the grouped VisiumHD astrocyte clusters show a striking enrichment within regions containing MHb and LHb_A (**Fig. 5E**). Although astrocytes have been implicated in regulating LHb activity (Y. Cui et al. 2018; Xin et al. 2025), their preferential localization to the MHb suggests that they may also contribute to the distinct cellular environment and regulation of MHb neurons. Surprisingly, aggregate expression of astrocyte marker genes from snMultiome appears depleted in the MHb of VisiumHD samples, (**Fig. S26**), despite the presence of a clearly annotated fine resolution VisiumHD astrocyte cluster (*BANKSY* 11) within the MHb (**Fig. S26**). This pattern likely reflects image processing challenges in the densely packed MHb, where astrocyte segmentation may capture transcripts from adjacent neurons, impacting accurate marker gene projection. Indeed, this MHb-specific astrocyte cluster shows poor transcriptional correspondence to the snMultiome astrocyte cluster when using cellular-segmented transcripts, but correspondence increases when restricted to nuclear-segmented transcripts (**Fig. S26**). Nevertheless, direct expression of canonical astrocyte marker genes supports the presence of astrocytes within the MHb (**Fig. S26**).

To quantitatively characterize these spatial relationships across all VisiumHD samples, we applied *CRAWDAD* (dos Santos Peixoto et al. 2025), which tests cell-type colocalization and separation across spatial distances (**Methods 4.24-4.25**). Consistent with anatomical observations, *CRAWDAD* identifies significant spatial associations between astrocytes, MHb cell types, and LHb_A. Consistent with white matter tracts at the lateral edge of the LHb, CRAWDAD found spatial associations between OPCs and LHb_C cell types, including the GABAergic LHb_C populations (**Fig. 5F**). Together, these results reveal distinct non-neuronal microenvironments across the Hb, with the MHb enriched for astrocytes and adjacent to ependymal/subependymal compartments lining the third ventricle, and the LHb intermingled with white matter tracts at the border of the thalamus.

### 2.8 Cell-cell communication reveals specialized molecular environments across the MHb and LHb

Although the MHb and LHb are anatomically adjacent, the extent to which these domains communicate through local molecular signaling remains unclear. We therefore used the cell-cell communication (CCC) tool *LIANA+* (Dimitrov et al. 2024) to infer spatially informed ligand–receptor (LR) interactions between Hb cell types (**Methods 4.26**). Across cell-type pairs, MHb_A Substance P and MHb_B cholinergic neurons show higher LR communication with one another compared to LR communication between LHb cell types, suggesting more extensive local signaling within the MHb (**Fig. 6A**). Strong LR communication is also notable between MHb_A Substance P neurons and ependymal/sub-ependymal cells (**Fig. 6A**). While portions of the LHb are also adjacent to the ependymal/sub-ependymal layer, LR communication is much weaker among these populations, suggesting stronger signaling relationships between the MHb and ventricle wall compared to LHb (**Fig. 6A**). Consistent with the spatial relationships identified by *CRAWDAD* (**Fig. 5E**), astrocytes show stronger LR communication with MHb than LHb cell types, whereas oligodendrocytes show stronger communication with LHb than MHb cell types. We next asked whether the inferred LR interactions were enriched for disease-associated genes curated by *OpenTargets* (Buniello et al. 2025). Excit_LHb and LHb_C LR-pairs account for the majority of neuronal enrichment for MDD-associated genes, while MHb_A LR-pairs are enriched for genes associated with substance dependence (**Fig. 6B-C**).

**Fig. 6:**
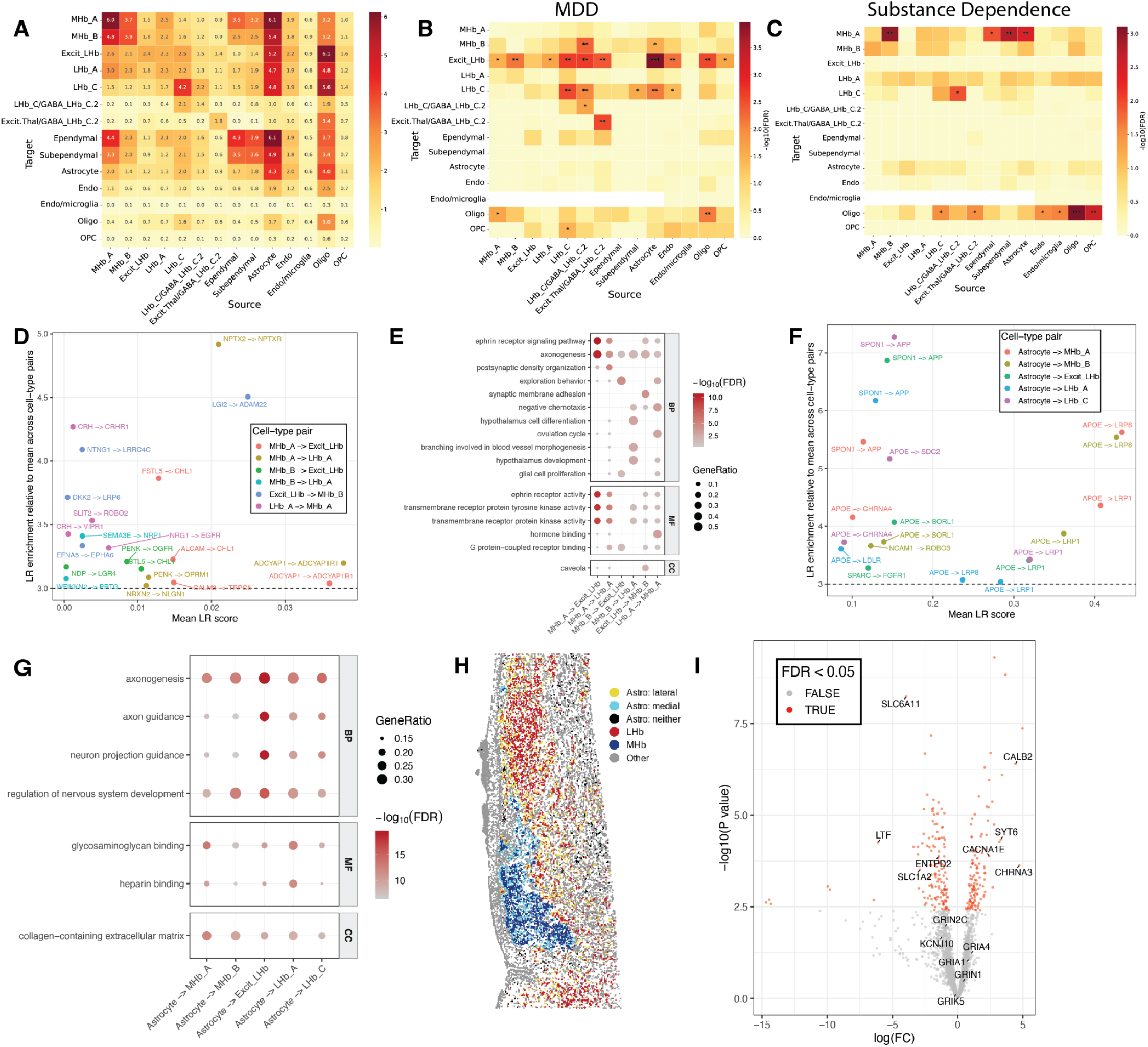
Spatially-informed cell-to-cell communication reveals distinct signaling relationships between LHb and MHb neurons and glial populations. (**A**) Heatmap showing the mean Ligand-Receptor (L-R) spatially-informed interaction global inflow score for significant (FDR < 0.05) L-R pairs computed with *LIANA+* across unique source-target (column and rows, respectively) cell type pairs (**Methods 4.26**). One-sided Fisher’s exact test enrichment heatmap for *OpenTargets* (Buniello et al. 2025) genes associated with (**B**) MDD or (**C**) SUD across ligands or receptors involved in L-R across source-targets pairs in (A). Color scales indicate −log_10_(p). ‘*’ denotes FDR < 0.05, ‘**” FDR < 0.01, and ‘***’ FDR < 0.001. (**D**) Interacting Hb neuron top L-R pairs by mean LR score. *Y*-axis shows the relative enrichment for each L-R pair in a specific source-target cell type pair compared to the mean score across all 196 (14 x 14) source-target cell type pairs for that L-R. A given L-R pair is shown up to 3 times across source-target cell pairs based on its mean L-R score. (**E**) Overrepresented GO terms from significant MHb to LHb L-R genes across specific interacting Hb neuronsource-target cell type pairs (**Methods 4.26**). (**F**) Similar to (D), but for the top astrocyte to Hb L-R pairs. (**G**). Similar to (E), but for astrocyte to Hb L-R genes. (**H**) Spatial plot of Br9090_1 depicting how astrocytes were labeled for differential expression (DE) (**Methods 4.27**). (**I**) DE volcano plot for LHb astrocytes against MHb astrocytes, with positive fold changes indicating higher expression in MHb (**Methods 4.27**). Genes are colored red if significant by adjusted p-value and spatial empirical adjusted p-value, and colored gray otherwise.

To further characterize potential molecular crosstalk between MHb and LHb, we focused on cell-type-specific LR interactions between neuronal populations in these subregions (**Fig. 6D**), which highlighted interactions involving LHb_A and MHb given their close spatial relationship (**Fig. S27**). We identified *PENK-OPRM1* signaling between *PENK*-expressing MHb_A and *OPRM1*-expressing LHb_A, suggesting a potential route for endogenous opioid signaling across the MHb-LHb boundary. Consistent with a potential role for the Hb in stress-related neuropsychiatric disorders (Jacinto et al. 2017; Baker et al. 2022), we identified interactions involving *CRH*-expressing LHb_A and its receptors *VIPR1* and *CRHR1* expressed by MHb_A. LHb_A also expresses the axon guidance molecule *SLIT2*, whereas MHb_A expresses its receptor *ROBO2*, suggesting potential *SLIT-ROBO* signaling across the LHb-MHb boundary. In addition, we identified contact-dependent Eph/Ephrin signaling between *EFNA5*-expressing Excit-LHb and *EPHA6*-expressing MHb_B, further supporting putative local signaling between neuronal populations at the LHb-MHb border. More broadly, GO enrichment across MHb-LHb LR interactions identified significant enrichment for ephrin signaling, GPCR binding, peptide receptor, and synaptic terms (**Fig. 6E**).

Finally, given that astrocytes play a critical role in regulating LHb firing dynamics in the rodent Hb (Y. Cui et al. 2018; Xin et al. 2025), we examined neuron-astrocyte signaling in the human Hb by comparing LR communication between astrocytes and all Hb neuronal populations. Cell-type-specific astrocyte-neuron interactions were largely dominated by the astrocyte-derived ligands *APOE* and *SPON1* (**Fig. 6F**), while broader astrocyte-neuron interactions were significantly enriched for axonogenesis-related and extracellular matrix-related GO terms across cell-type combinations (**Fig. 6G, Table S11**). Given evidence for differential astrocyte-mediated regulation of MHb and LHb neurons in rodents (Gampe et al. 2012; Y. Cui et al. 2018; Xin et al. 2025), we leveraged the high-resolution VisiumHD data to isolate MHb- and LHb-associated astrocytes and compared their transcriptional profiles (**Fig. 6H, Methods 4.27**). *ENTPD2*, a marker of MHb-specific non-stellate astrocytes in the rodent Hb (Gampe et al. 2012), was instead upregulated in human LHb-associated astrocytes (**Fig. 6I**). Notably, *SLC6A11* and *SLC1A2*, which encode astrocytic transporters involved in GABA and glutamate synaptic clearance, respectively, were significantly upregulated in LHb-associated astrocytes relative to MHb-associated astrocytes (**Fig. 6I**). In contrast, genes implicated in astrocyte-mediated regulation of LHb firing dynamics in rodents (Y. Cui et al. 2018; Xin et al. 2025) showed limited differential expression between human MHb- and LHb-associated astrocytes (*KCNJ10* and ionotropic glutamate receptors; **Table S12**), suggesting that these regulatory mechanisms may be more broadly shared across human Hb domains.

Altogether, these analyses identify putative molecular communication within spatially organized human Hb cell types, including interactions that may mediate crosstalk between the MHb and LHb and distinct signaling relationships between local neuronal and glial populations.

### 2.9 Neuropsychiatric and SUDs genetic risk is enriched in the medial habenula

How genetic risk translates into specific cellular contexts to produce neuropsychiatric disease phenotypes is unclear. We asked whether genome-wide association study (GWAS) signals for neuropsychiatric disorders, including SUDs, are enriched in human Hb cell types. First, we used *MAGMA* (Leeuw et al. 2015) to test enrichment of GWAS-associated gene sets among top Hb cell-type marker genes (**Methods 4.28**). We used both cellular- and extracellular-segmented VisiumHD data with *BANKSY* and *FICTURE* clusters (**Fig. S19**) as well as the snMultiome RNA assay to test GWAS enrichment. We reasoned that convergence across these biological and technical factors would provide supporting evidence for robust genetic risk within the human Hb (**Fig. 7A-C, Methods 4.28**). As an initial observation, enrichment for psychiatric GWAS genes is more consistent than SUD GWASs across cellular and spatial contexts for gene expression (**Fig. 7A-C**). Intriguingly, MHb shows increased enrichment of risk genes relative to LHb (**Fig. 7A**), with psychiatric GWASs accounting for the vast majority of enrichment. At cell-type resolution, enrichment is concentrated in Substance P MHb_A and MHb_C cell types (**Fig. 7C**). Overall, we identify consistent enrichment for externalizing, internalizing, MDD, and SCZ, as well as SUD GWAS genes in MHb across gene expression datasets. Within the LHb, panic disorder and SUD genes are enriched specifically in LHb_A, suggesting this LHb cell type may be particularly vulnerable compared to other LHb cell types. In the extracellular spatial data, enrichment is restricted to neuronal *FICTURE* and *BANKSY* clusters, with the vast majority of signal captured by *FICTURE* factors (**Fig. 7D**). This difference likely reflects the substantially finer spatial resolution of *FICTURE* within the extracellular compartment compared to *BANKSY* (**Fig. S19**). Given the highly heterogeneous composition of extracellular neuropil, this finer spatial decomposition may better resolve distinct neuronal microenvironments and their associated GWAS signal.

**Fig. 7:**
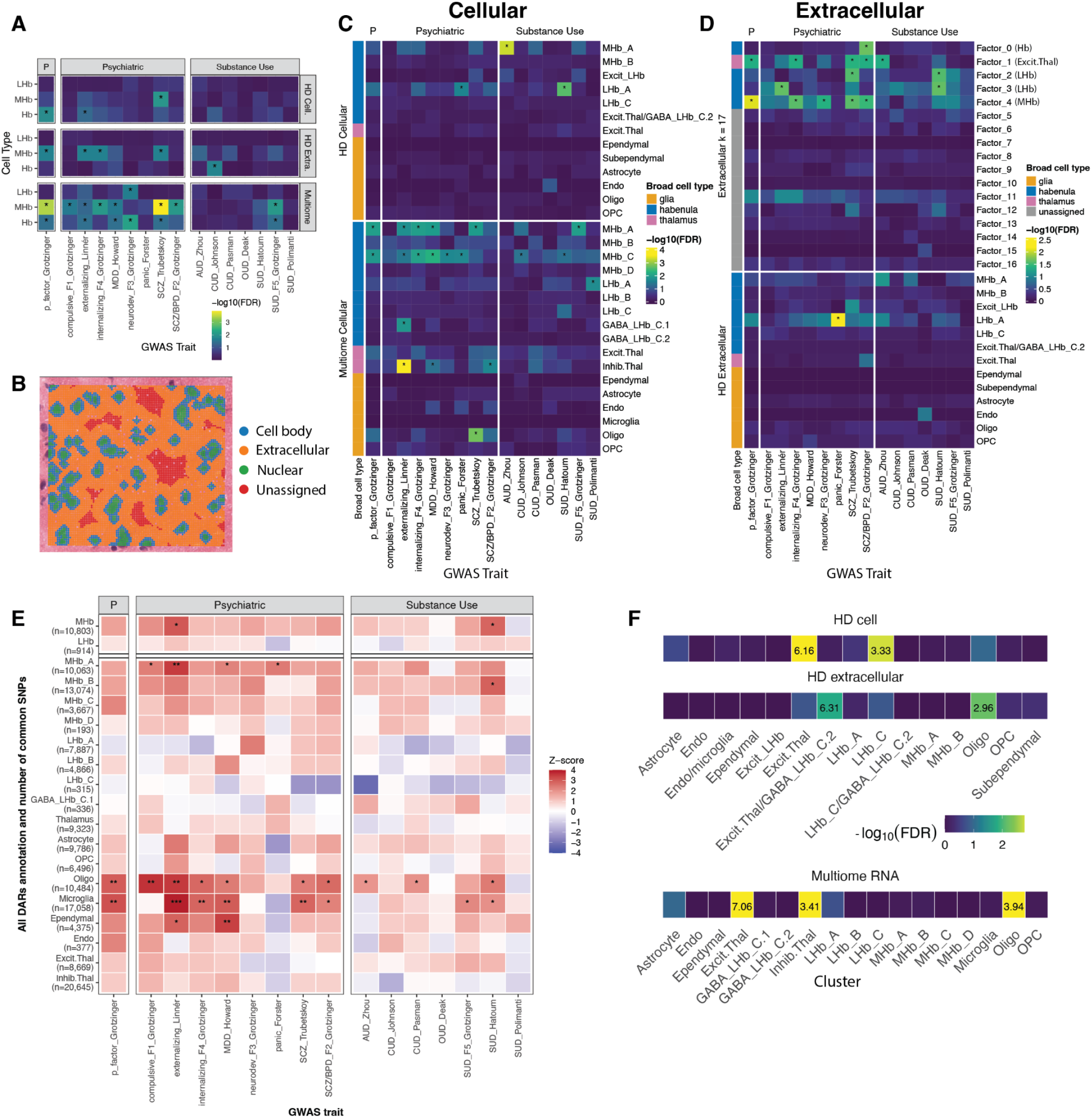
MHb gene expression and chromatin accessibility markers are enriched for genetic loci and gene expression variation associated with psychiatric and substance use disorders. **(A)** Heatmap of -log_10_ FDR-adjusted p-values showing whether top 200 *MeanRatio* marker genes per broad Hb cluster are associated with different GWASes across VisiumHD cellular, extracellular, and snMultiome datasets. ‘*’ denotes FDR < 0.05 with FDR computed by cluster across GWASes. GWASes were grouped by broad categories: psychiatric disorders, substance use disorders, or *P* factor (Grotzinger et al. 2026) (**Methods 4.28**). **(B)** Depiction of a small subset of 2um bin-level data showing the sets of bins used for HD analysis in (A), (C), and (D). HD cellular analysis used nuclear and cell-body bins; all extracellular analyses used bins in orange. **(C)** Similar to (A) but for all individual clusters from cellular datasets rather than collapsing LHb, MHb, or all Hb into one set. (**D**) Similar to (C) but for extracellular datasets rather than cellular ones; factors were computed with *FICTURE* (Si et al. 2024). (**E**) Heatmap of stratified linkage disequilibrium score regression (*S-LDSC*) associations between cell type chromatin marker regions (differentially accessible regions: DARs) and disorder-associated genetic variants from GWAS datasets. All DARs are shown in (E), while closed and open DARs are shown in **Fig. S29**. ‘*’ denotes FDR < 0.05 with FDR computed by cell type across GWASes. (**F**) One-sided Fisher’s exact test enrichment heatmap for differentially expressed genes associated with fentanyl self-administration in rat Hb (Magnard et al. 2026). Color scales indicate −log_10_(p), which were thresholded at −log_10_(p) = 3, and numbers within significant heatmap cells indicate odds ratios for the enrichments.

Since most GWAS signals reside in non-genic regulatory regions, we next asked whether these genetic associations were also evident within putative regulatory chromatin across Hb cell types. Using stratified linkage disequilibrium score regression (*S-LDSC*) in combination with the snMultiome ATAC data (Bulik-Sullivan et al. 2015), we tested enrichment of the same GWAS signals within the top differentially accessible regions (DARs) for each cell type (**Fig. 7E**, **Fig. S28**, **Fig. S29**, **Methods 4.29, 4.30**). Across DARs with either significantly increased or decreased accessibility in each cell type, oligodendrocytes account for the majority of significant GWAS enrichments, followed by microglia (**Fig. 7E**). Among Hb neurons, MHb_A Substance P and MHb_B cholinergic neurons are the only Hb cell types showing significant enrichment (**Fig. 7E**). Consistent with *MAGMA*, MHb shows greater *S-LDSC* signal than LHb, which is driven primarily by DARs with reduced accessibility in the corresponding cell type rather than those with increased accessibility (**Fig. S29**). In summary, across chromatin accessibility, nuclear gene expression, and gene expression with cellular and extracellular spatial clustering, the MHb, rather than the LHb, is the primary Hb subregion associated with genetic risk across neuropsychiatric and SUDs.

Finally, we asked whether disease-associated transcriptional changes identified by bulk RNA-seq could be mapped to specific human Hb cell types. Specifically, we leveraged a set of differentially expressed genes (DEGs) associated with chronic fentanyl self-administration in male rat Hb (Magnard et al. 2026), and tested enrichment among human cell-type marker genes across cellular and extracellular VisiumHD and snMultiome RNA datasets (**Fig. 7F, Methods 4.31**). Across modalities, rat fentanyl-associated DEGs are enriched in human LHb_C populations and thalamic cell types (**Fig. 7F**), suggesting the LHb_C group may be particularly sensitive to opioid-related transcriptional changes. Altogether, these analyses identify Hb cell types associated with genetic risk for neuropsychiatric and SUDs, as well as opioid-related transcriptional changes, highlighting distinct molecular substrates of disease susceptibility and transcriptional response within the human Hb.

## 3 Discussion

The Hb is a key component of reward circuitry that is implicated in neuropsychiatric disorders (Hu et al. 2020; Velasquez et al. 2014; F. Lin et al. 2026; Yalcinbas et al. 2025; Savitz et al. 2011) as well as an emerging target for region-specific therapeutic interventions (Hu et al. 2020; Z. Wang et al. 2024; Yang, Cui, et al. 2018), yet its spatio-molecular organization in the human brain is less defined compared to model organisms. Here, we integrated snMultiome and VisiumHD spatial transcriptomic profiling to generate a high-resolution cellular and spatial atlas of the adult human Hb. By leveraging conserved cell-type transcriptional programs across human, mouse, and zebrafish (Hashikawa et al. 2020; Pandey et al. 2018; Wallace et al. 2020), we derived evolutionarily informed cell-type annotations providing a cross-species framework for future translational investigation. From this consensus spatio-molecular atlas, we identified a rare population of GABAergic LHb neurons and characterized molecular programs associated with LHb and MHb neuron intrinsic physiology and neuromodulatory signaling, as well as cell type-specific regulatory relationships for disease-associated genes. We also resolved the spatial architecture of the human Hb, defining neuro-glial microenvironments, identifying patterns of cell-cell communication, and mapping neuropsychiatric genetic risk across genomic modalities onto spatially-organized Hb cell types.

Previous work established mouse-zebrafish Hb correspondences through joint transcriptional integration (Hashikawa et al. 2020; Pandey et al. 2018); we extend this framework by incorporating two human datasets and an additional mouse dataset while prioritizing cluster replicability independent of integration. Our annotations capture broad cellular populations, with potentially finer molecular and spatial specialization within these conserved groups, as demonstrated by the numerous MHb and LHb subdivisions in rodents (Hashikawa et al. 2020). While strong transcriptional correspondence supports conserved cellular identity, weak or absent correspondence does not necessarily indicate human-specific biology as populations may be missed because of sampling or taxonomic differences. Similarly, without nonhuman primate data, apparent mouse-human differences may reflect primate-enriched populations. Finally, transcriptional conservation does not establish evolutionary homology, as similar programs can arise through convergent evolution (Shafer 2019; Arendt et al. 2016). Rather than resolving cell-type evolutionary history (Shafer 2019; Arendt et al. 2016; Zhu et al. 2026), our framework aims to identify transcriptionally conserved populations that provide functionally relevant cellular correspondences across species.

Whether intrinsic GABAergic neurons exist within the Hb is unresolved, particularly given conflicting evidence from rodent studies regarding their presence and function (L. Zhang et al. 2018; Flanigan et al. 2020; Green et al. 2023; Nguyen et al. 2024; Quina et al. 2020). Moreover, reported GABAergic populations in rodents can co-release glutamate and function as excitatory rather than inhibitory neurons (Green et al. 2023; Nguyen et al. 2024; Quina et al. 2020). In our cross-species analyses, canonical GABAergic gene expression was conserved within the LHb_C cell type across vertebrates. However, snMultiome data further resolved two transcriptionally distinct GABAergic subpopulations in the human Hb without transcriptional evidence of glutamate co-expression. Orthogonal smFISH validation supported the presence of this rare population within the human Hb. Nevertheless, transcriptomic data alone cannot establish whether these neurons release GABA. Additionally, non-canonical GABA signaling has been documented in the rodent Hb (Kawai et al. 2025; U. Kim and Chung 2007; D.-G. Wang et al. 2006), emphasizing that molecular GABAergic identity does not necessarily imply inhibitory function. Future studies should further explore the synaptic targets and physiological effects of these GABAergic neurons. Together, these findings establish the presence of a rare GABAergic population in the human LHb, motivating future functional studies to understand its contribution to Hb circuit activity. The low abundance of this GABAergic population across humans, mice (Hashikawa et al. 2020; Wallace et al. 2020), and zebrafish (Pandey et al. 2018) snRNA-seq datasets suggests that substantially greater cellular sampling will be needed to fully characterize its molecular diversity.

Our analyses further support putative non-canonical GABAergic signaling in the human Hb, consistent with previous rodent studies (U. Kim and Chung 2007; D.-G. Wang et al. 2006; Kawai et al. 2025). Across mouse and human, *SLC12A5* (KCC2) follows a conserved expression gradient, with lowest expression in cholinergic MHb neurons and progressively higher expression across LHb populations, suggesting cell type-specific responses to GABA. Cholinergic MHb neurons also show the highest metabotropic GABA receptor expression, indicating distinct integration of chloride-dependent and -independent GABAergic signaling, consistent with rodent studies (Choi et al. 2016). Increasing neuronal *SLC12A5* (KCC2) expression to enhance inhibitory responses to GABA is being explored therapeutically in neurological disorders including Rett syndrome and epilepsy (Kahle et al. 2013; Tang et al. 2016; McMoneagle et al. 2024). Given that Hb hyperactivity is associated with MDD, KCC2 modulation may represent an additional potential therapeutic target within the human Hb.

Our snMultiome analyses revealed putative cell type-specific TF-cCRE-gene relationships, including regulatory interactions involving a range of ion channels and related genes. Because ion channel composition is a major determinant of neuronal activity, these relationships provide candidate mechanisms underlying the distinct physiological properties of Hb cell types. Given the established links between altered Hb firing dynamics and neuropsychiatric disease, cell type-specific regulation of ion channels may provide a molecular link between Hb cellular identity and disease-relevant physiology. By refining these regulatory networks around disease-associated genes, we identified candidate regulatory interactions that may help prioritize specific molecular mechanisms for functional investigation.

The small size of the human Hb has complicated resolving its spatial organization: high-resolution MRI cannot reliably distinguish the MHb from LHb (Rebecca P. Lawson et al. 2013), and spot-based Visium data similarly lacked sufficient resolution. The substantially higher resolution of VisiumHD enabled detailed molecular dissection of Hb architecture, complementing classic histological studies (Marburg 1944; Díaz et al. 2011) with cellular resolution and interrogation of cellular communication. Major Hb neuronal populations occupy distinct microenvironments, with highly ordered Substance P and cholinergic neurons in the MHb adjacent to ventricular ependymal and subependymal layers, whereas the LHb shows greater neuronal intermixing and extends into surrounding oligodendrocyte-rich white matter. Consistent with this organization, CCC analyses revealed stronger interactions between MHb populations and ependymal/subependymal cells and between LHb populations and oligodendrocytes, indicating distinct neural-glial signaling across the two domains. Whether the MHb and LHb communicate directly remains unresolved, but our analyses identify putative LR signaling between adjacent populations involving endogenous opioid, axon guidance, and contact-dependent ephrin signaling. Although these interactions require experimental validation, our findings demonstrate that Hb cell types occupy distinct cellular microenvironments and provide an anatomical framework for investigating how local interactions shape Hb function.

Astrocytes can directly modulate LHb burst firing in rodents (W. Cui et al. 2014; Y. Cui et al. 2018; Xin et al. 2025). In the human Hb, astrocytes showed strong associations with both the MHb and LHb_A, suggesting that astrocyte interactions extend beyond the LHb. Astrocyte-neuron LR signaling across the Hb was broadly dominated by axonal maintenance programs, whereas broader transcriptional comparisons of MHb- and LHb-associated astrocytes revealed higher expression of the GABA and glutamate transporters *SLC6A11* and *SLC1A2* in LHb-associated astrocytes, suggesting differential astrocyte-mediated regulation of the local synaptic environment. In contrast, genes previously implicated in astrocyte-mediated regulation of LHb burst firing showed limited differential expression between MHb- and LHb-associated astrocytes, suggesting that some aspects of astrocyte-neuronal regulation are shared across the Hb. Together, these findings suggest that astrocytes may contribute to both shared and domain-specific regulation of neuronal function across the human Hb.

Integration of GWAS-associated neuropsychiatric and SUD risk across multiple genomic modalities revealed greater enrichment of disease-associated genetic risk in the MHb than the LHb. In particular, internalizing, externalizing, MDD, SCZ, and SUD risk genes were consistently enriched across gene expression contexts of the MHb. These findings are notable given the disproportionate focus on the LHb in neuropsychiatric disease, particularly MDD, where LHb dysfunction has been extensively implicated (Bray 2018; R. P. Lawson et al. 2017; Li et al. 2011; Yang, Cui, et al. 2018; Yang, Wang, et al. 2018). Although cholinergic signaling in the MHb has been studied extensively in the context of nicotine addiction (Pang et al. 2016; Zheng et al. 2026; K. Kim and Picciotto 2023), our results suggest a broader role for the MHb in neuropsychiatric and SUD risk, particularly within the MHb substance P population, which showed enrichment of genetic risk in both transcriptional and regulatory profiles. Overall, the MHb may represent an underappreciated substrate of vulnerability to neuropsychiatric and SUDs and warrants greater consideration in studies of the cellular mechanisms linking genetic variation to disease.

This atlas defines conserved human Hb cell types and their distinct molecular, regulatory, and spatial contexts, providing a framework for understanding how cellular specialization shapes Hb function and disease vulnerability. As a resource to the scientific community, all data generated through this study is made available through interactive web applications.

## 4 Methods

### 4.1 Postmortem human tissue samples

Postmortem human brain tissue from the Lieber Institute for Brain Development (LIBD) Human Brain and Tissue Repository was collected through the following sites and protocols at the time of autopsy and/or organ donation: the Office of the Chief Medical Examiner of the State of Maryland, under the Maryland Department of Health IRB protocol #12–24, the Department of Pathology at Western Michigan University Homer Stryker MD School of Medicine, Gift of Life Michigan, and the County of Santa Clara Medical Examiner-Coroner Office, all under WCG IRB protocol #20111080. Demographics and other information for the 12 donors are listed in **Table S1**. Neurotypical controls were defined as those with no history of lifetime DSM5 psychiatric diagnoses and no lifetime substance use disorder diagnoses for ethanol, drugs of abuse, or prescription medications. Details of tissue acquisition, handling, processing, dissection, clinical characterization, diagnoses, neuropathological examinations, and quality control measures are previously described (Lipska et al. 2006). Fresh frozen tissue blocks containing the Hb were microdissected and stored at -80°C prior to experiments. Given the small size and midline position of the Hb, it can often be distorted during brain hemisectioning, slabbing, and flash-freezing, making it difficult to identify and dissect across donors. During dissection, multiple coronal slabs were inspected and several landmarks were used to estimate the approximate location of the Hb, including segmented putamen (SP), lateral geniculate nucleus (LGN), substantia nigra (SN), pulvinar nucleus of the thalamus (Th), and tail of caudate (C) (**Figure S2**).

### 4.2 Anatomical validation

To confirm the presence of Hb, each tissue block was subjected to anatomic verification via RNAscope multiplexed single molecule fluorescence in situ hybridization (smFISH) using the RNAscope Fluorescent Multiplex Kit v.2 and 4-plex Ancillary Kit (Advanced Cell Diagnostics ACD, Cat# 323100 and 323120) as previously described (Maynard et al. 2020). Given donor variability, a single expert experimenter carefully trimmed each block until white matter landmarks indicative of Hb became visible. Two serial sections were then collected on separate slides, and the latter slide was used for RNAscope to verify anatomical presence of Hb. The probes (Advanced Cell Diagnostics) and assigned Opal dyes (Akoya Biosciences) were as follows: *TAC3* - Opal Dye 570 (MHb marker; ACD, Cat# 507301; Akoya, Cat# FP1488001KT)*, GPR151* - Opal Dye 690 (LHb marker; ACD, Cat# 51137-C2; Akoya, Cat# FP1497001KT)*, POU4F1* - Opal Dye 620 (general Hb marker; ACD, Cat# 438441-C4; Akoya, Cat# FP1495001KT), and *MBP* - Opal Dye 520 (white matter tracts; ACD, Cat# 411051-C3; Akoya, Cat# FP1487001KT). Blocks passed anatomical quality control if co-expression of *GPR151* and *POU4F1* was detected at the midline in the expected dorsal-ventral position. Not all blocks initially expressed *TAC3* as MHb is located more centrally, and some blocks were not yet at an intermediate plane. Positive Hb signals were visually distinct from false positives, which included 1) fluorescent trapping from adjacent choroid plexus and blood vessels and 2) expression of *TAC3* in neighboring structures, such as the Th or LGN Individual blocks underwent multiple rounds of trimming and RNAscope until positive Hb signals were detected.

When the correct anatomical plane containing Hb was achieved, the block was scored with a razor to isolate Hb region and systematically cryosectioned for all assays as follows: 1) 2-10 um section banked that could be used for either RNAscope or VisiumHD, 2) 2-100 um sections for snMultiome. This sampling scheme was repeated 3 times resulting in 6 banked sections, and 3 tubes for snMultiome (**Fig. 1**). Each aliquot of tissue for snMultiome was weighed, with weights ranging from 30.1-83.6 mg. Lastly, an additional 2 cryosections were collected, and one was subjected to a final confirmation RNAscope to verify Hb was still within the sampled region. If Hb signal was not present, the final tissue aliquot for snMultiome would not be included.

### 4.3 snMultiome (snRNA-seq and snATAC-seq)

For 10 donors, we performed single nucleus RNA sequencing (snRNA-seq) and ATAC sequencing (snATAC-seq) with the 10x Genomics Multiome assay (Chromium Next GEM Single Cell Multiome Reagent Kit A, PN-1000284). We prepared nuclei from each donor using a modified version of the Nuclei Isolation from Complex Tissues for Single Cell Multi-ome ATAC + Gene Expression Sequencing (CG000375 Rev B). Briefly, this preparation involved first dissociating Hb tissue in lysis buffer using a glass dounce followed by incubation on ice for 2 minutes. Homogenate was then filtered and centrifuged at 500 rcf at 4℃ for 5 minutes (all centrifugation steps were carried out using these settings). Supernatant was removed, and the pellet was incubated in a buffer composed of 1X PBS, 10% BSA, and RNAse inhibitor (PBS+BSA Buffer) was gently introduced (1X phosphate buffered solution,Corning, Cat# 21-040-CV; MACS BSA stock solution, Miltenyi Biotec, Cat# 130-091-376; Protector RNAse inhibitor, Roche, Cat# 3335402001) for 5 minutes prior to resuspension. Homogenate was centrifuged again, supernatant was removed, and pellet was resuspended in 1 mL of the PBS+BSA Buffer for immunostaining with anti-NeuN488 (1:500), for 30-45 minutes on ice, mixing gently every 5 minutes (Sigma-Aldrich, Cat# MAB377X). After staining, the sample was centrifuged, supernatant was removed, then the pellet was resuspended in a minimum of 1 mL of PBS+BSA buffer passed through a 35 um filter. Finally, nuclei were labeled with 7-AAD (1:100; Sigma-Aldrich, Cat# SML1633-1ML).

Samples were sorted on a MoFlow Legacy (Beckman Coulter) cell sorter at the Bloomberg Flow Cytometry and Immunology Core. Samples were split sorted into NeuN+ neurons and NeuN-glia into respective 2 mL lo-bind tubes containing 500 uL of BSA and 5,000 units of RNAse inhibitor. While we aimed to enrich for neurons, this sorting strategy allowed us flexibility to spike in glial nuclei as needed to meet the minimum recommended amount of nuclei for proceeding with downstream snMultiome steps. To verify successful immunolabeling, a few microliters from each sample were sorted onto a glass slide and visualized with an EVOS microscope. We targeted a minimum of 76,000 sorted NeuN+ nuclei, and supplemented with NeuN-nuclei to reach this minimum if necessary. 9 of the 10 samples were NeuN enriched, donor Br8518 required glial spike-in (**Table S2**). Following sorting, nuclei were gently centrifuged, supernatant was removed, nuclei were permeabilized using 0.1 X lysis buffer for 2 minutes, and then washed with wash buffer (wash buffer contained the following reagents: Tris-HCl, NaCl, BSA, Tween-20, DTT, RNAse inhibitor, and nuclease free water). Nuclei were centrifuged, supernatant was removed, and the pellet was resuspended in 5uL of prepared diluted nuclei buffer. An aliquot of 1 uL of the resuspended nuclei were stained with trypan blue and nuclei were counted using a hemocytometer.

Approximately 11,025 - 19,069 nuclei were carried forward into the Chromium Next GEM Single Cell Multiome ATAC + Gene Expression (CG000338, Rev F) protocol, which was followed according to manufacturer’s instructions. Briefly, nuclei were subjected to transposition followed by GEM generation. Samples were then pre-amplified and split into gene expression (GEX) and ATAC fractions for library preparation. Sequencing was performed on an Illumina Novaseq6000 at the Johns Hopkins Single Cell Transcriptomics Core. 450,000,000 read pairs for GEX libraries and 450,000,000 read pairs for ATAC libraries.

### 4.4 snMultiome raw data processing

FASTQs were processed with *<u>Cell Ranger ARC</u>* <u>v2.0.2</u> (10x Genomics), which performs alignment, filtering, barcode counting, peak calling, and creates count matrices for both ATAC and RNA assays. The reference genome used was GRCh38-2020-A-2.0.0 (10x Genomics), which combines ENSEMBL v98 (Dyer et al. 2025) and GENCODE v32 (Frankish et al. 2019, 2023).

*<u>Cell Ranger ARC</u>* failed to perform joint cell-calling, due to algorithm restrictions and divergent quality control thresholds between assays. We thus performed a secondary analysis with *Cell Ranger ARC’s* reanalyze pipeline v2.0.2 using the barcode selection strategy. The reanalyze pipeline does a secondary analysis on the peak-barcode matrix (dimensionality reduction, cluster-based analysis, feature linkage and transcription factor analysis) by combining the previous outputs with a list of valid barcodes (barcodes with cells) extracted from the individual runs from <u>cellranger-count v7.2.0</u> (10x Genomics) for RNA and cellranger-atac count v2.1.0 (10x Genomics) for <u>chromatin accessibility</u>. For both analyses, the assay chemistry flag (--chemistry=ARC-v1) was specified to correctly recognize the data as standalone snMultiome assays.

The barcodes produced by each pipeline were translated into their corresponding snMultiome barcodes automatically by *Cell Ranger ARC* to associate ATAC and RNA data for the same nuclei. After translating the barcodes, we retained only the barcodes shared across both RNA and ATAC assays, leaving the dataset ready for downstream analysis (**Fig. S3A**).

*Cell Ranger ARC* detected a total of 26,401 nuclei. In contrast, *<u>Cell Ranger</u>* count on the RNA assay identified 84,177 nuclei, while *<u>Cell Ranger</u> ATAC* detected 78,921 nuclei in the ATAC assay (**Fig. S3B**). After filtering for valid barcodes identified independently in both RNA and ATAC assays, a total of 62,950 nuclei were recovered (63% of the 100K targeted nuclei, **Table S2**).

### 4.5 snMultiome data quality control

We evaluated nuclei quality and quantity in the *<u>Cell Ranger ARC</u>* reanalyze results based on two key criteria: (a) the presence of previously identified Hb-specific neuron marker genes (Yalcinbas et al. 2025), and (b) standard nuclei quality control metrics applied separately to the RNA and ATAC assays (**Fig. S4**).

To evaluate the Hb recovery rate (a), we measured and compared the percentages of global Hb-specific, and thalamus-specific neurons, in both standard *Cell Ranger ARC* and *Cell Ranger ARC* reanalyze log-count normalized data. Two marker gene sets were used for this analysis: “Literature-Based” (LB) and “Data-Driven” (DD) (Yalcinbas et al. 2025) (**Table S3**).

Next, we performed nuclei quality control using standard metrics (b) using isOutlier() from *scuttle* v1.12.0 (McCarthy et al. 2017) to identify outlier nuclei based on the median absolute deviation (MAD). Nuclei with high mitochondrial content, low RNA library size (Unique Molecular Identifier [UMI] counts), and low detected gene counts were flagged as outliers in the RNA assay (**Fig. S4A**). For the ATAC assay, low ATAC library size (UMI counts), low fragment counts, and low transcription start site (TSS) enrichment scores determined outlier low quality nuclei (**Fig. S4B**).

TSS enrichment is a widely used per-cell QC metric in single-cell and ATAC-seq, but its dynamic range depends strongly on library complexity and protocol (De Rop et al. 2024). Following current snATAC best practices, we used dataset-specific, distribution-based thresholds on TSS enrichment and related QC metrics rather than fixed ENCODE cutoffs (Heumos et al. 2023). Finally, the Nucleosome Signal (NS) metric used to evaluate nucleosome-free regions (NFRs) and mononucleosome regions in ATAC data was set at NS > 2 to exclude low quality nuclei, consistent with <u>regular analysis practices</u>.

6,043 (9.6%) of 62,950 nuclei with low quality in RNA were removed (either due to a high percent of mitochondrial gene expression, low UMI counts, or low number of detected genes), while only 1,466 (2.33%) nuclei with low quality nuclei were removed in ATAC (either due to a low TSS score, low UMI counts, or low fragment counts). After dropping low quality nuclei, a total of 55,702 (88.49%) high quality nuclei were retained for downstream analysis.

### 4.6 snMultiome cluster computation

Clustering was performed using the QC-filtered data. Integration by sample was performed with *Harmony* v1.2.0 (Korsunsky et al. 2019). For the RNA assay, we constructed a Shared Nearest Neighbor (SNN) graph using *Harmony*-corrected PCA embeddings (dimensions 1 to 30), and for ATAC assay we computed an SNN graph using *Harmony*-corrected LSI embeddings (dimensions 2 to 20). To integrate both modalities, we computed a Weighted Nearest Neighbor (WNN) graph using FindMultiModalNeighbors() from *Seurat* v5.0.1 (Hao et al. 2024) with the *Harmony*-integrated assays and a total of 30 nearest neighbors.

Batch correction was first performed using *Harmony* v1.2.0 (Korsunsky et al. 2019) on both the RNA and ATAC assays. Clustering at the gene expression level was performed using the Shared Nearest Neighbor (SNN) algorithm (Waltman and van Eck 2013; Stuart et al. 2019) on 30 *Harmony*-corrected reduced dimensions and with the Leiden algorithm (resolution = 0.8) (Traag et al. 2019). Clustering was implemented using FindNeighbors() and FindClusters() from *Seurat* v5.0.1 (Hao et al. 2024).

For clustering nuclei on the ATAC assay, dimensionality reduction was carried out using Latent Semantic Indexing (LSI) as implemented in *Signac* v1.13.0. This method computes term frequency-inverse document frequency (TF-IDF) (Hao et al. 2024), followed by singular value decomposition (SVD), as previously described (Stuart et al. 2019). LSI applies a two-step normalization procedure to account for differences in sequencing depth across nuclei and to prioritize rare peaks. RunTFIDF() from *Signac* was used with a default scale.factor of 10,000 to compute log(*TF* × *IDF*). The dimensionality reduction for both modalities was determined by observing the standard deviation of each reduction using the *Seurat* ElbowPlot() function.

To integrate the two modalities, a Weighted Nearest Neighbor (WNN) graph was constructed to identify nearest neighbors based on a weighted combination of the two modalities. The *Harmony* (Korsunsky et al. 2019) reduction (1 to 30 dimensions) was used for RNA, and the LSI reduction (2 to 20 dimensions) for ATAC, with the default number of multimodal neighbors (knn) set to 30. The FindMultiModalNeighbors() *Seurat* function was used. The Uniform Manifold Approximation and Projection (UMAP) (McInnes et al. 2020) technique was applied to the clustering results for visualization purposes (SNN, LSI and WNN).

For modularity optimization, 48 clustering combinations were run under three criteria: (a) four clustering algorithms: original Louvain (Blondel et al. 2008), Louvain with multilevel refinement (Rotta and Noack 2011), Smart Local Moving (SLM) (Waltman and van Eck 2013), and Leiden (Traag et al. 2019); (b) four clustering resolutions: 0.8, 1, 1.5, and 2 (the number of clusters increased with resolution value); (c) and three numbers of multimodal neighbors to compute (knn): 20, 30, and 40. The resulting 48 clustering results were filtered based on minimizing singletons (donor purity) and maximizing the representation of clusters across both modalities. Based on prior benchmarking (Heumos *et al*., 2023) and empirical evaluation, Leiden clustering at resolution = 2 with knn = 30 was selected for downstream analyses. It was overall similar to knn = 30 resolution = 1, knn = 40 resolution = 1, and knn = 40 resolution = 2 Leiden clustering results.

A total of 55,516 high quality nuclei resulting in 40 WNN clusters were obtained, having 19 Hb clusters identified, covering 55% (30,617 nuclei) of the total nuclei.

### 4.7 snMultiome cluster cell-type annotation

A preliminary WNN clustering annotation was performed through spatial registration (Maynard et al. 2020; Pardo et al. 2022; Huuki-Myers et al. 2024) of the RNA assay (40 WNN clusters; Leiden resolution = 2 and knn = 30) to our previous Hb single-nucleus annotated cell types (Yalcinbas et al. 2025). Briefly, spatial registration first computes differential expression *t*-statistics for each individual cluster within a dataset and then computes the correlation of cluster *t*-statistics for all pairwise cluster comparisons across datasets. This approach identifies matched cross-dataset clusters with shared transcriptional variability, enabling an initial snMultiome cluster annotation grounded in our previous Hb single–nucleus work (**Fig. S5**). Annotations were further adjusted based on hierarchical clustering. As an additional quality-control effort, we used *scDblFinder* v1.22.0 (Germain et al. 2022) with default parameters to identify putative doublet cells and dropped any doublet-enriched cluster (doublet proportion >= 50%, **Fig. S5**). This resulted in a final count of 35 fine resolution clusters and 54,193 nuclei for all downstream analysis. *Harmony* integration and dimension reduction was repeated as previously described in Methods 4.6 after dropping the doublet-enriched clusters.

These 35 cluster annotations were further refined based on several targeted marker gene analyses. Out of the two initial astrocyte clusters, one had clear expression of ependymal markers and was annotated as ‘Ependymal’ (*FOXJ1, TPPP3, PIFO, CD24, TMEM212, RARRES2*, **Fig. S5**). Out of the MHb annotated clusters, two matched to the MHb.1 label (C.10.MHb.1, C.14.MHb.1) and one matched to the MHb.2 label (C.07.MHb.2), where these correspond to the well described Substance-P and Cholinergic neuronal populations of the MHb. Cluster C.11 initially matched to both MHb.1 and .2, but due to high levels of cholinergic marker expression, was subsequently labeled as a MHb.2 fine resolution cluster (*CHAT, SLC5A7, SLC18A3*, **Fig. S5**).

Fine resolution snMultiome clusters that matched a single cluster in Yalcinbas and Ajanaku et al. were grouped together for a total of 17 mid-resolution clusters. These 17 clusters were the input for the subsequent cross-species analysis of transcriptional cell-type conservation (**Methods 4.8**).

### 4.8 Cross-species conservation of transcriptomic cell types with *MetaNeighbor*

The *MetaNeighbor* v1.28.0 (Crow et al. 2018; Fischer et al. 2021) framework was used to identify replicable transcriptomic clusters both within a dataset and across species to define conserved cross-species habenula cell types. *MetaNeighbor* scores cell-cell expression similarity to identify replicable cell types across datasets; when applied across species, it can identify conserved cell types. In all cases, the top 2,000 highly variable genes were used when running *MetaNeighbor* functions. To start, the Wallace et al. mouse and Pandey et al. zebrafish datasets were split by sample and *de novo* clustered using *Seurat* functions: NormalizeData( normalization.method = ’RC’, scale.factor = 1e6), FindVariableFeatures(nfeatures = 2000), ScaleData(), RunPCA(), FindNeighbors(dims = 1:20), and FindClusters(resolution = 0.8). We identified replicable metaclusters of these cross-sample *de novo* clusters using default and BestVsNext *MetaNeighbor* functions: MetaNeighborUS(), MetaNeighborUS(one_vs_best = TRUE, symmetric_output = FALSE), and extractMetaClusters(). These metaclusters were annotated through marker gene expression (**Fig. S6**). We identified replicable mouse cross-dataset clusters between the Wallace et al. metaclusters and the finely annotated Hashikawa et al. dataset to align all mouse cell-type annotations (**Fig. S6**).

When performing cross-dataset comparisons with *MetaNeighbor*, all datasets were first filtered to the subset of shared genes. In the case of cross-species comparisons, all datasets were filtered to the subset of shared one-to-one orthologs, sourced from ENSEMBL Release 115. For the full cross-species analysis presented in **Fig. 2A-B**, this was a set of 9,522 one-to-one orthologs. Pseudo-one-to-one orthologs were constructed for zebrafish by summing the counts of one-to-many ortholog matches to human genes, to account for the numerous A and B paralog pairs present within zebrafish due to a whole genome duplication, similar to (Hahn et al. 2023). Groups of cross-species clusters were identified with MetaNeighborUS() (**Fig. 2A**), MetaNeighborUS(one_vs_best = TRUE, symmetric_output = FALSE), and makeClusterGraph(low_threshold = 0.5) (**Fig. 2B**).

From the final cross-species consensus cell-type annotations (**Fig. 2F)**, the top 50 1-vs-All markers per snMultiome cell type were derived as in **Methods 4.11**. Duplicate genes in these top 50 marker sets were assigned to their higher ranked cell type determined by fold change; for example, a gene ranked as the top 5th marker for cell-type A compared to the top 40th marker for cell-type B was retained as a cell-type A marker **Table S9**. Per-cell marker set enrichments of these top ∼50 markers were computed for the mouse habenula datasets (**Fig. S8**) and the VisiumHD data (**Fig. 5**) using the *MetaMarkers* v0.0.1 (Fischer and Gillis 2021) functions: score_cells(log1p(cpm(expression))) and compute_marker_enrichment(). Briefly, these functions compute expression enrichment scores within individual cells for each marker set under the null of equal expression across all marker sets. We *Z*-scored these enrichment values for visualization purposes in **Fig. S8** and **Fig. 5**.

### 4.9 LHb_C subclustering to identify Hb GABAergic neurons

The RNA assay of LHb_C neurons was integrated across donors with default *Seurat* functions: FindIntegrationAnchors() and IntegrateData(). The integrated data was then clustered as in **Methods 4.8** (**Fig. S9N**), but with a resolution parameter of 0.25 in FindClusters(resolution = 0.25) to isolate the clear SLC32A1+/GAD2+ cells as in **Fig. 2C**

### 4.10 Inhibitory Population Validation

To confirm the presence of local inhibitory cells within the Hb, RNAscope was performed on Hb sections from four donors. The same procedure and products were used as listed in the anatomic verification section above with the exception of probes. The following probe and opal dye combination was used: *SLC17A6* (ACD, Cat#415671) - Opal dye 620 , *MBP* (ACD, 411051-C2) - Opal dye 520, *SLC32A1* (ACD, Cat# 415681-C3) - Opal Dye 570, and *POU4F1* (ACD, 438441-C4) - Opal Dye 690. The resulting fluorescence was then imaged on an Akoya Biosciences *PhenoImager* at 20X magnification. After imaging, *Phenochart* was used to annotate each section to allow for fluorescence unmixing. Unmixing was completed using *InForm*, the resulting unmixed tiles were then stitched together in *HALO*.

*HALO* was then used to perform analysis to identify positive cells. First an annotation was created for every section where the ependymal region and thalamus were excluded to avoid false positive calling given the large amount of highly positive VGAT and VGLUT cells in these regions. Using the FISH-IF module in *HALO*, DAPI was treated as an FISH dye and all other opal dyes were treated as IF dyes. 12 phenotypes were created to identify various cell types. After adjusting the parameters of each dye to avoid over or under calling puncta analysis was performed. The resulting data were then exported in .csv files.

For each section, cells were initially annotated as POU4F1+, SLC32A1+ (VGAT) or POU4F1/SLC32A1+ based on a probe threshold of >= 5. A boundary box to approximate the area of the habenula was computed using the 5th and 95th percentiles of the *X* and *Y* coordinates for the POU4F1+ cells, as in **Fig. S9**. We used a sliding window approach to compute the average probe counts within the habenula region, using a window that spanned the length of the habenula region and moved medial->lateral through the habenula region, with a width of 100 units and a step size of 25 units (**Fig. S9**).

### 4.11 Cell-type marker quantification

Briefly, we computed two forms of markers depending on the downstream application: 1-vs-All markers and *MeanRatio* markers (Huuki-Myers, Montgomery, et al. 2025). 1-vs-All markers were computed for the cellular VisiumHD data using registration_wrapper() from *spatialLIBD* v1.20.1 (Pardo et al. 2022), which internally pseudobulks data by cluster, performs basic expression filtering, and generates enrichment statistics by gene. We filtered to retain genes with positive fold changes (greater expression in the target cluster) that were significant (FDR < 0.05). In Fisher’s exact test (**Methods 4.31**) and *MAGMA* (**Methods 4.28**), we used *MeanRatio* markers as gene sets (**Table S13**). In each relevant dataset, we first pseudobulked by cluster with registration_pseudobulk() from *spatialLIBD* v1.20.1 (Pardo et al. 2022), then applied get_mean_ratio() from *DeconvoBuddies* v1.1.5 (Huuki-Myers, Montgomery, et al. 2025) on logcounts. We filtered to genes with *MeanRatio* > 1.05; *MeanRatio*s represent the ratio of mean expression in the target cluster to the mean expression in the next-highest cluster. Finally, we retained up to 200 genes per cluster, ranking by *MeanRatio*. For the HD cellular, HD extracellular, and snMultiome datasets, we ran the *MeanRatio* workflow at three cell-type resolutions: grouping together all Hb cell types, grouping together MHb and grouping LHb cell types (**Fig. 7A**), and finally leaving the individually annotated clusters as-is (**Fig. 7C,D**). HD extracellular markers were computed by aggregating expression from extracellular bins (**Methods 4.22**) by their corresponding cells and assigning the cells labels determined in the cellular clustering and annotation (**Methods 4.23**).

### 4.12 *MetaNeighbor* and cell-type marker GO term enrichment

GO terms with >= 10 and <=100 genes were used in two complementary analyses to characterize transcriptional profiles of Hb cell types. First, using only the Hb neuronal cell types, the *MetaNeighbor* function MetaNeighbor(genesets = go_sets) was used to quantify the strength of each GO term in predicting cell-type transcriptional profiles (AUROC metric, *X*-axis **Fig. S7E**). Second, using the top 50 1-vs-All markers per cell type, enrichment for the same GO terms was computed using Fisher’s Exact test, implemented through the hypergeometric distribution. Specifically, the R function phyper(overlap - 1, go_length, universe_term, celltype_length, lower.tail = FALSE) was used to compute p-values for each GO term, where overlap is the intersection between a GO term and a marker set, go_length is the number of genes in the GO term, celltype_length is the number of genes in the marker set, and universe_term is the total number of background genes minus the number of genes in the GO term. The number of background genes was the full gene annotations of the snMultiome data; GO terms were filtered to this background gene set. P-values from phyper() were FDR-adjusted using p.adjust(method = ’BH’) (*Y*-axis **Fig. S7E**).

### 4.13 snMultiome peaks identification

To construct a unified peak set for chromatin accessibility analysis across clusters, we computed peak calling on 18 mid-resolution WNN clusters with Signac::CallPeaks() v1.16.1 (Stuart et al. 2021), which internally uses *MACS3* v3.0.4 (Y. Zhang et al. 2008; Gaspar 2018), and grouped by cluster. Accessibility over the union of resulting peaks was quantified with FeatureMatrix() from *Signac* v1.16.1 (Stuart et al. 2021), and a new chromatin assay was generated. GC content was computed using RegionStats() and the new assay was normalized by TF-IDF.

### 4.14 Identification of peak-gene-transcription-factor trios

Peak-gene-transcription-factor correlation trios were computed independently by cell type, first filtering out unexpressed genes and peaks. Endo and MHb.3 cell types were dropped due to having less than 200 cells, a cutoff used to ensure meaningful estimation of trio relationships. Transcription factor (TF) binding motifs were obtained using *JASPAR2020* v0.99.10 (Baranasic 2020), and motif accessibility deviations were computed with *chromVAR* v1.30.1 (Schep et al. 2017) through the *Signac* v1.16.1 wrapper RunChromVAR()(Stuart et al. 2021). Following the recommended *TRIPOD* v1.0.0 workflow (Jiang et al. 2022), clustering resolutions were explored for forming metacells (small groups of highly similar cells), the units with which *TRIPOD* computes trios (**Fig. 4A,C**). TRIPOD::optimizeResolution() was run with resolutions 0.5, 1, 2, 3, 4, 6, 8, 12, 16, 24, and 32 for each cell type, where higher resolutions tend to generate more metacells composed of fewer cells. In the three cell types where all criteria could be met, resolution was decided by choosing the value yielding the smallest number >= 80 of metacells where at most 20% of metacells had at least 20 constituent cells. In the remaining 14 cell types, the resolution yielding the minimal number of metacells >= 40 was used (**Fig. S10A,C**). Metacells, while often composed of some cells from multiple donors, tended to mostly aggregate cells from the same donor (**Fig. S10B**). After metacells were obtained, trios were computed with TRIPOD::getTrios(fdr.thres = 0.05, sign = “positive”, model.name = “TRIPOD”) for test levels 1 and 2, matching by *Y_j_*, which represents controlling for transcription factor expression. Briefly, level 1 tests for peak-gene correlations that exist when transcription factor expression is held roughly constant; level 2 tests for an interaction effect where increasing transcription factor expression induces larger gene-expression increases per unit increase in peak accessibility. We retained only such trios where both gene expression and peak accessibility were positive in at least 10 metacells (70.5%, **Table S10**). In total, we found 74,075 level 1 trios and 134,725 level 2 trios. To form a more robust set of results, the intersection of trios was taken across *TRIPOD* test levels 1 and 2, yielding 358 robust trios across cell types (**Table S10**).

### 4.15 Network of transcription factors and their putative regulated genes

*RCy3* v2.28.1 (Gustavsen et al. 2019) was used to generate *Cytoscape* v3.10.4 (Shannon et al. 2003) gene-TF networks based on level 2 trios. Level 2 trios were specifically selected to capture cases where increasing TF expression mediates gene-expression increases through an interaction with peak accessibility. To keep networks focused and visually feasible, only habenula and astrocyte trios were considered, and trios were filtered to include just those with the largest 50% of coef values (indicating strongest interaction effect). We limited genes and TFs to those associated with risk for MDD or substance dependence as determined by *OpenTargets* (**Methods 4.26**) (Buniello et al. 2025). For each TF, only the genes present in the top 3 trios by coef were displayed.

### 4.16 Visium H&E data generation

Spatial transcriptomics was performed on three donors using the Visium Spatial Gene Expression platform from 10X Genomics (PN-1000184). Each Visium slide contained four capture areas from the same donor on which one scored Hb section was placed at the time of initial cryosectioning. We prioritized the anterior-posterior level at which both MHb and LHb were present, with the four sections collected 110um apart. Visium slides were processed following the Visium Spatial Gene Expression Reagent Kits user guide (CG000239, Rev C). Briefly, slides were fixed in chilled 100% methanol for 30 minutes, then stained with hematoxylin and eosin prior to imaging on a Leica CS2 slide scanner. Slides were then decoverslipped and sections were permeabilized for 18 minutes and a reverse transcription reaction was performed before completing second strand synthesis. cDNA amplification was performed, amplified cDNA was purified, and cDNA concentration was determined before proceeding to library generation. Library generation required fragmentation and ligation prior to adding a unique sample index. Libraries were quality controlled and sequenced at Johns Hopkins Single Cell and Transcriptomics Core targeting 60,0000 read pairs per spot on an Illumina Novaseq X.

### 4.17 Visium H&E data processing

Similar to our other Visium H&E projects (Huuki-Myers, Divecha, et al. 2025; Huuki-Myers et al. 2024; Maynard et al. 2021), sample slide images were first processed using *VistoSeg* (Tippani et al. 2023). *VistoSeg* was used to divide the Visium sample slides into individual capture areas using the VistoSeg::splitSlide() function. Individual capture area images from *VistoSeg* were then spatially aligned to Visium Slide capture areas using the *Loupe Browser* (version 7.0.1, 10x Genomics). This alignment enabled precise correspondence between histological features and individual capture spots on the slide.

The Visium FASTQ and image data were processed with the 10x *<u>SpaceRanger</u>* pipeline v2.1.0, using human reference genome GRCh38 and 2020-A annotation. This results in 33,641 total spots in-tissue, with a median 3,129 spots a sample (min=1,711, max=3,815). Counts were detected for 26,614 genes. In-tissue spots have a median of 873 UMIs, and a median of 405 detected genes (**Table S14**).

For quality control, localOutliers() from *SpotSweeper* v1.3.0 (Totty et al. 2025) was applied to identify local outliers in three quality control metrics: low library size, low detected gene count, and high rate of mitochondrial gene expression. A total of 232 spots (0.69%) were discarded as part of the spot-level quality control resulting in 33,409 total spots available for spatial domain clustering.

### 4.18 Visium H&E spatial domain identification and annotation

Principal component (PC) analysis was run with *scater* v1.34.0 (McCarthy et al. 2017) using the top 1,198 (10%) highly variable genes detected with getTopHVGs() from *scran* v1.34.0 (Lun et al. 2016). Batch correction was then performed with *Harmony* v1.2.3 (Korsunsky et al. 2019), using RunHarmony(group.by.vars = c(’sample_id’, ’brain_id’)) on the PCs. Visium spots were then clustered into spatial domains (SpD) using *BayesSpace* v1.16.0, a spatially aware clustering method (Zhao et al. 2021), using the function spatialCluster(nrep = 50000)on a range of *k* (equivalent to *q* for *BayesSpace* notation) values from 2 to 28 in increments of 1. Spatial registration of the SpDs against multiome clusters was performed with registration_wrapper() from *spatialLIBD* v1.21.5 (Maynard et al. 2020; Pardo et al. 2022; Huuki-Myers et al. 2024). However, even at *k* = 28, *BayesSpace* SpDs failed to identify separate MHb and LHb SpDs (**Fig. S12**). The Visium H&E data can be interactively explored on the *spatialLIBD* interactive website we provided for it.

### 4.19 VisiumHD Data Generation

Fresh frozen tissue sections banked for VisiumHD were processed in pairs using the VisiumHD Spatial Gene Expression platform (10x Genomics, Pleasanton, CA) according to manufacturer’s protocol. Briefly, sections were removed from -80°C storage, tissues were fixed in chilled methanol, dehydrated in isopropanol, and then H&E-stained following the procedures outlined in CG000684 rev B. In brief, Gills II Hematoxylin was applied uniformly over the tissue for 1 minute, hematoxylin was discarded and the tissue was washed in Milli-Q ultrapure water prior to bluing buffer application. After an additional wash in Milli-Q ultrapure water, slides were immersed in alcoholic eosin and subsequently washed in Milli-Q ultrapure water. Slides were then temporarily mounted in 85% glycerol and imaged on a Leica Aperio CS2 digital pathology slide scanner (Leica Biosystems) equipped with a color camera using a 20x and 40x 0.75 NA objective and saved as Tagged Image Files (.tiff). Following coverslip removal, Regions of Interest (ROIs) were selected through precise application of the Visium Cassette S3 gasket (CG000730 Rev A). ROIs were subsequently destained in 0.1N HCL for 15 minutes at 42°C and washed in Tris-EDTA (pH 8) followed by 1x PBS pH 7.4.

Steps pertaining to *in situ* hybridization, ligation, CytAssist-Enabled probe release, and library construction steps were then followed according to manufacturer’s instructions in CG000685 rev B. Following a 15 minute permeabilization at room temperature in a 0.7% Tween-20/PBS, the Visium Human Transcriptome probe kit v2 (PN-1000466) was applied to each ROI and allowed to hybridize to accessible RNA at 50°C overnight. Following an 18-22hr incubation, hybridized slides were washed in FFPE Post Hyb Wash Buffer and ligated using a ligation enzyme diluted in 2x probe ligation buffer for 1 hour at 37°C. While tissues were washed in post ligation wash buffer and then allowed to acclimate to room temperature, a VisiumHD slide was removed from -80°C storage and warmed to room temperature for 30-60 minutes, after which it was washed in 0.1x SCC. The VisiumHD slide was then placed in a Visium 2-port cassette S2 and RNase enzyme was allowed to equilibrate the capture areas. During the VisiumHD slide equilibration, the two tissue slides were stained in 10% Alcoholic Eosin, washed in PBS, and then mounted on the Visium CytAssist instrument. After the VisiumHD slide equilibration and subsequent drying, the CytAssist instrument run was initiated.

After the 30-minute instrument run, during which time ligated probes on tissue slides were transferred and captured on the VisiumHD capture areas, the VisiumHD slide was washed in Buffer EB and immediately placed in the Visium 2-port cassette. Extension enzyme and buffer were then applied to each capture area and probes were extended. Following probe extension, probes were eluted and then pre-amplified to generate ample material for downstream library construction. To determine the ideal number of additional cycles needed to amplify each library for the sample index PCR, pre-amplified samples were quantified using KAPA SYBR Fast qPCR Master Mix Universal (Roche KK4600) on the CFX Opus 96 (Bio-Rad 12011319). Unique indices from the dual index plate TS Set A were assigned to each sample so that VisiumHD libraries could be pooled and demultiplexed in downstream sequencing runs. All 5 VisiumHD Gene expression libraries were sequenced on a single lane of Illumina’s NovaSeq X at Psomagen (Rockville, MD) using the following parameters: R1(43)-i7(10)-i5(10)-R2S(50). An average of 659,123,173 reads per sample were generated across the five samples.

### 4.20 VisiumHD data processing

The VisiumHD FASTQ and image data were processed with the <u>10x *SpaceRanger* pipeline</u> v4.0.1, using human reference genome GRCh38 and 2024-A annotation (**Table S15**). This resulted in 3,511,220 total 8μm bins in-tissue, with a median 490,166 bins per sample (min=395,569, max=557,860. Counts were detected for 17,931 genes. In-tissue bins have a median of 108 UMIs, and a median of 92 detected genes.

### 4.21 VisiumHD quality control

Briefly, quality control was performed to remove technical artifacts, while preserving most of the data. Such an artifact was observed only in Br8433, marked by characteristically low total UMI counts on the rightmost portion of the tissue (**Fig. S13A**). To identify the exact boundary on the *X* axis (array_col), a sliding-window approach was utilized by observing how mean UMI counts within a 10-unit_array_col_ window change as the window was moved laterally across the tissue. At array_col = 783, whose sliding window includes 8μm bins satisfying array_col >= 783 and array_col < 793, mean UMI qualitatively shifts from a mild, jagged decrease to a perfectly smooth, precipitous decline, suggesting low-UMI bins begin at array_col = 793 (**Fig. S13B**). To ensure problematic bins were excluded from all analyses (i.e. those involving 2μm bins and individual cells), the vertical nature of the artifact was exploited, and spatial coordinates corresponding to array_col > 793 (i.e. pxl_col_in_fullres > 34326) were excluded in Br8433.

For all analyses, only in-tissue bins or cells were kept, and genes, bins, or cells with no expression were dropped. For cell-level analyses, cells with less than 10 total UMI were dropped. As each capture area initially included two spatially adjacent sections, dbscan(eps = 300) from *dbscan* v1.2.2 (Hahsler et al. 2019) was used to partition capture areas into a total of 10 tissue sections (two per capture area), with sample_id in each downstream *SpatialExperiment* v1.18.1 object (Righelli et al. 2022) referring to tissue section in all VisiumHD analyses.

### 4.22 VisiumHD: determining the number of spatial factors k for *FICTURE*

Briefly, spatial factors were computed with *FICTURE* v0.0.5.2 (Si et al. 2024) with two sets of input data: all bins and just extracellular bins (**Fig. S14A**). To define extracellular bins, bin2cell.expand_labels(max_bin_distance = 7) from *bin2cell* v0.3.3 (Polański et al. 2024) was run on cellular bins (i.e. labels_he_expanded), and the set difference with cellular bins was taken, extracting just the set of bins surrounding each defined cell. For both extracellular and all-bin datasets, raw 2 μm bin-level counts were library size normalized and log2 transformed [log_2_(x + 1)] with computeLibraryFactors() and logNormCounts() from *scuttle* v1.18.0 (McCarthy et al. 2017). Batch correction was performed using cleaningY(p = 1) from *jaffelab* v0.99.34 (Collado-Torres et al. 2024) to fit a linear model and regress out the effect of the capture area. Following this residualization, any previously zero counts were set to zero, preserving sparsity of the data. Doing so is important to enable use of memory-efficient sparse matrix representations that render downstream analyses feasible. Next, in the rare case when negative counts were introduced by residualization (n = 15 and 14 genes for all-bin and extracellular datasets, respectively), a small constant was added to the affected genes’ counts such that the minimum non-zero count after correction was equal to the before-correction non-zero minimum count. The log_2_(x + 1) transformation was reverted by applying the expression 2^x^ - 1 for counts x. To cluster all samples under the same labels, a single input TSV file was generated from the five capture areas’ counts, where spatial coordinates for each capture area were staggered and placed adjacently. A buffer of at least 5% of the length of the largest capture area was placed between each capture area to ensure different samples were not clustered similarly near their shared boundaries. Finally, spatial factors were computed with ficture run_together from *FICTURE* v0.0.5.2 (Si et al. 2024). For the all-bin data, candidate numbers of factors including 3, 4, 5, …, 40, 70, and 100 were computed. We selected the minimal k where spatial registration (Maynard et al. 2020; Pardo et al. 2022; Huuki-Myers et al. 2024) to snMultiome cell types using layer_stat_cor(top_n = NULL) from *spatialLIBD* v1.20.1 (Pardo et al. 2022) indicated clear and distinct factors corresponding separately to Hb and thalamus: it was k = 8. For the extracellular data, ficture run_together was run with k = 17 factors.

### 4.23 VisiumHD cell type identification

To define cells, *bin2cell* v0.3.3 (Polański et al. 2024) was used (**Table S16**). Nuclear segmentation of each capture area was performed with bin2cell.stardist(prob_thresh=0.01), defining the cell body with a small region (4 μm dilation) surrounding each nucleus using bin2cell.expand_labels() (**Fig. S14A**). 2 μm bin-level gene expression was aggregated by the cellular segmentations with bin2cell.bin_to_cell().

To first find spatially variable genes (SVGs), rasterizeGeneExpression(fun = ’sum’) from *SEraster* v1.0.0 (Aihara et al. 2024) was run for each sample on the 8μm bin-level data, specifying a sample-specific resolution such that rasterized bins were 100μm wide. Doing so reduces the data size and computation time for SVG identification while still identifying SVGs as noted by *SEraster*’s documentation (Aihara et al. 2024). *nnSVG* v1.12.0 (Weber et al. 2023) was then run after expression-based filtering. Specifically, nnSVG::filter_genes() was applied such that non-mitochondrial genes with at least 3 counts in 5% of rasterized 100μm bins were retained. Then, bins with less than 10 gene counts were dropped; a mean of 7351 genes (41%) was retained. To select dataset-wide SVGs, ranks for genes considered in all ten samples were averaged, and the top 1,000 SVGs by average rank were chosen. After subsetting to top SVGs and staggering spatial coordinates for each tissue section (sample_id) such that each lay in an adjacent range, a dataset-wide embedding was computed on logcounts with computeBanksy(compute_agf = TRUE) from *BANKSY* v1.4.0 (Singhal et al. 2024). PCA was performed on the embedding, followed by RunHarmony() from *Harmony* v1.2.3 (Korsunsky et al. 2019), harmonizing by tissue section (sample_id). Finally, Leiden clustering of the *Harmony*-corrected embedding was performed with Banksy::clusterBanksy() using lambda = 0.2 at candidate resolutions 0.1 through 2 (in increments of 0.1), 4, and 8 before selecting the optimal resolution value.

To determine the optimal clustering resolution, an average of three metrics was taken for each resolution, with resolution = 1.8 holding the highest average (**Fig. S15A**). Briefly, spatial registration (Maynard et al. 2020; Pardo et al. 2022; Huuki-Myers et al. 2024) was performed for each clustering resolution against cell types defined in the snMultiome data, and metrics aimed broadly to favor resolutions whose clusters registered uniquely to a variety of cell types. Correlation of *t*-statistics between datasets was computed with layer_stat_cor(top_n = NULL) from *spatialLIBD* v1.20.1 (Pardo et al. 2022). The first metric measured the fraction of non-Hb cell types (case A, **Fig. S15**) that had at least one cluster registering only to that cell type with layer_confidence == ’good’ after using spatialLIBD::annotate_registered_clusters(cutoff_merge_ratio = 0.1) (Pardo et al. 2022). Similarly, the second metric measured the fraction of Hb cell types with good unique mappings (case B, **Fig. S15**). The third metric measured how many Hb subtypes had corresponding registered clusters, allowing for non-unique mappings to potentially multiple MHb or multiple LHb cell types (e.g. some clusters may map to both MHb.1 and MHb.2, but to no lateral or non-Hb cell types (case C, **Fig. S15**). Finally, for each clustering resolution the number of clusters mapping ambiguously (in such a way as to not to count towards the first three metrics) was counted, and resolutions with more than 5 such clusters were excluded from consideration (ambiguous, **Fig. S15B**). Using these spatial registration-based metrics, we determined that the optimal resolution was 1.8.

We then annotated the resulting 27 clusters at resolution = 1.8 using the spatial registration against the snMultiome data (**Fig. S16**). Briefly, uniquely mapping clusters were assigned the corresponding cell type; multi-mapping clusters were resolved via unique mappings to the fine-resolution snMultiome data (clusters 8 and 15) (**Fig. S16B**) or using spatial distribution (clusters 9, 13, and 14) (**Fig. S16C,D**). Multi-mapping clusters with no clear resolution via those methods (clusters 2, 5) were given dual labels with the two cell types to which they registered (e.g. Endo/microglia) (**Fig. S16A**). Finally, cluster 16 exhibited transcriptional ambiguity and sample specificity (notable presence only in Br9902; see **Fig. S18A**), for which it was dropped from further analyses (**Fig. S16E**).

### 4.24 Anatomy deformation of the Br9902 donor

Tissue sections from Br9902 exhibited a markedly different spatial organization from the remaining four donors. In both sections, the ependymal/subependymal layers extend deep into the tissue and the relative positions of MHb_A and MHb_B are reversed, suggesting alteration of Hb dorsal/ventral anatomy during slab freezing rather than true biological variation (**Fig. S21**, **Fig. S25**). Because cell colocalization and cell-cell communication analyses depend on accurate spatial relationships between cell types, this donor was excluded from coordinate-based downstream analyses.

### 4.25 Quantifying spatial arrangement of cell types with *CRAWDAD*

Relative spatial arrangements of VisiumHD cell types were computed with *CRAWDAD* v1.0.1 (Dos Santos Peixoto et al. 2025), dropping donor Br9902 due to anatomy deformation (**Methods 4.24**). Relationships were tested at spatial scales of 100, 200, 500, 1,000, and 5,000 μm, values passed to makeShuffledCells(). makeShuffledCells() and findTrends() were run separately for each of the 8 samples. For visualization of results, we retained pairs of cell types where *Z*-scores agreed in sign across all samples and where significance (Bonferroni-adjusted p-value < 0.05) was achieved in at least 4 of 8 samples. To visualize dataset-wide statistics, *Z*-scores and spatial scales were averaged across samples. *Z*-score color scales were capped at twice the magnitude of the significance threshold returned by correctZBonferroni().

### 4.26 Cell to cell communication: *LIANA+*

We inferred spatially resolved cell-cell communication using *LIANA+* v1.7.1 (Dimitrov et al. 2024). Analyses were conducted separately for each tissue section, using cell-resolved VisiumHD spatial transcriptomic data with annotated cell type, with each cell type considered as a potential sender (source) and target population (target). We used the *LIANA+* consensus ligand-receptor resource, which integrates interactions from multiple expert-curated resources. For each tissue section, a distance-weighted spatial connectivity matrix was constructed from cell coordinates using a bandwidth of 5,000 μm (**Fig. S27A-B**), for connectivity between MHb and LHb. We then applied the *LIANA+* inflow method to quantify, for each target cell, spatially weighted ligand signals from neighboring cells of each source cell type in conjunction with receptor expression in the target cell. Cell-level inflow scores were summarized across target cell types using *LIANA+* global specificity analysis, yielding a mean inflow score (lr_mean) and a one-sided permutation-based P value for each source cell type, target cell type, and ligand-receptor pair. Within each sample, interactions with a nominal permutation P < 0.05 and a positive mean inflow score were considered significant. Interactions significant in at least four samples were retained (**Table S17**). Sample-level mean inflow scores were then averaged for each source-target-ligand-receptor combination. For visualization in **Fig. 6A**, sample-averaged interactions were further summarized across ligand-receptor pairs for each source-target cell-type pair.

Based on the overall communication patterns, we selected representative source-target cell-type pairs with relatively high communication signals for further characterization. For visualization of prioritized communications within these pairs, LR scores were first averaged within samples and then across samples, and interactions represented in at least four samples were retained. For each LR pair, relative enrichment was calculated as the mean interaction score for a given source-target cell-type pair divided by the mean score for the same LR pair across all reproducible mapped source-target cell-type combinations. Interactions with enrichment ≥3 were considered preferentially enriched. Among these enriched interactions, the top four LR pairs within each selected source-target cell-type pair were selected based on their mean interaction scores across samples (**Fig. 6D-E**).

We further performed Gene Ontology (GO) enrichment analysis using the unique ligand and receptor genes from LR interactions that were reproducibly detected in at least four samples and showed ≥3-fold enrichment, as described above. All genes represented in the same spatial transcriptomic dataset were used as the background gene universe. GO enrichment (*ClusterProfiler* v4.14.6 (Xu et al. 2024)) was performed separately for Biological Process (BP), Molecular Function (MF), and Cellular Component (CC). P values were adjusted using the Benjamini–Hochberg procedure, and GO terms with FDR < 0.05 were considered significant (**Table S18**). For visualization, the five most significant GO terms, ranked by FDR, were selected for each source-target cell-type pair within each ontology category. Unique GO terms contributed by these cell-type pairs were then combined within each comparison, with up to 30 terms with the lowest FDR displayed (**Fig. 6H-I**).

To assess whether cell-cell communication patterns were preferentially associated with psychiatric disease risk, we performed LR enrichment analyses using genes associated with major depressive disorder and substance dependence obtained from *OpenTargets* (Buniello et al. 2025). For each gene, we calculated the maximum *OpenTargets* genetic-association score across the available genetic evidence sources and retained genes with a score ≥0.1. Disease-associated genes were mapped to ligand and receptor roles using the OmniPath (Türei et al. 2016) intercellular interaction database, with HLA genes excluded. An LR pair was classified as disease-associated if either its ligand or receptor was an Open Targets disease-associated gene. *LIANA+*-derived LR interactions were restricted to those reproducibly detected in at least four samples. For each source-target cell-type pair, we tested whether disease-associated LR pairs were enriched relative to the universe of all unique LR pairs retained after the sample-reproducibility filter using a one-sided Fisher’s exact test. P values were adjusted across source-target cell-type pairs separately for each disease using the Benjamini-Hochberg FDR procedure (**Fig. 6B-C**).

### 4.27 Astrocyte differential expression

Briefly, we performed a differential expression analysis to find transcriptional differences between astrocytes located in the MHb vs those in the LHb. To prepare the cell-level data, we followed the process described in **Methods 4.23**, except we omitted the bin2cell.expand_labels() step, intentionally forming cells composed only of nuclei. We observed contamination of astrocyte cluster 11, located exclusively in the MHb, with MHb-specific transcriptional signal (**Fig. S15B**); this effect disappeared with the nuclear-only data. To classify astrocytes, we first only considered cells annotated (**Methods 4.23**) as MHb, LHb, or astrocytes; the dual *Excit.Thal/GABA_LHb_C.2* cluster was not considered habenula here. For each tissue section separately, we performed a KD-tree search with nn2(k = 10) with *RANN* v2.6.2 (Jefferis et al. 2013) to find each astrocyte’s 10 closest MHb and 10 closest LHb neighbors. If the mean of the distance to the MHb (LHb) neighbors was within 125 μm, the astrocyte was classified as MHb (LHb), else it was assigned the label “neither” (**Fig. 6F,G**). After assigning cell-level labels, the data were pseudobulked by tissue section and label. We checked differential expression of MHb and LHb astrocytes while modeling the number of cells and mitochondrial proportions as covariates. Briefly, this consisted of a workflow involving calcNormFactors(), filterByExpr.DGEList(), and voomLmFit(adaptive.span = TRUE, sample.weights = TRUE) (blocking by donor) from *edgeR* v4.6.2 (Chen et al. 2025), followed by eBayes() from *limma* v3.64.0 (Ritchie et al. 2015). To form a null background to empirically test the significance of the labeling approach we employed, astrocyte labels were shuffled randomly 1,000 times (between MHb, LHb, or neither), and an otherwise identical DE was conducted. We computed empirical p-values for each gene by checking the number of times the null-background *t*-statistics exceeded *t*-statistics from the properly labeled results in magnitude, ultimately correcting these values using the Benjamini-Hochberg procedure.

### 4.28 Enrichment of marker sets for risk for substance and psychiatric disorders with *MAGMA*

Gene-set association analyses were performed using *MAGMA* v1.10 (Leeuw et al. 2015) with GWAS summary statistics for European populations and psychiatric and substance use disorders. We used summary statistics for the following studies:

- *AUD_Zhou*, profiling problematic alcohol use (Zhou et al. 2023)

- *compulsive_F1_Grotzinger*, *SCZ/BPD_F2_Grotzinger*, *neurodev_F3_Grotzinger*, *internalizing_F4_Grotzinger*, *SUD_F5_Grotzinger*, and *p_factor_Grotzinger* (Grotzinger et al. 2026)

- *CUD_Johnson* (Johnson et al. 2020)

- externalizing_Linnér (Karlsson Linnér et al. 2021)

- *CUD_Pasman*, profiling lifetime cannabis use (Pasman et al. 2018)

- *MDD_Howard* (Howard et al. 2019)

- *OUD_Deak* (Deak et al. 2022)

- *panic_Forster* (Forstner et al. 2021)

- *SCZ_Trubetskoy* (Trubetskoy et al. 2022)

- *SUD_Hatoum* (Hatoum et al. 2023)

- *SUD_Polimanti* (Polimanti et al. 2020)

SNPs were mapped to genes using genomic coordinates from hg19, assigning variants located strictly within the genomic coordinates of transcriptional boundaries of each gene. Gene-level association statistics were calculated using the SNP-wise mean model, accounting for local linkage disequilibrium using the 1,000 Genomes European reference panel (The 1000 Genomes Project Consortium et al. 2015) (**Table S19**). We reported results for clusters whose gene sets contained >= 10 genes with computed statistics across all GWAS sets. Gene sets were *MeanRatio* markers computed among the datasets described in **Methods 4.11**. We then used competitive gene-set analysis to test whether genes within each set showed stronger gene-level associations with psychiatric and SUDs than genes outside the set; *MAGMA* internally adjusts statistics for gene size, gene density, and inverse minor allele count. Statistical significance was evaluated using two-sided competitive-test P values and corrected by cluster across all tested GWAS gene sets using the Benjamini-Hochberg procedure.

### 4.29 Enrichment of substance-use and psychiatric risk in differentially accessible regions using *S-LDSC*

We used stratified linkage disequilibrium score regression (*S-LDSC*) in *LDSC* v1.0.1 (Bulik-Sullivan et al. 2015) to test whether SNP-based heritability for a range of psychiatric and substance use disorders was enriched in genomic annotations representing habenula region- and cell type-specific chromatin accessibility. Binary SNP annotations were constructed separately for open, closed, and combined differential accessibility region (DAR) sets using hg19/GRCh37 genomic coordinates. SNPs overlapping a DAR in the corresponding annotation were assigned a value of 1, and all other SNPs were assigned a value of 0. Custom LD scores were calculated using European-ancestry reference genotypes from the 1000 Genomes Project Phase 3 (The 1000 Genomes Project Consortium et al. 2015) with a 1-cM LD window and were restricted to the same SNP set and order as the baseline-LD reference (Gazal et al. 2017). Each custom annotation was evaluated separately conditional on the baseline-LD v2.2 annotation set.

GWAS summary statistics were harmonized using munge_sumstats.py implemented in *LDSC* v1.0.1 (Bulik-Sullivan et al. 2015), restricted to *HapMap3* SNPs (Altshuler et al. 2010), and converted to *Z* statistics using available association and sample-size information. *HapMap3* regression weights excluding the HLA region and European-ancestry allele frequencies were used in the *S-LDSC* regression. Analyses were restricted to annotations containing more than 100 analyzed SNPs. For each annotation, we estimated the proportion of total SNP heritability explained and the corresponding enrichment, defined as the proportion of SNP heritability divided by the proportion of analyzed SNPs contained within the annotation. Results were summarized using *Z* statistics, with P values adjusted for multiple testing using the Benjamini–Hochberg false discovery rate (FDR) procedure within each cell type/state (**Fig. 7E**, **Table S20**).

### 4.30 Differentially Accessible Regions

Differentially accessible regions (DARs) were computed at the cell level at two cell-type resolutions (**Table S21**). At the fine resolution, all annotated snMultiome cell types were included individually. A broader resolution was also computed where all MHb cell types were grouped together, and all LHb were similarly grouped. At each of the 2 resolutions, FindMarkers(logfc.threshold = 1) from *Seurat* v5.3.0 (Stuart et al. 2019) was run to compute 1-vs-all comparisons on the normalized ATAC data (i.e. the data slot). We filtered results to just those with adjusted p-value < 0.2. For *S-LDSC* (**Methods 4.29**), we took just up to 1000 DARs for each of the open and closed directions, ranking by abs(avg_log2FC).

### 4.31 Fisher’s exact test; rat Hb bulk DEGs

We systematically tested whether our *MeanRatio* markers (**Methods 4.11**) were enriched for differentially expressed genes (FDR < 0.05) found in a reference rat habenula study (**Fig. 7F**) (Magnard et al. 2026). Each enrichment test used fisher.test(alternative = “greater”) using the intersection of this study’s post-QC genes and all genes considered for differential expression in the reference dataset as the universe. Tests operated with human ENSEMBL gene identifiers; starting with the reference rat DEGs, convert_orthologs(non121_strategy = “keep_popular”) from *orthogene* v1.14.01 (Schilder et al. 2026) was used to find human orthologs as gene symbols, followed by mapIds() from *AnnotationDbi* v1.70.0 (Hervé Pagès 2017) to convert symbols to ENSEMBL identifiers for comparison.

## 5 Statements

### 5.1 Data availability statement

Raw data (FASTQ files, images for Visium and VisiumHD) generated as part of this study have been deposited to the Neuroscience Multi-Omic Archive (NeMO) under collection <u>col-av55so6</u>. We also created interactive web applications to explore the data available at https://research.libd.org/Hb_multiome/#interactive-websites. Processed data files (R objects) used to make the interactive apps can be downloaded through the fetch_data() function from *spatialLIBD* v1.25.5 or newer (Pardo et al. 2022).

All source code for this project is available at https://github.com/LieberInstitute/Hb_multiome and https://github.com/LieberInstitute/Habenula_Visium under the MIT License and permanently archived on Zenodo (Cardinault et al. 2026; Eagles et al. 2026). The GitHub repositories include log files with specific software versions used for each analysis.

### 5.2 Acknowledgments

We gratefully thank the families who donated tissue to make this research possible. We thank the Essel Foundation and the families of Connie and Stephen Lieber and Milton and Tamar Maltz for their generous support of this work. We would like to acknowledge the contributions of Dr. Fernando Goes, the late Dr. Llewellyn B. Bigelow, Anna Brandtjen, and James Tooke for sample curation and clinical characterization. We also thank the Office of the Chief Medical Examiner of the State of Maryland for their collaboration on tissue collection. We thank Keri Martinowich and Robert Phillips for critical reading of the manuscript and feedback. We thank the Joint High Performance Computing Exchange (JHPCE) for providing computing resources for these analyses. We thank Hao Zhang and Joe Margolick from Flow Cytometry Cell Sorting Core Facility at Bloomberg School of Public Health, Johns Hopkins University (supported by 1S10OD016315-01,1S10RR13777001 and in part by CFAR: 5P30AI094189-04 (Chaisson). Portions of some figures were created with BioRender.com.

### 5.3 Author Contributions

Conceptualization: BM, LCT, KRM.

Data Curation: JMW, NJE, RAM.

Formal Analysis: KDM, JMW, NJE, CL, CSC.

Funding acquisition: BM, KRM.

Investigation: KDM, SVB, AC, RZ, SEM, HRD.

Project Administration: BM, LCT, KRM.

Resources: ADS, RAB, JEK, TMH.

Software: NJE, CL, LAHM.

Supervision: BM, LCT, KRM.

Visualization: KDM, JMW, NJE, CL, CSC, SVB.

Writing – original draft: KDM, JMW, NJE, CL, CSC, SVB, BM, LCT, KRM.

Writing – review & editing: KDM, JMW, NJE, CL, CSC, SVB, BM, LCT, KRM.

## References

Ables, Jessica L., Kwanghoon Park, and Inés Ibañez-Tallon. 2023. “Understanding the Habenula: A Major Node in Circuits Regulating Emotion and Motivation.” Pharmacological Research 190 (April): 106734. 10.1016/j.phrs.2023.106734.

Aihara, Gohta, Kalen Clifton, Mayling Chen, et al. 2024. “SEraster: A Rasterization Preprocessing Framework for Scalable Spatial Omics Data Analysis.” Bioinformatics 40 (7): btae412. 10.1093/bioinformatics/btae412.

Aizawa, Hidenori, Megumi Kobayashi, Sayaka Tanaka, Tomoki Fukai, and Hitoshi Okamoto. 2012. “Molecular Characterization of the Subnuclei in Rat Habenula.” The Journal of Comparative Neurology 520 (18): 4051–66. 10.1002/cne.23167.

Aizawa, Hidenori, Miho Matsumata, Laura Ayaka Noguera Oishi, et al. 2025. “Potassium Release From the Habenular Astrocytes Induces Depressive-Like Behaviors in Mice.” Glia 73 (4): 759–72. 10.1002/glia.24647.

Altshuler, David M., Richard A. Gibbs, Leena Peltonen, et al. 2010. “Integrating Common and Rare Genetic Variation in Diverse Human Populations.” Nature 467 (7311): 52–58. 10.1038/nature09298.

Andres, K. H., M. von Düring, and R. W. Veh. 1999. “Subnuclear Organization of the Rat Habenular Complexes.” The Journal of Comparative Neurology 407 (1): 130–50. 10.1002/(sici)1096-9861(19990428)407:1%3C130::aid-cne10%3E3.0.co;2-8.

Antolin-Fontes, Beatriz, Kun Li, Jessica L. Ables, et al. 2020. “The Habenular G-Protein–Coupled Receptor 151 Regulates Synaptic Plasticity and Nicotine Intake.” Proceedings of the National Academy of Sciences of the United States of America 117 (10): 5502–9. 10.1073/pnas.1916132117.

Arendt, Detlev, Jacob M. Musser, Clare V. H. Baker, et al. 2016. “The Origin and Evolution of Cell Types.” Nature Reviews Genetics 17 (12): 744–57. 10.1038/nrg.2016.127.

Bailly, Julie, Florence Allain, Eric Schwartz, et al. 2023. “Habenular Neurons Expressing Mu Opioid Receptors Promote Negative Affect in a Projection-Specific Manner.” Biological Psychiatry 93 (12): 1108–17. 10.1016/j.biopsych.2022.09.013.

Baker, Phillip M., Victor Mathis, Lucas Lecourtier, et al. 2022. “Lateral Habenula Beyond Avoidance: Roles in Stress, Memory, and Decision-Making With Implications for Psychiatric Disorders.” Frontiers in Systems Neuroscience 16 (March): 826475. 10.3389/fnsys.2022.826475.

Baldwin, Philip R., Raul Alanis, and Ramiro Salas. 2011. “The Role of the Habenula in Nicotine Addiction.” Journal of Addiction Research & Therapy Suppl 1 (2): 002. 10.4172/2155-6105.S1-002.

Baranasic, Damir. 2020. JASPAR2020: Data Package for JASPAR Database (Version 2020). V. 0.99.10. Released. http://jaspar.genereg.net/.

Blondel, Vincent D., Jean-Loup Guillaume, Renaud Lambiotte, and Etienne Lefebvre. 2008. “Fast Unfolding of Communities in Large Networks.” Journal of Statistical Mechanics: Theory and Experiment 2008 (10): P10008. 10.1088/1742-5468/2008/10/P10008.

Bosque-Cordero, Karl Y., Shikun Hou, and Elizabeth J. Glover. 2026. “Neuronal and Astrocytic Adaptations in the Lateral Habenula during Withdrawal from Chronic Ethanol.” Preprint, bioRxiv, August 10. 10.64898/2026.08.04.742855.

Bray, Natasha. 2018. “Bursting with Depression.” Nature Reviews Neuroscience 19 (4): 181–181. 10.1038/nrn.2018.28.

Bulik-Sullivan, Brendan K., Po-Ru Loh, Hilary K. Finucane, et al. 2015. “LD Score Regression Distinguishes Confounding from Polygenicity in Genome-Wide Association Studies.” Nature Genetics 47 (3): 291–95. 10.1038/ng.3211.

Buniello, Annalisa, Daniel Suveges, Carlos Cruz-Castillo, et al. 2025. “Open Targets Platform: Facilitating Therapeutic Hypotheses Building in Drug Discovery.” Nucleic Acids Research 53 (D1): D1467–75. 10.1093/nar/gkae1128.

Cardinault, Cynthia S., Nicholas J. Eagles, and Werner, Jonathan M. 2026. LieberInstitute/Hb_multiome: V0 - Preprint. V. v0-preprint. Zenodo, released September 28. 10.5281/ZENODO.23019505.

Chen, Yunshun, Lizhong Chen, Aaron T. L. Lun, Pedro L. Baldoni, and Gordon K. Smyth. 2025. “edgeR v4: Powerful Differential Analysis of Sequencing Data with Expanded Functionality and Improved Support for Small Counts and Larger Datasets.” Nucleic Acids Research 53 (2): gkaf018. 10.1093/nar/gkaf018.

Choi, Kyuhyun, Youngin Lee, Changwoo Lee, et al. 2016. “Optogenetic Activation of Septal GABAergic Afferents Entrains Neuronal Firing in the Medial Habenula.” Scientific Reports 6 (1): 34800. 10.1038/srep34800.

Ciscato, Maria, Mathilde Chouvaeff, and Alexandre Mourot. 2025. “Nicotinic Receptors in the Medial Habenula to Interpeduncular Nucleus Pathway: Modulators of Reward, Aversion and Emotion.” The European Journal of Neuroscience 62 (11): e70352. 10.1111/ejn.70352.

Civelli, Olivier. 2012. “Orphan GPCRs and Neuromodulation.” Neuron 76 (1): 12–21. 10.1016/j.neuron.2012.09.009.

Collado-Torres, Leonardo, Andrew E. Jaffe, and Emily E. Burke. 2024. Jaffelab: Commonly Used Functions by the Jaffe Lab. Released.

Crow, Megan, Anirban Paul, Sara Ballouz, Z. Josh Huang, and Jesse Gillis. 2018. “Characterizing the Replicability of Cell Types Defined by Single Cell RNA-Sequencing Data Using MetaNeighbor.” Nature Communications 9 (1): 884. 10.1038/s41467-018-03282-0.

Cui, Wanpeng, Hiroaki Mizukami, Michiko Yanagisawa, et al. 2014. “Glial Dysfunction in the Mouse Habenula Causes Depressive-Like Behaviors and Sleep Disturbance.” Articles. Journal of Neuroscience 34 (49): 16273–85. 10.1523/JNEUROSCI.1465-14.2014.

Cui, Yihui, Yan Yang, Zheyi Ni, et al. 2018. “Astroglial Kir4.1 in the Lateral Habenula Drives Neuronal Bursts in Depression.” Nature 554 (7692): 323–27. 10.1038/nature25752.

De Rop, Florian V., Gert Hulselmans, Chris Flerin, et al. 2024. “Systematic Benchmarking of Single-Cell ATAC-Sequencing Protocols.” Nature Biotechnology 42 (6): 916–26. 10.1038/s41587-023-01881-x.

Deak, Joseph D., Hang Zhou, Marco Galimberti, et al. 2022. “Genome-Wide Association Study in Individuals of European and African Ancestry and Multi-Trait Analysis of Opioid Use Disorder Identifies 19 Independent Genome-Wide Significant Risk Loci.” Molecular Psychiatry 27 (10): 3970–79. 10.1038/s41380-022-01709-1.

Díaz, Eugenia, Daniela Bravo, Ximena Rojas, and Miguel L. Concha. 2011. “Morphologic and Immunohistochemical Organization of the Human Habenular Complex.” The Journal of Comparative Neurology 519 (18): 3727–47. 10.1002/cne.22687.

Dimitrov, Daniel, Philipp Sven Lars Schäfer, Elias Farr, et al. 2024. “LIANA+ Provides an All-in-One Framework for Cell–Cell Communication Inference.” Nature Cell Biology 26 (9): 1613–22. 10.1038/s41556-024-01469-w.

Dos Santos Peixoto, Rafael, Brendan F. Miller, Maigan A. Brusko, et al. 2025. “Characterizing Cell-Type Spatial Relationships across Length Scales in Spatially Resolved Omics Data.” Nature Communications 16 (1): 350. 10.1038/s41467-024-55700-1.

Dyer, Sarah C., Olanrewaju Austine-Orimoloye, Andrey G. Azov, et al. 2025. “Ensembl 2025.” Nucleic Acids Research 53 (D1): D948–57. 10.1093/nar/gkae1071.

Eagles, Nicholas J., Cynthia S. Cardinault, Leonardo Collado-Torres, et al. 2026. LieberInstitute/Habenula_Visium: V0 - Preprint. V. v0-preprint. Zenodo, released September. 10.5281/zenodo.23019474.

Fischer, Stephan, Megan Crow, Benjamin D. Harris, and Jesse Gillis. 2021. “Scaling up Reproducible Research for Single Cell Transcriptomics Using MetaNeighbor.” Nature Protocols 16 (8): 4031–67. 10.1038/s41596-021-00575-5.

Fischer, Stephan, and Jesse Gillis. 2021. “How Many Markers Are Needed to Robustly Determine a Cell’s Type?” iScience 24 (11): 103292. 10.1016/j.isci.2021.103292.

Flanigan, Meghan E., Hossein Aleyasin, Long Li, et al. 2020. “Orexin Signaling in GABAergic Lateral Habenula Neurons Modulates Aggressive Behavior in Male Mice.” Nature Neuroscience 23 (5): 638–50. 10.1038/s41593-020-0617-7.

Forstner, Andreas J., Swapnil Awasthi, Christiane Wolf, et al. 2021. “Genome-Wide Association Study of Panic Disorder Reveals Genetic Overlap with Neuroticism and Depression.” Molecular Psychiatry 26 (8): 4179–90. 10.1038/s41380-019-0590-2.

Fortin, Jean-Simon, Mathilde Lafleur, Charles Parisien, and Sébastien Hétu. 2025. “The Habenula in Mood Disorders: A Systematic Review of Human Studies.” Molecular Psychiatry 30 (10): 4948–70. 10.1038/s41380-025-03105-x.

Frankish, Adam, Sílvia Carbonell-Sala, Mark Diekhans, et al. 2023. “GENCODE: Reference Annotation for the Human and Mouse Genomes in 2023.” Nucleic Acids Research 51 (D1): D942–49. 10.1093/nar/gkac1071.

Frankish, Adam, Mark Diekhans, Anne-Maud Ferreira, et al. 2019. “GENCODE Reference Annotation for the Human and Mouse Genomes.” Nucleic Acids Research 47 (D1): D766–73. 10.1093/nar/gky955.

Gampe, Kristine, Klaus Hammer, Ágnes Kittel, and Herbert Zimmermann. 2012. “The Medial Habenula Contains a Specific Nonstellate Subtype of Astrocyte Expressing the Ectonucleotidase NTPDase2.” Glia 60 (12): 1860–70. 10.1002/glia.22402.

García-Durán, Laura, Antonio Flores-Burgess, Noelia Cantero-García, et al. 2021. “Galanin(1-15) Potentiates the Antidepressant-like Effects Induced by Escitalopram in a Rat Model of Depression.” International Journal of Molecular Sciences 22 (19): 10848. 10.3390/ijms221910848.

Gardon, O., L. Faget, P. Chu Sin Chung, A. Matifas, D. Massotte, and B. L. Kieffer. 2014. “Expression of Mu Opioid Receptor in Dorsal Diencephalic Conduction System: New Insights for the Medial Habenula.” Neuroscience 277 (September): 595–609. 10.1016/j.neuroscience.2014.07.053.

Gaspar, John M. 2018. “Improved Peak-Calling with MACS2.” Preprint, Bioinformatics, December 17. 10.1101/496521.

Gazal, Steven, Hilary K. Finucane, Nicholas A. Furlotte, et al. 2017. “Linkage Disequilibrium–Dependent Architecture of Human Complex Traits Shows Action of Negative Selection.” Nature Genetics 49 (10): 1421–27. 10.1038/ng.3954.

Geisler, Stefanie, Karl Hermann Andres, and Rüdiger W. Veh. 2003. “Morphologic and cytochemical criteria for the identification and delineation of individual subnuclei within the lateral habenular complex of the rat.” Journal of Comparative Neurology 458 (1): 78–97. 10.1002/cne.10566.

Germain, Pierre-Luc, Aaron Lun, Carlos Garcia Meixide, Will Macnair, and Mark D. Robinson. 2022. “Doublet Identification in Single-Cell Sequencing Data Using scDblFinder.” Preprint, F1000Research, May 16. 10.12688/f1000research.73600.2.

Green, Matthew V., David A. Gallegos, Jane-Valeriane Boua, Luke C. Bartelt, Arthy Narayanan, and Anne E. West. 2023. “Single-Nucleus Transcriptional Profiling of GAD2-Positive Neurons From Mouse Lateral Habenula Reveals Distinct Expression of Neurotransmission- and Depression-Related Genes.” Biological Psychiatry Global Open Science 3 (4): 686–97. 10.1016/j.bpsgos.2023.04.004.

Grotzinger, Andrew D., Josefin Werme, Wouter J. Peyrot, et al. 2026. “Mapping the Genetic Landscape across 14 Psychiatric Disorders.” Nature 649 (8096): 406–15. 10.1038/s41586-025-09820-3.

Gustavsen, Julia A., Shraddha Pai, Ruth Isserlin, Barry Demchak, and Alexander R. Pico. 2019. “RCy3: Network Biology Using Cytoscape from within R.” F1000Research 8 (December): 1774. 10.12688/f1000research.20887.3.

Hahn, Joshua, Aboozar Monavarfeshani, Mu Qiao, et al. 2023. “Evolution of Neuronal Cell Classes and Types in the Vertebrate Retina.” Nature 624 (7991): 415–24. 10.1038/s41586-023-06638-9.

Hahsler, Michael, Matthew Piekenbrock, and Derek Doran. 2019. “**Dbscan** : Fast Density-Based Clustering with R.” Journal of Statistical Software 91 (1). 10.18637/jss.v091.i01.

Hao, Yuhan, Tim Stuart, Madeline H. Kowalski, et al. 2024. “Dictionary Learning for Integrative, Multimodal and Scalable Single-Cell Analysis.” Nature Biotechnology 42 (2): 293–304. 10.1038/s41587-023-01767-y.

Hashikawa, Yoshiko, Koichi Hashikawa, Mark A. Rossi, et al. 2020. “Transcriptional and Spatial Resolution of Cell Types in the Mammalian Habenula.” Neuron 106 (5): 743–758.e5. 10.1016/j.neuron.2020.03.011.

Hatoum, Alexander S., Sarah M. C. Colbert, Emma C. Johnson, et al. 2023. “Multivariate Genome-Wide Association Meta-Analysis of over 1 Million Subjects Identifies Loci Underlying Multiple Substance Use Disorders.” Nature Mental Health 1 (3): 210–23. 10.1038/s44220-023-00034-y.

Herkenham, M., and W. J. Nauta. 1979. “Efferent Connections of the Habenular Nuclei in the Rat.” The Journal of Comparative Neurology 187 (1): 19–47. 10.1002/cne.901870103.

Hervé Pagès, Marc Carlson. 2017. AnnotationDbi. Bioconductor, released. 10.18129/B9.BIOC.ANNOTATIONDBI.

Heumos, Lukas, Anna C. Schaar, Christopher Lance, et al. 2023. “Best Practices for Single-Cell Analysis across Modalities.” Nature Reviews Genetics 24 (8): 550–72. 10.1038/s41576-023-00586-w.

Howard, David M., Mark J. Adams, Toni-Kim Clarke, et al. 2019. “Genome-Wide Meta-Analysis of Depression Identifies 102 Independent Variants and Highlights the Importance of the Prefrontal Brain Regions.” Nature Neuroscience 22 (3): 343–52. 10.1038/s41593-018-0326-7.

Hu, Hailan, Yihui Cui, and Yan Yang. 2020. “Circuits and Functions of the Lateral Habenula in Health and in Disease.” Nature Reviews Neuroscience 21 (5): 277–95. 10.1038/s41583-020-0292-4.

Huuki-Myers, Louise A., Heena R. Divecha, Svitlana V. Bach, et al. 2025. “APOE E4 Alzheimer’s Risk Converges on an Oligodendrocyte Subtype in the Human Entorhinal Cortex.” Preprint, bioRxiv, November 20. 10.1101/2025.11.20.689483.

Huuki-Myers, Louise A., Kelsey D. Montgomery, Sang Ho Kwon, et al. 2025. “Benchmark of Cellular Deconvolution Methods Using a Multi-Assay Dataset from Postmortem Human Prefrontal Cortex.” Genome Biology 26 (1): 88. 10.1186/s13059-025-03552-3.

Huuki-Myers, Louise A., Abby Spangler, Nicholas J. Eagles, et al. 2024. “A Data-Driven Single-Cell and Spatial Transcriptomic Map of the Human Prefrontal Cortex.” Science 384 (6698): eadh1938. 10.1126/science.adh1938.

Improgo, Ma. Reina D., Michael D. Scofield, Andrew R. Tapper, and Paul D. Gardner. 2010. “The Nicotinic Acetylcholine Receptor CHRNA5/A3/B4 Gene Cluster: Dual Role in Nicotine Addiction and Lung Cancer.” Progress in Neurobiology 92 (2): 212–26. 10.1016/j.pneurobio.2010.05.003.

Jacinto, Luis R., Rui Mata, Ashley Novais, Fernanda Marques, and Nuno Sousa. 2017. “The Habenula as a Critical Node in Chronic Stress-Related Anxiety.” Experimental Neurology 289 (March): 46–54. 10.1016/j.expneurol.2016.12.003.

Jefferis, Gregory, Samuel E. Kemp, Sunil Arya, and David Mount. 2013. “RANN: Fast Nearest Neighbour Search (Wraps ANN Library) Using L2 Metric.” July 24. 10.32614/CRAN.package.RANN.

Jiang, Yuchao, Yuriko Harigaya, Zhaojun Zhang, Hongpan Zhang, Chongzhi Zang, and Nancy R. Zhang. 2022. “Nonparametric Single-Cell Multiomic Characterization of Trio Relationships between Transcription Factors, Target Genes, and Cis-Regulatory Regions.” Cell Systems 13 (9): 737–751.e4. 10.1016/j.cels.2022.08.004.

Johnson, Emma C., Ditte Demontis, Thorgeir E. Thorgeirsson, et al. 2020. “A Large-Scale Genome-Wide Association Study Meta-Analysis of Cannabis Use Disorder.” The Lancet Psychiatry 7 (12): 1032–45. 10.1016/S2215-0366(20)30339-4.

Kahle, Kristopher T., Tarek Z. Deeb, Martin Puskarjov, et al. 2013. “Modulation of Neuronal Activity by Phosphorylation of the K–Cl Cotransporter KCC2.” Trends in Neurosciences 36 (12): 726–37. 10.1016/j.tins.2013.08.006.

Karlsson Linnér, Richard, Travis T. Mallard, Peter B. Barr, et al. 2021. “Multivariate Analysis of 1.5 Million People Identifies Genetic Associations with Traits Related to Self-Regulation and Addiction.” Nature Neuroscience 24 (10): 1367–76. 10.1038/s41593-021-00908-3.

Kawai, Takafumi, Ping Dong, Konstantin Bakhurin, Henry H. Yin, and Huanghe Yang. 2025. “Calcium-Activated Ion Channels Drive Atypical Inhibition in Medial Habenula Neurons.” Science Advances 11 (12): eadq2629. 10.1126/sciadv.adq2629.

Kim, Kristen, and Marina R. Picciotto. 2023. “Nicotine Addiction: More than Just Dopamine.” Current Opinion in Neurobiology 83 (December): 102797. 10.1016/j.conb.2023.102797.

Kim, Uhnoh, and Lee-yup Chung. 2007. “Dual GABAergic Synaptic Response of Fast Excitation and Slow Inhibition in the Medial Habenula of Rat Epithalamus.” Journal of Neurophysiology 98 (3): 1323–32. 10.1152/jn.00575.2007.

Kononoff, Jenni, Marsida Kallupi, Adam Kimbrough, Dana Conlisk, Giordano de Guglielmo, and Olivier George. 2018. “Systemic and Intra-Habenular Activation of the Orphan G Protein-Coupled Receptor GPR139 Decreases Compulsive-Like Alcohol Drinking and Hyperalgesia in Alcohol-Dependent Rats.” New Research. eNeuro 5 (3). 10.1523/ENEURO.0153-18.2018.

Korsunsky, Ilya, Nghia Millard, Jean Fan, et al. 2019. “Fast, Sensitive and Accurate Integration of Single-Cell Data with Harmony.” Nature Methods 16 (12): 1289–96. 10.1038/s41592-019-0619-0.

Laboute, Thibaut, Stefano Zucca, Matthew Holcomb, et al. 2023. “Orphan Receptor GPR158 Serves as a Metabotropic Glycine Receptor: mGlyR.” Science 379 (6639): 1352–58. 10.1126/science.add7150.

Lawson, R. P., C. L. Nord, B. Seymour, et al. 2017. “Disrupted Habenula Function in Major Depression.” Molecular Psychiatry 22 (2): 202–8. 10.1038/mp.2016.81.

Lawson, Rebecca P., Wayne C. Drevets, and Jonathan P. Roiser. 2013. “Defining the Habenula in Human Neuroimaging Studies.” Neuroimage 64 (January): 722–27. 10.1016/j.neuroimage.2012.08.076.

Leeuw, Christiaan A. de, Joris M. Mooij, Tom Heskes, and Danielle Posthuma. 2015. “MAGMA: Generalized Gene-Set Analysis of GWAS Data.” PLOS Computational Biology 11 (4): e1004219. 10.1371/journal.pcbi.1004219.

Li, Bo, Joaquin Piriz, Martine Mirrione, et al. 2011. “Synaptic Potentiation onto Habenula Neurons in the Learned Helplessness Model of Depression.” Nature 470 (7335): 535–39. 10.1038/nature09742.

Liang, Tingming, Li Guo, and Chang Liu. 2012. “Genome-Wide Analysis of Mir-548 Gene Family Reveals Evolutionary and Functional Implications.” Journal of Biomedicine and Biotechnology 2012: 679563. 10.1155/2012/679563.

Lin, Feiteng, Kayleigh Casmey, Sierra A. Codeluppi-Arrowsmith, and Gustavo Turecki. 2026. “The Habenula’s Role in Major Depressive Disorder: Recent Insights from Preclinical and Human Studies.” Translational Psychiatry 16 (1): 77. 10.1038/s41398-026-03867-0.

Lin, Jinju, Qin Li, Xiaohua Lei, and Huashan Zhao. 2022. “The Emerging Roles of GPR158 in the Regulation of the Endocrine System.” Frontiers in Cell and Developmental Biology 10 (November): 1034348. 10.3389/fcell.2022.1034348.

Lipska, Barbara K., Amy Deep-Soboslay, Cynthia Shannon Weickert, et al. 2006. “Critical Factors in Gene Expression in Postmortem Human Brain: Focus on Studies in Schizophrenia.” Biological Psychiatry 60 (6): 650–58. 10.1016/j.biopsych.2006.06.019.

Lun, Aaron T. L., Davis J. McCarthy, and John C. Marioni. 2016. “A Step-by-Step Workflow for Low-Level Analysis of Single-Cell RNA-Seq Data with Bioconductor.” F1000Research 5 (October): 2122. 10.12688/f1000research.9501.2.

Magnard, Robin, Daianna Gonzalez-Padilla, Ege A. Yalcinbas, et al. 2026. “Transcriptional Response to Chronic Long-Access Fentanyl Self-Administration in Rat Habenula and Amygdala.” Addiction Biology 31 (7): e70179. 10.1111/adb.70179.

Marburg, Otto. 1944. The Structure and Fiber Connections of the Human Habenula. 80: 211–33. 10.1002/cne.900800205.

Matsumoto, Masayuki, and Okihide Hikosaka. 2007. “Lateral Habenula as a Source of Negative Reward Signals in Dopamine Neurons.” Nature 447 (7148): 1111–15. 10.1038/nature05860.

Maynard, Kristen R., Leonardo Collado-Torres, Lukas M. Weber, et al. 2021. “Transcriptome-Scale Spatial Gene Expression in the Human Dorsolateral Prefrontal Cortex.” Nature Neuroscience 24 (3): 425–36. 10.1038/s41593-020-00787-0.

Maynard, Kristen R., Madhavi Tippani, Yoichiro Takahashi, et al. 2020. “Dotdotdot: An Automated Approach to Quantify Multiplex Single Molecule Fluorescent in Situ Hybridization (smFISH) Images in Complex Tissues.” Nucleic Acids Research 48 (11): e66. 10.1093/nar/gkaa312.

McCarthy, Davis J., Kieran R. Campbell, Aaron T. L. Lun, and Quin F. Wills. 2017. “Scater: Pre-Processing, Quality Control, Normalization and Visualization of Single-Cell RNA-Seq Data in R.” Bioinformatics 33 (8): 1179–86. 10.1093/bioinformatics/btw777.

McInnes, Leland, John Healy, and James Melville. 2020. “UMAP: Uniform Manifold Approximation and Projection for Dimension Reduction.” arXiv:1802.03426. Preprint, arXiv, September 18. 10.48550/arXiv.1802.03426.

McMoneagle, Erin, Jin Zhou, Shiyao Zhang, et al. 2024. “Neuronal K+-Cl-Cotransporter KCC2 as a Promising Drug Target for Epilepsy Treatment.” Acta Pharmacologica Sinica 45 (1): 1–22. 10.1038/s41401-023-01149-9.

Milanesi, Elena, Cristian Bonvicini, Chiara Congiu, et al. 2015. “The Role of GRIK4 Gene in Treatment-Resistant Depression.” Genetics Research 97 (July): e14. 10.1017/S0016672315000142.

Müller, Ulf J., Moritz Ahrens, Veronika Vasilevska, et al. 2021. “Reduced Habenular Volumes and Neuron Numbers in Male Heroin Addicts: A Post-Mortem Study.” European Archives of Psychiatry and Clinical Neuroscience 271 (5): 835–45. 10.1007/s00406-020-01195-y.

Nakamura, Tomoya, Trang Van Thi Nguyen, Takumi Maeda, and Hiroyuki Ichijo. 2024. “Substantial Projections from the Lateral Division of the Lateral Habenula to the Dorsal Raphe Nucleus and from the Lateral Habenula to the Contralateral Ventral Tegmental Area.” Heliyon 10 (22): e40234. 10.1016/j.heliyon.2024.e40234.

Namboodiri, Vijay Mohan K., Jose Rodriguez-Romaguera, and Garret D. Stuber. 2016. “The Habenula.” Current Biology 26 (19): R873–77. 10.1016/j.cub.2016.08.051.

Neel, Dylan V., Himanish Basu, Georgia Gunner, et al. 2023. “Gasdermin-E Mediates Mitochondrial Damage in Axons and Neurodegeneration.” Neuron 111 (8): 1222–1240.e9. 10.1016/j.neuron.2023.02.019.

Nguyen, Thi Van Trang, Tomoya Nakamura, and Hiroyuki Ichijo. 2024. “Topographic Organization of Glutamatergic and GABAergic Parvalbumin-Positive Neurons in the Lateral Habenula.” eNeuro 11 (7): ENEURO.0069-24.2024. 10.1523/ENEURO.0069-24.2024.

Notomi, Takuya, and Ryuichi Shigemoto. 2004. “Immunohistochemical Localization of Ih Channel Subunits, HCN1-4, in the Rat Brain.” The Journal of Comparative Neurology 471 (3): 241–76. 10.1002/cne.11039.

Paddock, Silvia, Gonzalo Laje, Dennis Charney, et al. 2007. “Association of GRIK4 with Outcome of Antidepressant Treatment in the STAR*D Cohort.” The American Journal of Psychiatry 164 (8): 1181–88. 10.1176/appi.ajp.2007.06111790.

Pandey, Shristi, Karthik Shekhar, Aviv Regev, and Alexander F. Schier. 2018. “Comprehensive Identification and Spatial Mapping of Habenular Neuronal Types Using Single-Cell RNA-Seq.” Current Biology 28 (7): 1052–1065.e7. 10.1016/j.cub.2018.02.040.

Pang, Xueyan, Liwang Liu, Jennifer Ngolab, et al. 2016. “Habenula Cholinergic Neurons Regulate Anxiety during Nicotine Withdrawal via Nicotinic Acetylcholine Receptors.” Neuropharmacology 107 (August): 294–304. 10.1016/j.neuropharm.2016.03.039.

Pardo, Brenda, Abby Spangler, Lukas M. Weber, et al. 2022. “spatialLIBD: An R/Bioconductor Package to Visualize Spatially-Resolved Transcriptomics Data.” BMC Genomics 23 (1): 434. 10.1186/s12864-022-08601-w.

Pasman, Joëlle A., the 23andMe Research Team, The Substance Use Disorders Working Group of the Psychiatric Genomics Consortium, et al. 2018. “GWAS of Lifetime Cannabis Use Reveals New Risk Loci, Genetic Overlap with Psychiatric Traits, and a Causal Effect of Schizophrenia Liability.” Nature Neuroscience 21 (9): 1161–70. 10.1038/s41593-018-0206-1.

Polański, Krzysztof, Raquel Bartolomé-Casado, Ioannis Sarropoulos, et al. 2024. “Bin2cell Reconstructs Cells from High Resolution Visium HD Data.” Bioinformatics 40 (9): btae546. 10.1093/bioinformatics/btae546.

Polimanti, Renato, Raymond K. Walters, Emma C. Johnson, et al. 2020. “Leveraging Genome-Wide Data to Investigate Differences between Opioid Use vs. Opioid Dependence in 41,176 Individuals from the Psychiatric Genomics Consortium.” Molecular Psychiatry 25 (8): 1673–87. 10.1038/s41380-020-0677-9.

Poller, W. C., R. Bernard, C. Derst, T. Weiss, V. I. Madai, and R. W. Veh. 2011. “Lateral Habenular Neurons Projecting to Reward-Processing Monoaminergic Nuclei Express Hyperpolarization-Activated Cyclic Nucleotid-Gated Cation Channels.” Neuroscience 193 (October): 205–16. 10.1016/j.neuroscience.2011.07.013.

Popat, Aashka K., Rhiana C. Simon, Beatriz B. Aoyama, et al. 2026. “Mu Opioid Receptor mRNA and Protein Localization Across the Rat and Mouse Habenula.” The Journal of Comparative Neurology 534 (6): e70175. 10.1002/cne.70175.

Quina, Lely A., Andrew Walker, Glenn Morton, Victor Han, and Eric E. Turner. 2020. “GAD2 Expression Defines a Class of Excitatory Lateral Habenula Neurons in Mice That Project to the Raphe and Pontine Tegmentum.” Research Article: New Research. eNeuro 7 (3). 10.1523/ENEURO.0527-19.2020.

Righelli, Dario, Lukas M. Weber, Helena L. Crowell, et al. 2022. “SpatialExperiment: Infrastructure for Spatially-Resolved Transcriptomics Data in R Using Bioconductor.” Bioinformatics 38 (11): 3128–31. 10.1093/bioinformatics/btac299.

Ritchie, Matthew E., Belinda Phipson, Di Wu, et al. 2015. “Limma Powers Differential Expression Analyses for RNA-Sequencing and Microarray Studies.” Nucleic Acids Research 43 (7): e47–e47. 10.1093/nar/gkv007.

Roberson, Sara, and Marnie E. Halpern. 2018. “Development and Connectivity of the Habenular Nuclei.” Seminars in Cell & Developmental Biology 78 (June): 107–15. 10.1016/j.semcdb.2017.10.007.

Rotta, Randolf, and Andreas Noack. 2011. “Multilevel Local Search Algorithms for Modularity Clustering.” Journal of Experimental Algorithmics (JEA) 16 (July): 2.3:2.1-2.3:2.27. 10.1145/1963190.1970376.

Roy, Nisa, and Ishwar Parhar. 2022. “Habenula Orphan G-Protein Coupled Receptors in the Pathophysiology of Fear and Anxiety.” Neuroscience & Biobehavioral Reviews 132 (January): 870–83. 10.1016/j.neubiorev.2021.11.008.

Santos Peixoto, Rafael dos, Brendan F. Miller, Maigan A. Brusko, et al. 2025. “Characterizing Cell-Type Spatial Relationships across Length Scales in Spatially Resolved Omics Data.” Nature Communications 16 (1): 350. 10.1038/s41467-024-55700-1.

Savitz, Jonathan B., Allison C. Nugent, Wendy Bogers, et al. 2011. “Habenula Volume in Bipolar Disorder and Major Depressive Disorder: A High Resolution MRI Study.” Biological Psychiatry 69 (4): 336–43. 10.1016/j.biopsych.2010.09.027.

Schep, Alicia N., Beijing Wu, Jason D. Buenrostro, and William J. Greenleaf. 2017. “chromVAR: Inferring Transcription-Factor-Associated Accessibility from Single-Cell Epigenomic Data.” Nature Methods 14 (10): 975–78. 10.1038/nmeth.4401.

Schilder, Brian M., Alan E. Murphy, and Nathan G. Skene. 2026. “Orthogene: A Bioconductor Package to Easily Map Genes within and across Hundreds of Species.” Preprint, bioRxiv, January 21. 10.64898/2026.01.17.700094.

Shafer, Maxwell E. R. 2019. “Cross-Species Analysis of Single-Cell Transcriptomic Data.” Frontiers in Cell and Developmental Biology 7 (September): 175. 10.3389/fcell.2019.00175.

Shannon, Paul, Andrew Markiel, Owen Ozier, et al. 2003. “Cytoscape: A Software Environment for Integrated Models of Biomolecular Interaction Networks.” Genome Research 13 (11): 2498–504. 10.1101/gr.1239303.

Si, Yichen, ChangHee Lee, Yongha Hwang, et al. 2024. “FICTURE: Scalable Segmentation-Free Analysis of Submicron-Resolution Spatial Transcriptomics.” Nature Methods 21 (10): 1843–54. 10.1038/s41592-024-02415-2.

Singhal, Vipul, Nigel Chou, Joseph Lee, et al. 2024. “BANKSY Unifies Cell Typing and Tissue Domain Segmentation for Scalable Spatial Omics Data Analysis.” Nature Genetics 56 (3): 431–41. 10.1038/s41588-024-01664-3.

Song, Sukwoon, Miseon Kang, Jiyoung Lee, et al. 2025. “Role of Phospholipase Cη1 in Lateral Habenula Astrocytes in Depressive-like Behavior in Mice.” Experimental & Molecular Medicine 57 (4): 872–87. 10.1038/s12276-025-01432-1.

Sriram, Krishna, and Paul A. Insel. 2018. “G Protein-Coupled Receptors as Targets for Approved Drugs: How Many Targets and How Many Drugs?” Molecular Pharmacology 93 (4): 251–58. 10.1124/mol.117.111062.

Stamatakis, Alice M., and Garret D. Stuber. 2012. “Activation of Lateral Habenula Inputs to the Ventral Midbrain Promotes Behavioral Avoidance.” Nature Neuroscience 15 (8): 1105–7. 10.1038/nn.3145.

Stuart, Tim, Andrew Butler, Paul Hoffman, et al. 2019. “Comprehensive Integration of Single-Cell Data.” Cell 177 (7): 1888–1902.e21. 10.1016/j.cell.2019.05.031.

Stuart, Tim, Avi Srivastava, Shaista Madad, Caleb A. Lareau, and Rahul Satija. 2021. “Single-Cell Chromatin State Analysis with Signac.” Nature Methods 18 (11): 1333–41. 10.1038/s41592-021-01282-5.

Tan, Wanqin, Yoko Ikoma, Yusuke Takahashi, et al. 2024. “Anxiety Control by Astrocytes in the Lateral Habenula.” Neuroscience Research 205 (August): 1–15. 10.1016/j.neures.2024.01.006.

Tang, Xin, Julie Kim, Li Zhou, et al. 2016. “KCC2 Rescues Functional Deficits in Human Neurons Derived from Patients with Rett Syndrome.” Proceedings of the National Academy of Sciences 113 (3): 751–56. 10.1073/pnas.1524013113.

The 1000 Genomes Project Consortium, Corresponding authors, Adam Auton, et al. 2015. “A Global Reference for Human Genetic Variation.” Nature 526 (7571): 68–74. 10.1038/nature15393.

Tian, Jing, Ziran Huang, and Wen Zhang. 2025. “Gpr176 Modulates the Firing Pattern of Parvalbumin-Positive Interneurons in the Orbitofrontal Cortex of Mouse.” Molecular Brain 18 (November): 81. 10.1186/s13041-025-01254-2.

Tippani, Madhavi, Heena R. Divecha, Joseph L. Catallini, et al. 2023. “VistoSeg : Processing Utilities for High-Resolution Images for Spatially Resolved Transcriptomics Data.” Biological Imaging 3: e23. 10.1017/S2633903X23000235.

Totty, Michael, Stephanie C. Hicks, and Boyi Guo. 2025. “SpotSweeper: Spatially Aware Quality Control for Spatial Transcriptomics.” Nature Methods 22 (7): 1520–30. 10.1038/s41592-025-02713-3.

Traag, V. A., L. Waltman, and N. J. van Eck. 2019. “From Louvain to Leiden: Guaranteeing Well-Connected Communities.” Scientific Reports 9 (1): 5233. 10.1038/s41598-019-41695-z.

Trubetskoy, Vassily, Antonio F. Pardiñas, Ting Qi, et al. 2022. “Mapping Genomic Loci Implicates Genes and Synaptic Biology in Schizophrenia.” Nature 604 (7906): 502–8. 10.1038/s41586-022-04434-5.

Türei, Dénes, Tamás Korcsmáros, and Julio Saez-Rodriguez. 2016. “OmniPath: Guidelines and Gateway for Literature-Curated Signaling Pathway Resources.” Nature Methods 13 (12): 966–67. 10.1038/nmeth.4077.

Velasquez, Kenia M., David L. Molfese, and Ramiro Salas. 2014. “The Role of the Habenula in Drug Addiction.” Frontiers in Human Neuroscience 8 (March): 174. 10.3389/fnhum.2014.00174.

Wagner, Franziska, René Bernard, Christian Derst, Leon French, and Rüdiger W. Veh. 2016. “Microarray Analysis of Transcripts with Elevated Expressions in the Rat Medial or Lateral Habenula Suggest Fast GABAergic Excitation in the Medial Habenula and Habenular Involvement in the Regulation of Feeding and Energy Balance.” Brain Structure and Function 221 (9): 4663–89. 10.1007/s00429-016-1195-z.

Wagner, Franziska, Torsten Weiss, and Rüdiger W. Veh. 2017. “Electrophysiological Properties of Neurons and Synapses in the Lateral Habenular Complex (LHb).” Pharmacology Biochemistry and Behavior, The lateral habenula. From the neuroanatomy to the implication in CNS disorders, vol. 162 (November): 38–45. 10.1016/j.pbb.2017.07.008.

Wallace, Michael L., Kee Wui Huang, Daniel Hochbaum, Minsuk Hyun, Gianna Radeljic, and Bernardo L. Sabatini. 2020. “Anatomical and Single-Cell Transcriptional Profiling of the Murine Habenular Complex.” eLife 9 (February): e51271. 10.7554/eLife.51271.

Waltman, Ludo, and Nees Jan van Eck. 2013. “A Smart Local Moving Algorithm for Large-Scale Modularity-Based Community Detection.” The European Physical Journal B 86 (11): 471. 10.1140/epjb/e2013-40829-0.

Wang, D. G., N. Gong, B. Luo, and T. L. Xu. 2006. “Absence of GABA Type A Signaling in Adult Medial Habenular Neurons.” Neuroscience 141 (1): 133–41. 10.1016/j.neuroscience.2006.03.045.

Wang, Zhiyan, Chao Jiang, Lingxiao Guan, et al. 2024. “Deep Brain Stimulation of Habenula Reduces Depressive Symptoms and Modulates Brain Activities in Treatment-Resistant Depression.” Nature Mental Health 2 (9): 1045–52. 10.1038/s44220-024-00286-2.

Weber, Lukas M., Arkajyoti Saha, Abhirup Datta, Kasper D. Hansen, and Stephanie C. Hicks. 2023. “nnSVG for the Scalable Identification of Spatially Variable Genes Using Nearest-Neighbor Gaussian Processes.” Nature Communications 14 (1): 4059. 10.1038/s41467-023-39748-z.

Wolfe, Clara I. C., Eun-Kyung Hwang, Elfrieda C. Ijomor, Agustin Zapata, Alexander F. Hoffman, and Carl R. Lupica. 2022. “Muscarinic Acetylcholine M2 Receptors Regulate Lateral Habenula Neuron Activity and Control Cocaine Seeking Behavior.” Research Articles. Journal of Neuroscience 42 (28): 5552–63. 10.1523/JNEUROSCI.0645-22.2022.

Wyler, Steven, Surbhi, Newton Cao, Warda Merchant, Angie Bookout, and Laurent Gautron. 2024. “Gpr149 Is Involved in Energy Homeostasis in the Male Mouse.” PeerJ 12 (January): e16739. 10.7717/peerj.16739.

Xin, Qianqian, Junying Wang, Jinkun Zheng, et al. 2025. “Neuron-Astrocyte Coupling in Lateral Habenula Mediates Depressive-like Behaviors.” Cell 188 (12): 3291–3309.e24. 10.1016/j.cell.2025.04.010.

Xu, Shuangbin, Erqiang Hu, Yantong Cai, et al. 2024. “Using clusterProfiler to Characterize Multiomics Data.” Nature Protocols 19 (11): 3292–320. 10.1038/s41596-024-01020-z.

Yalcinbas, Ege A., Bukola Ajanaku, Erik D. Nelson, et al. 2025. “Transcriptomic Analysis of the Human Habenula in Schizophrenia.” American Journal of Psychiatry 182 (11): 991–1006. 10.1176/appi.ajp.20240776.

Yang, Yan, Yihui Cui, Kangning Sang, et al. 2018. “Ketamine Blocks Bursting in the Lateral Habenula to Rapidly Relieve Depression.” Nature 554 (7692): 317–22. 10.1038/nature25509.

Yang, Yan, Hao Wang, Ji Hu, and Hailan Hu. 2018. “Lateral Habenula in the Pathophysiology of Depression.” Current Opinion in Neurobiology, Neurobiology of Disease, vol. 48 (February): 90–96. 10.1016/j.conb.2017.10.024.

Zahm, Daniel S., and David H. Root. 2017. “Review of the Cytology and Connections of the Lateral Habenula, an Avatar of Adaptive Behaving.” Pharmacology, Biochemistry, and Behavior 162 (November): 3–21. 10.1016/j.pbb.2017.06.004.

Zeng, Hongkui. 2022. “What Is a Cell Type and How to Define It?” Cell 185 (15): 2739–55. 10.1016/j.cell.2022.06.031.

Zhang, Limei, Vito S. Hernández, Jerome D. Swinny, et al. 2018. “A GABAergic Cell Type in the Lateral Habenula Links Hypothalamic Homeostatic and Midbrain Motivation Circuits with Sex Steroid Signaling.” Translational Psychiatry 8 (1): 50. 10.1038/s41398-018-0099-5.

Zhang, Yong, Tao Liu, Clifford A. Meyer, et al. 2008. “Model-Based Analysis of ChIP-Seq (MACS).” Genome Biology 9 (9): R137. 10.1186/gb-2008-9-9-r137.

Zhao, Edward, Matthew R. Stone, Xing Ren, et al. 2021. “Spatial Transcriptomics at Subspot Resolution with BayesSpace.” Nature Biotechnology 39 (11): 1375–84. 10.1038/s41587-021-00935-2.

Zheng, Zhi-Wei, Peng-Xiang Min, Yi-Ling Luo, et al. 2026. “Molecular Brake on Firing Pattern Transitions in MHbChAT Neurons to Mediate Nicotine-Withdrawal-Induced Anxiety.” Neuron 114 (13): 2441–2455.e7. 10.1016/j.neuron.2026.02.020.

Zhou, Hang, Rachel L. Kember, Joseph D. Deak, et al. 2023. “Multi-Ancestry Study of the Genetics of Problematic Alcohol Use in over 1 Million Individuals.” Nature Medicine 29 (12): 3184–92. 10.1038/s41591-023-02653-5.

Zhu, Yuanzhen, Guang Li, Peter W. H. Holland, Günter P. Wagner, and Sebastian M. Shimeld. 2026. “Towards a Rational Terminology for Cell Types.” No. 2026061059. Preprint, Preprints, June 12. 10.20944/preprints202606.1059.v1.

