## Supplementary Figures and Tables for "Multi-omic spatial mapping of the human habenula defines the molecular and anatomical organization of medial and lateral subregions"

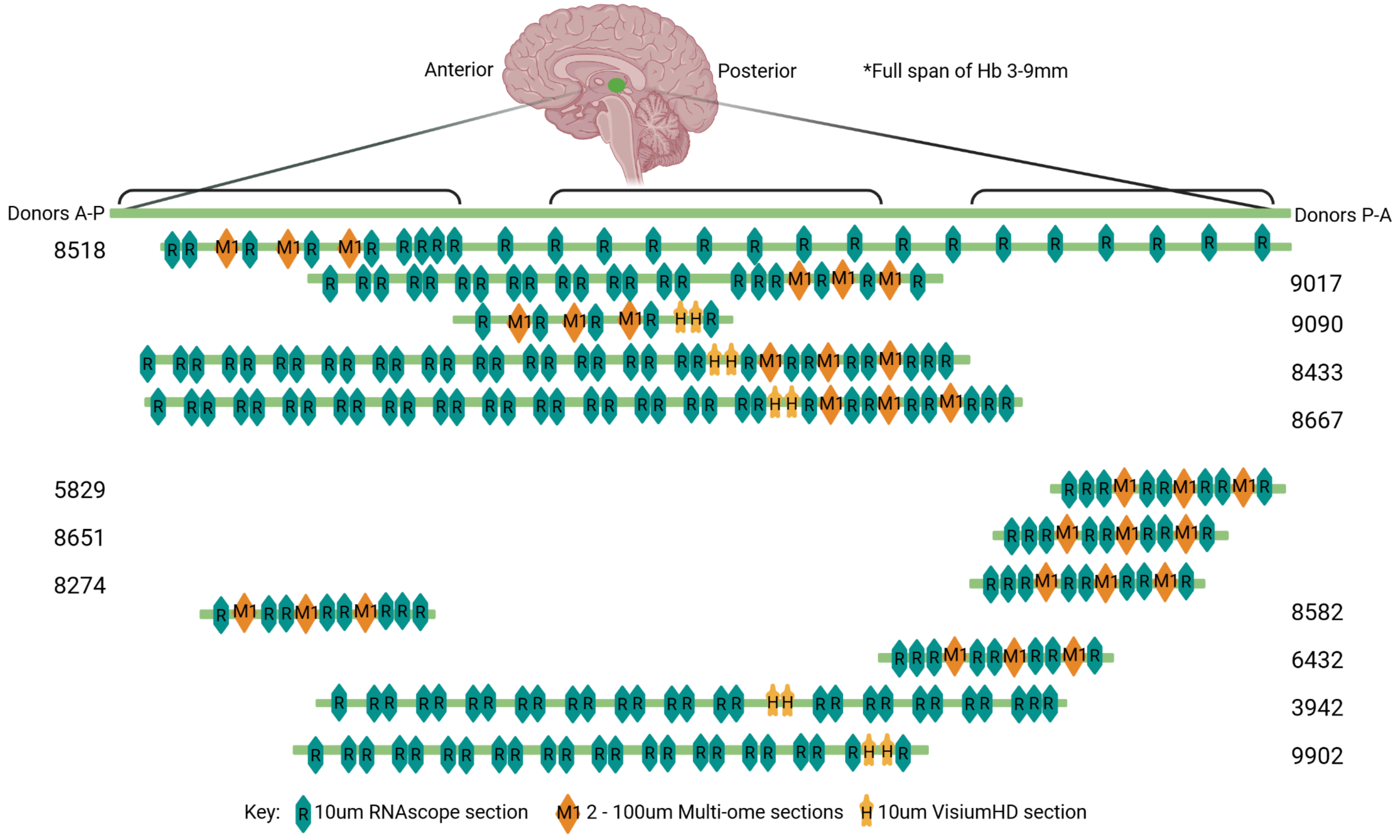


**Fig. S1: Approximate Hb tissue collection per donor.** Collection strategy was individualized per donor based on the anatomic verification RNAscope. The sagittal brain diagram shows the anterior (A) to posterior (P) direction from which each donor was cut. Along the sides are the brain numbers of each donor; the side on which the number is listed dictates the cutting face and directionality of cryosectioning for each Hb tissue block. An approximate start site at which tissue was collected for each assay is represented by each donors' green bar. In general, the following collection series was repeated until RNAscope was no longer positive for putative Hb markers: two 100um sections for snMultiome (orange diamond), two 10 um sections for RNAscope (blue hexagon). Depending on the results of the initial anatomic verification RNAscope, the series was adjusted, swapping out the two10um RNAscope cryosections for two 10um VisiumHD sections (yellow star). VisiumHD sections were targeted at a plane capturing both MHb and LHb at their largest volumes.

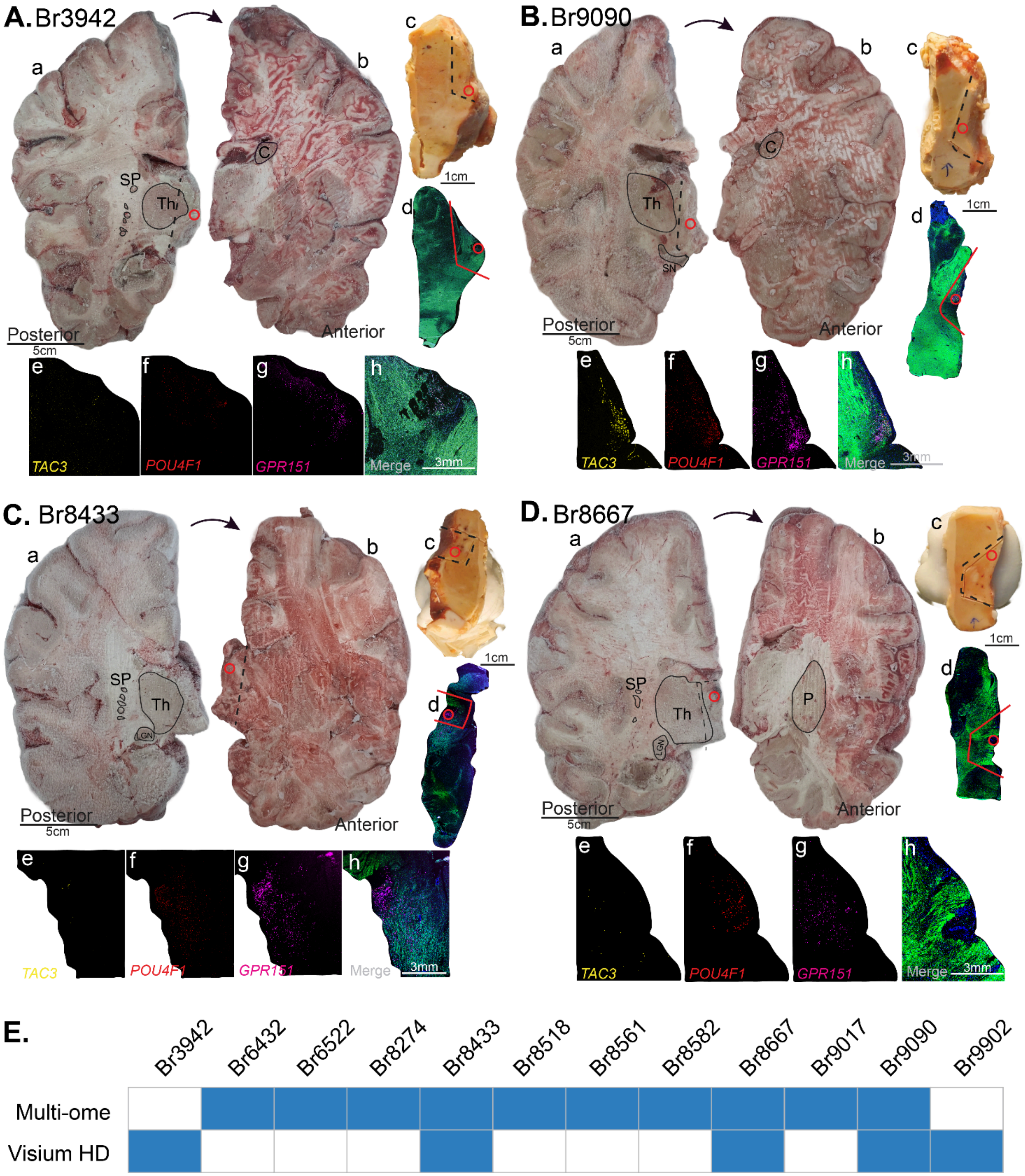

A. Br3942
a
b
c
d
SP
Th
C
1cm
Posterior
5cm
Anterior
e
f
g
h
TAC3
POU4F1
GPR151
Merge
3mm
B. Br9090
a
b
c
d
Th
C
SN
1cm
Posterior
5cm
Anterior
e
f
g
h
TAC3
POU4F1
GPR151
Merge
3mm
C. Br8433
a
b
c
d
SP
Th
LGN
1cm
Posterior
5cm
Anterior
e
f
g
h
TAC3
POU4F1
GPR151
Merge
3mm
D. Br8667
a
b
c
d
SP
Th
LGN
P
1cm
Posterior
5cm
Anterior
e
f
g
h
TAC3
POU4F1
GPR151
Merge
3mm
E.
Br3942
Br6432
Br6522
Br8274
Br8433
Br8518
Br8561
Br8582
Br8667
Br9017
Br9090
Br9902
Multi-ome
Visium HD

**Fig. S2: Anatomical localization of Hb within tissue slabs and summary of data collection per donor.** Anterior and posterior frozen brain slab faces for four donors used in this study are shown with the surrounding landmark structures demarcated. Dissected tissue blocks along with RNAscope for *TAC3*, *POU4F1*, *GPR151* from the entire section and higher magnification insets of positive Hb signals. (**A.a.**) Donor 3942, on the posterior face of the slab, the segmented putamen “SP” and thalamus “Th” were identified. (**A.b.**) On the anterior face, the caudate “C” was observed. The dashed line shows where the block in (**A.c.**), was dissected and the red circle demarcates the Hb. (**A.c.**) Dissected tissue block with dashed line showing the scoring marks. Scored tissue was used in the specified assay(s) and the red circle demarcates the Hb. (**A.d.**) RNAscope image of the entire tissue section when the Hb was first identified. Score marks and Hb identification are labeled in red. (**A.e-h.**) High magnification single channel fluorescent RNAscope images for MHb and LHb marker genes. (**B**, **C**, **D**) Similar to A with variations specific to each donor. For instance, in (B), Substantia Nigra “SN” was identified and in (C) and (D) lateral geniculate nucleus “LGN” was observed. (**E**) Lists each donor included in this study and the assays collected (marked by a blue rectangle). Abbreviations: Caudate (C), Lateral geniculate nucleus (LGN), Putamen (P), Segmented Putamen (SP), Substantia Nigra (SN), Thalamus (Th).

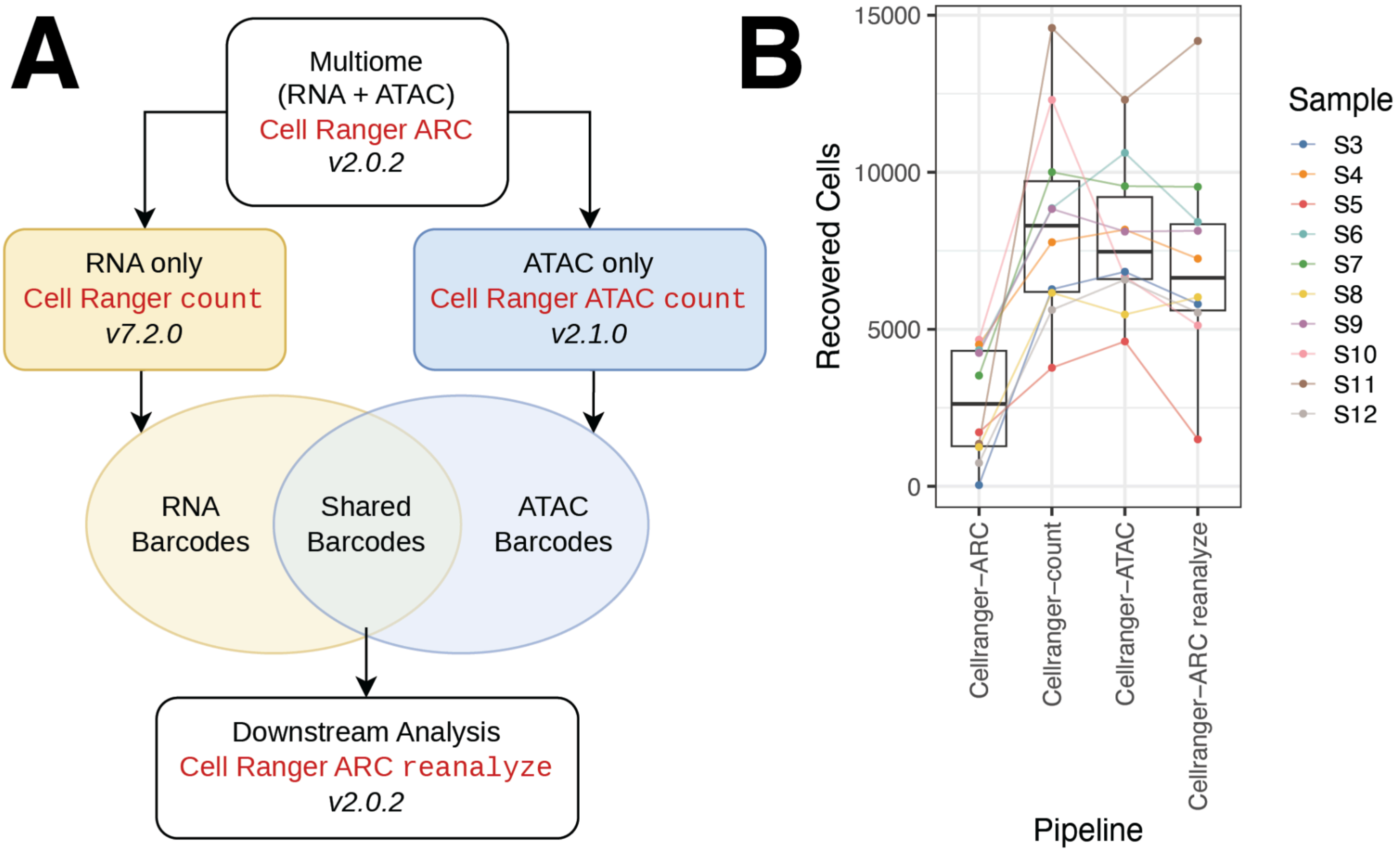


**Fig. S3: Custom *Cell Ranger ARC* pipeline for processing the snMultiome raw data.** (**A**) FASTQs were initially processed with *Cell Ranger ARC*, but joint cell-calling failed owing to assay-specific QC restrictions. To recover valid cells, RNA and ATAC libraries were reprocessed independently using *Cell Ranger* `count` and *Cell Ranger ATAC*, generating standalone sets of high-confidence barcodes (**Methods 4.4**). Because RNA and ATAC libraries use distinct barcode sequences, standalone barcodes passing their respective QC steps were translated to their corresponding multiome barcodes by line-number matching. The intersection of translated RNA and ATAC barcodes yielded the final curated set of shared barcodes. This shared RNA and ATAC valid barcodes set was provided to *Cell Ranger ARC* `reanalyze`, which performed downstream dimensionality reduction, clustering, and feature-gene linkage analysis. (**B**) Boxplots of the number of cells passing QC by each *Cell Ranger* pipeline.

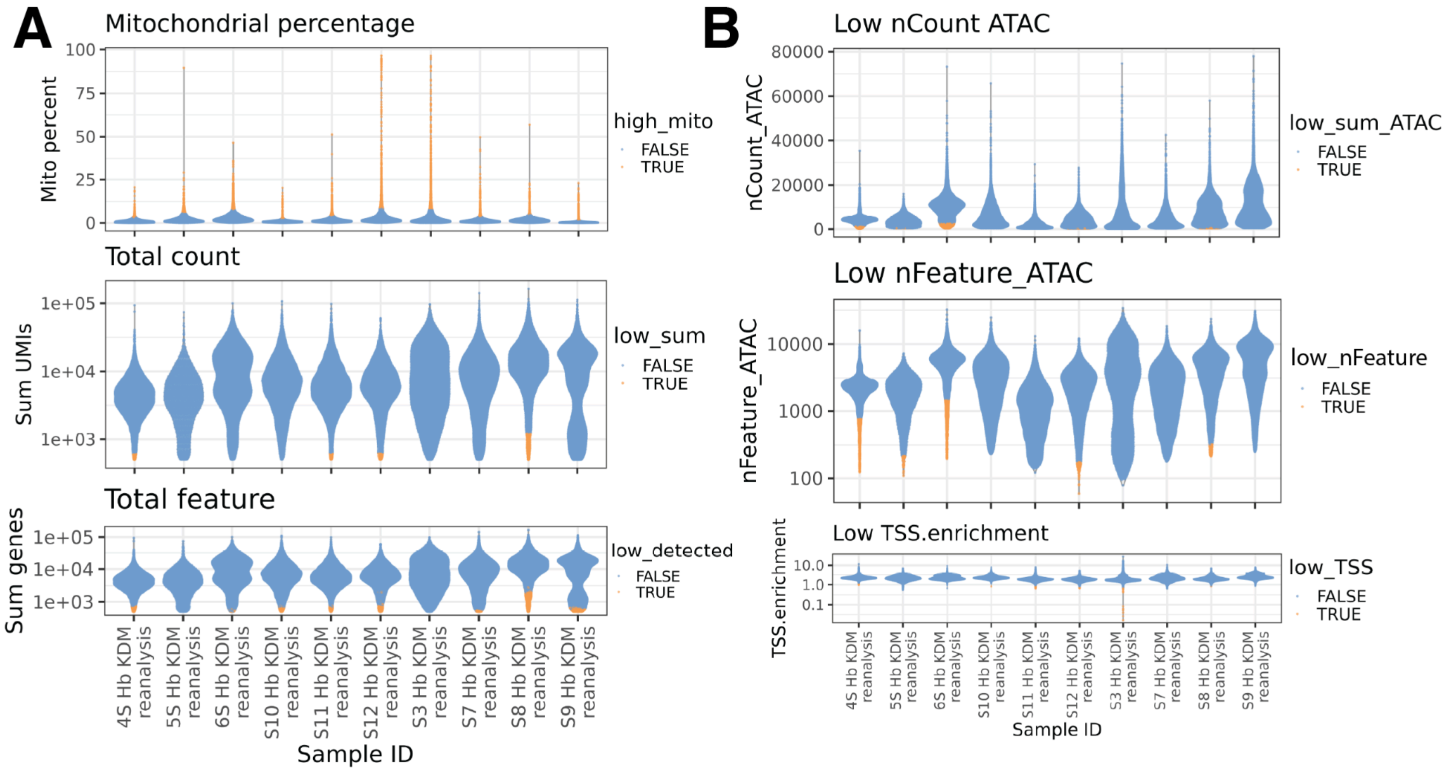


**Fig. S4: RNA and ATAC multiome data quality control metrics.** (**A**) Violin plots for the RNA quality metrics: mitochondrial gene expression percent, total number of RNA unique molecular identifiers (UMIs) counts (total count), and total number of detected genes (total feature). (**B**) Violin plots for the ATAC quality metrics: total number of ATAC UMI counts (nCount ATAC), number of detected fragments (nFeature ATAC), and Transcription Start Site (TSS) enrichment scores (TSS enrichment).

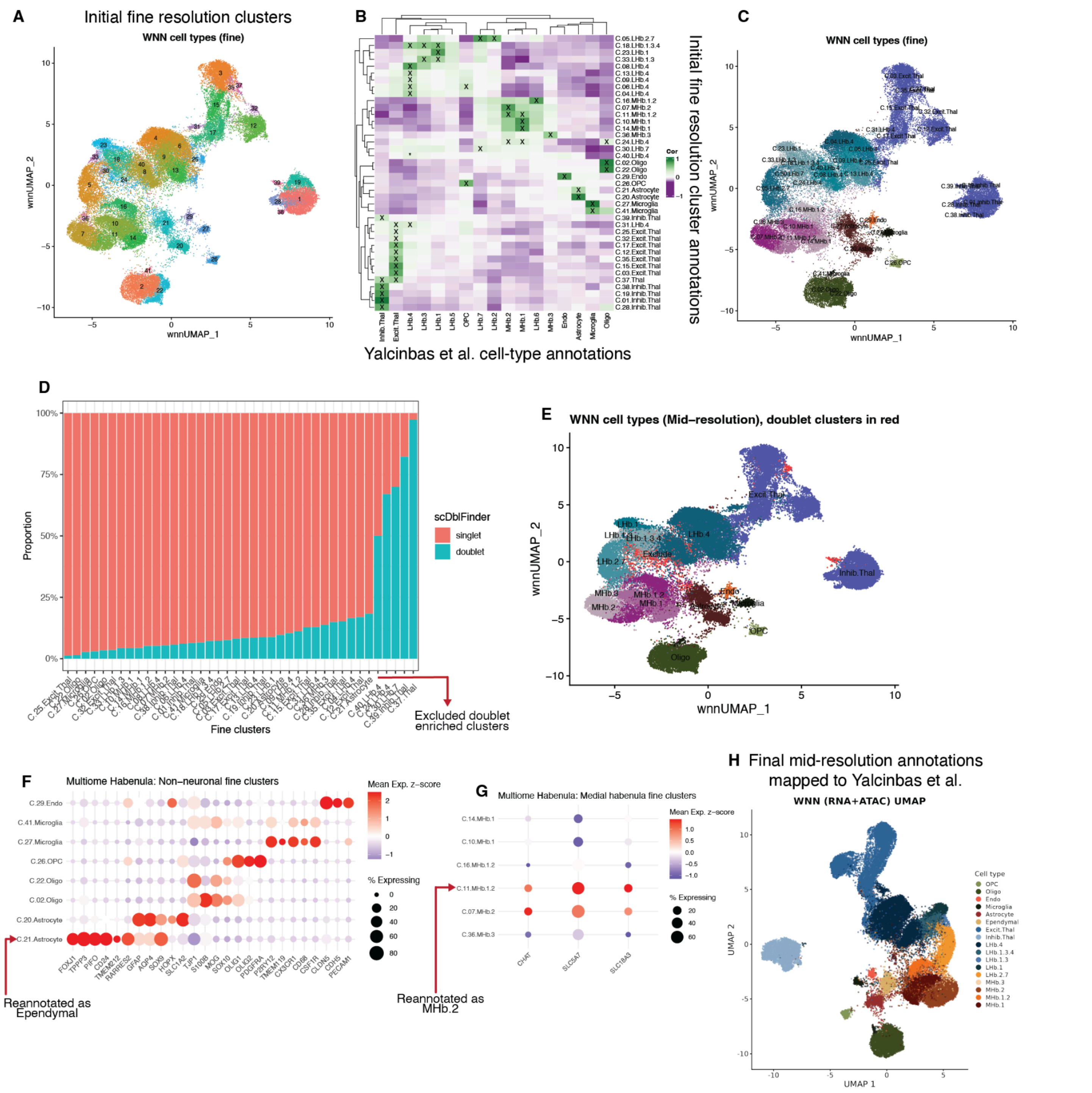


**Fig. S5: Annotation of fine and mid-resolution snMultiome clusters.** (**A**) WNN UMAP of 40 fine resolution clusters. (**B**) Transcriptional cluster registration between the Hb snMultiome fine resolution clusters (rows) and Yalcinbas et al. annotated clusters (columns). Correlations are computed across transcriptome-wide 1-vs-All DE t-statistics to identify transcriptionally similar clusters, with the 'x' representing high confidence matches (**Methods 4.20**). (**C**) WNN UMAP of the initial cluster annotations derived from panel B. (**D**) Stacked barplots depicting the proportion of cells annotated as doublets per fine resolution cluster. We excluded clusters with > 50% called doublets. (**E**) WNN UMAP with the doublet excluded clusters in red. (**F**) Bubble plot of non-neuronal marker genes, highlighting a fine resolution astrocyte cluster as an ependymal population. (**G**) Bubble plot of cholinergic marker genes, highlighting a fine resolution MHb.1.2 cluster as a cholinergic cluster. (**H**) WNN UMAP with annotated clusters at the mid-resolution used as the starting point for cross-species comparisons.

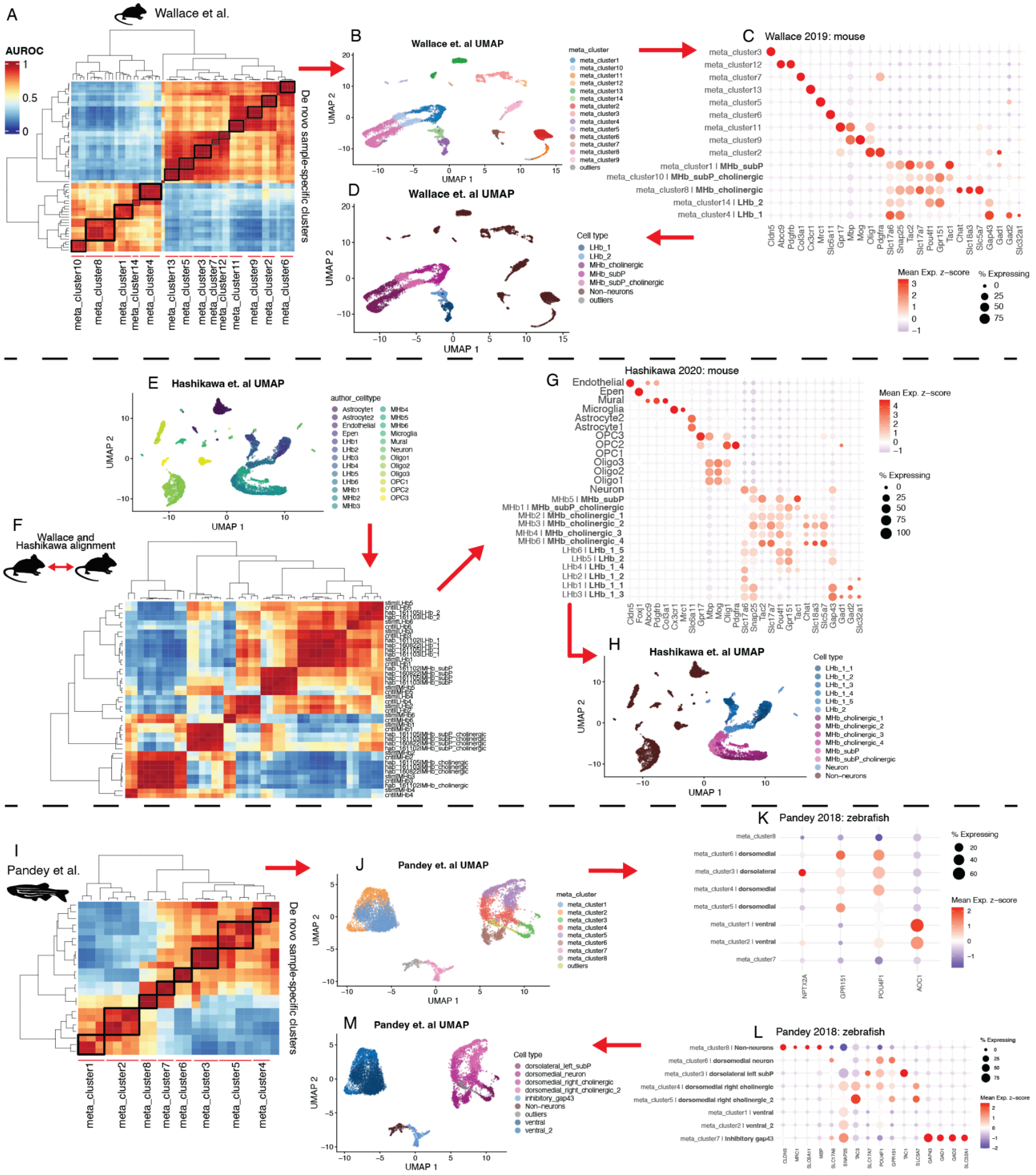

A
Wallace et al.
AUROC
De novo sample-specific clusters
B
Wallace et. al UMAP
C
Wallace 2019: mouse
D
Wallace et. al UMAP
Cell type
LHb_1
LHb_2
MHb_cholinergic
MHb_subP
MHb_subP_cholinergic
Non-neurons
outliers
E
Hashikawa et. al UMAP
F
Wallace and Hashikawa alignment
G
Hashikawa 2020: mouse
H
Hashikawa et. al UMAP
I
Pandey et al.
J
Pandey et. al UMAP
K
Pandey 2018: zebrafish
L
Pandey 2018: zebrafish
M
Pandey et. al UMAP

**Fig. S6: Mouse and Zebrafish reannotations.** Hierarchically clustered heatmaps of pairwise *MetaNeighbor* Area Under the Receiver Operating Characteristic curve (AUROC) scores, quantifying cross-cluster transcriptomic replicability for three different datasets: (**A**) mouse (Wallace et al. 2020), (**F**) across two mouse datasets (Wallace et al. 2020; Hashikawa et al. 2020), or (**I**) zebrafish (Pandey et al. 2018). In (A) and (I), metaclusters, replicable groups of de novo clusters, are in bold. (**B**) UMAP of the Wallace et al. dataset (A) annotated by replicable *MetaNeighbor*-identified clusters. (**C**) Marker genes for the clusters in (B), where the intensity of the color and size of the spot denote the expression of the respective gene. Cluster names in the rows show in bold the initial Hb cluster mouse annotations (**Fig. 2**). (**D**) Similar to (B), but colored by initial Hb cluster mouse annotations (**Fig. 2**). (**E**) UMAP of the Hashikawa et al. dataset with the original cluster labels. (**G**) Similar to (C), but for the Hashikawa et al. mouse dataset. (**H**) Similar to (D), but for the Hashikawa et al. dataset. (**I**) Similar to (A), but for the Pandey et al. zebrafish dataset . (**J**) Similar to (B), but for the Pandey et al. dataset. (**K**) Similar to (C), but for zebrafish regional Hb markers; *NPTX2A* (dorsolateral), *GPR151* and *POU4F1* (dorsomedial), and *AOC1* (ventral). (**L**) Similar to (C), but prioritizing neuronal gene markers. Cell-type annotations for the *MetaNeighbor* metaclusters were further refined in (L). (**M**) Similar to (J), but with the refined cluster names from (L).

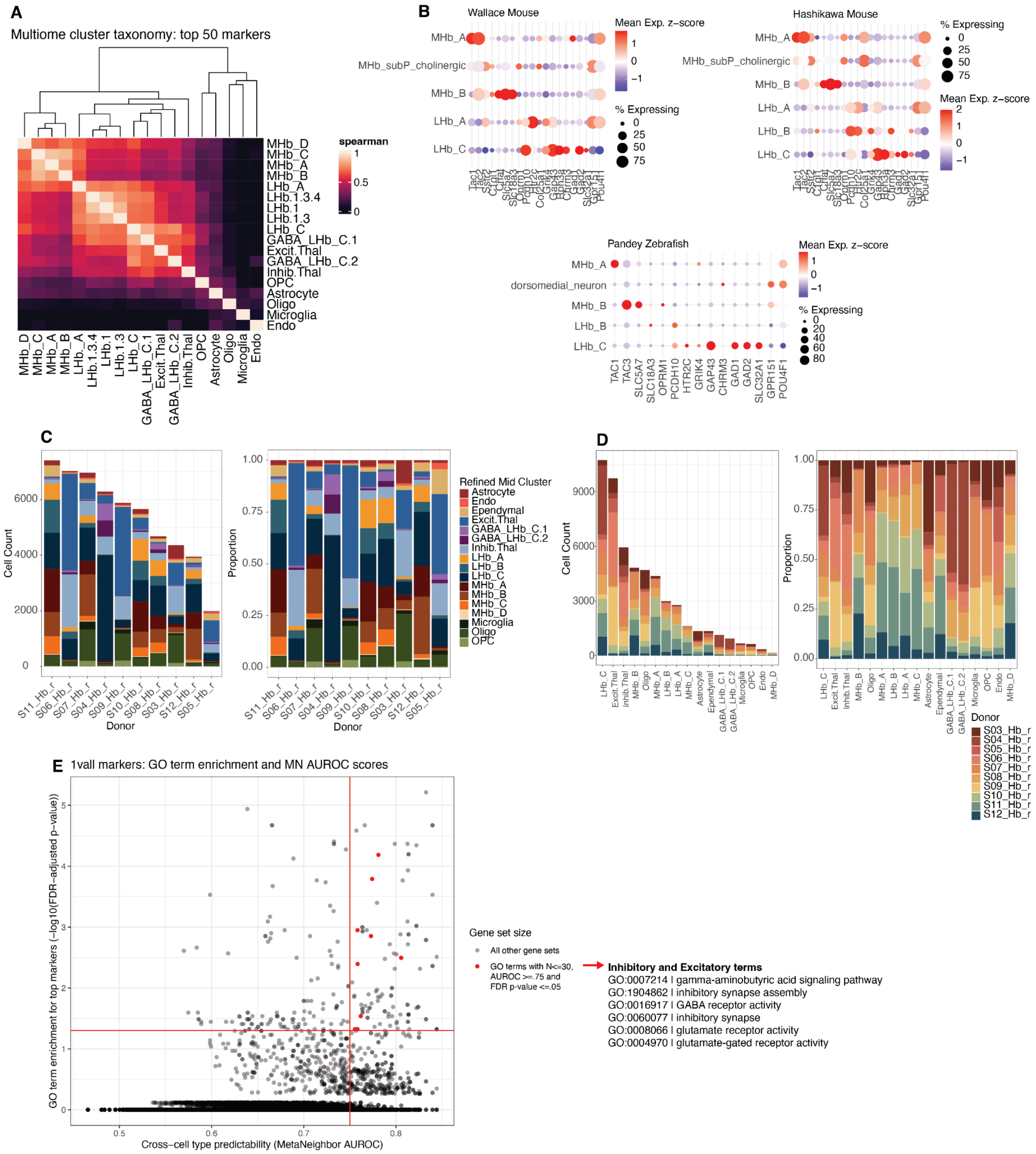
A
Multiome cluster taxonomy: top 50 markers
spearman
1
0.5
0
MHb_D
MHb_C
MHb_A
MHb_B
LHb_A
LHb.1.3.4
LHb.1
LHb.1.3
LHb_C
GABA_LHb_C.1
Excit.Thal
GABA_LHb_C.2
Inhib.Thal
OPC
Astrocyte
Oligo
Microglia
Endo
B
Wallace Mouse
Hashikawa Mouse
Pandey Zebrafish
Mean Exp. z−score
% Expressing
MHb_subP_cholinergic
dorsomedial_neuron
LHb_B
TAC1
TAC3
SLC5A7
SLC18A3
OPRM1
PCDH10
HTR2C
GRIK4
GAP43
CHRM3
GAD1
GAD2
SLC32A1
GPR151
POU4F1
C
Cell Count
Proportion
Donor
Refined Mid Cluster
Astrocyte
Endo
Ependymal
Excit.Thal
GABA_LHb_C.1
GABA_LHb_C.2
Inhib.Thal
LHb_A
LHb_B
LHb_C
MHb_A
MHb_B
MHb_C
MHb_D
Microglia
Oligo
OPC
S11_Hb_r
S06_Hb_r
S07_Hb_r
S04_Hb_r
S09_Hb_r
S10_Hb_r
S08_Hb_r
S03_Hb_r
S12_Hb_r
S05_Hb_r
D
Donor
S03_Hb_r
S04_Hb_r
S05_Hb_r
S06_Hb_r
S07_Hb_r
S08_Hb_r
S09_Hb_r
S10_Hb_r
S11_Hb_r
S12_Hb_r
E
1vall markers: GO term enrichment and MN AUROC scores
GO term enrichment for top markers (−log10(FDR−adjusted p−value))
Cross−cell type predictability (MetaNeighbor AUROC)
Gene set size
All other gene sets
GO terms with N<=30, AUROC >=.75 and FDR p-value <=.05
Inhibitory and Excitatory terms
GO:0007214 | gamma-aminobutyric acid signaling pathway
GO:1904862 | inhibitory synapse assembly
GO:0016917 | GABA receptor activity
GO:0060077 | inhibitory synapse
GO:0008066 | glutamate receptor activity
GO:0004970 | glutamate-gated receptor activity

**Fig. S7: Final snMultiome cross-species consensus cell-type annotations** (**A**) Hierarchical clustering of transcriptional similarity between mid-resolution snMultiome clusters. (**B**) Bubble plots of top human snMultiome markers per matched consensus Hb cell type in the Wallace, Hashikawa, and Pandey et al. non-human datasets (left to right). (**C**) Stacked barplots of cell counts (left) and cell proportions (right) of each final snMultiome cross-species consensus cluster per snMultiome human donor (columns). (**D**) Stacked barplots of cell counts (left) and cell proportions (right) of each human donor per final snMultiome cross-species consensus cluster (columns). (**E**) Scatter plot identifying GO terms that are enriched in cell-type marker gene sets (*Y*-axis) and contribute to cell-type replicability across donors (*X*-axis, **Methods 4.12**) for the consensus snMultiome Hb cell types. GO terms are filtered for gene set size, as well as thresholded on the two statistics (highlighted in red), with the 6 GO terms used to identify the genes used in **Fig. 3A** and **D** in text.

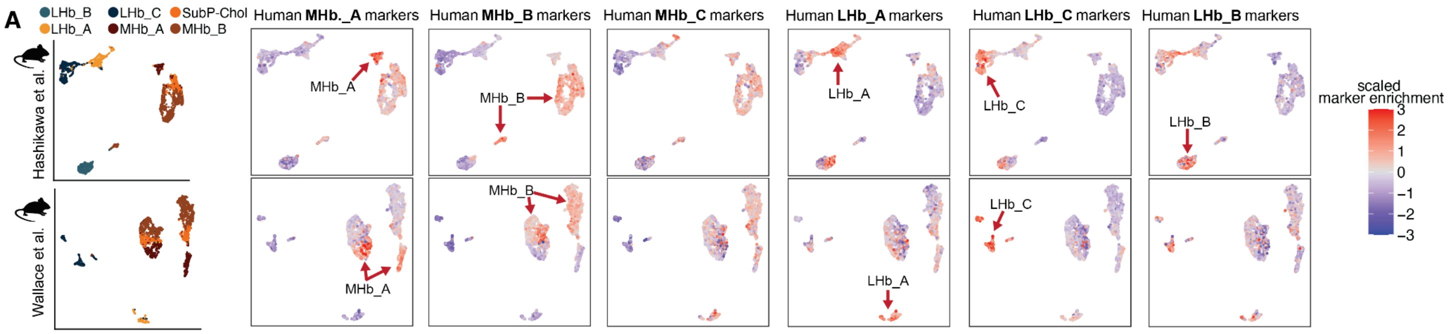


**Fig. S8: Projection of top human snMultiome marker genes to mouse datasets.** (**A**) UMAPs of the Hb neuron clusters in the Hashikawa et al. (top row) and Wallace et al. (bottom row) mouse datasets. The first two UMAPs are colored by the cross-species consensus cell type annotations and in the following UMAPS, cells are colored by *Z*-scored marker enrichment of aggregate human snMultiome marker sets (**Methods 4.8**).

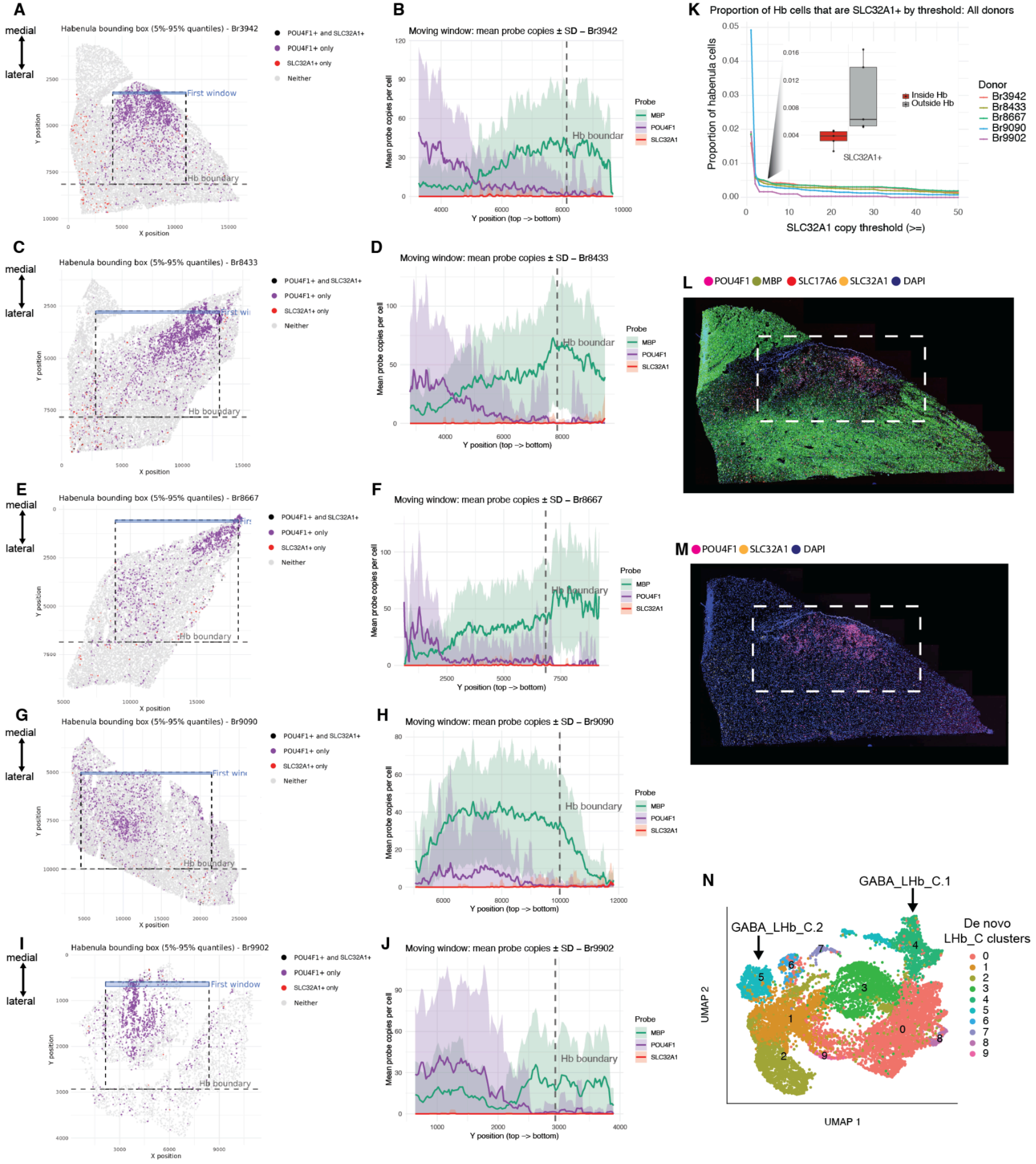
A
medial
lateral
Habenula bounding box (5%-95% quantiles) - Br3942
POU4F1+ and SLC32A1+
POU4F1+ only
SLC32A1+ only
Neither
First window
Hb boundary
Y position
X position
B
Moving window: mean probe copies ± SD – Br3942
Mean probe copies per cell
Y position (top –> bottom)
Probe
MBP
POU4F1
SLC32A1
Hb boundar
C
Habenula bounding box (5%-95% quantiles) - Br8433
D
Moving window: mean probe copies ± SD – Br8433
E
Habenula bounding box (5%-95% quantiles) - Br8667
F
Moving window: mean probe copies ± SD – Br8667
G
Habenula bounding box (5%-95% quantiles) - Br9090
H
Moving window: mean probe copies ± SD – Br9090
Hb boundary
I
Habenula bounding box (5%-95% quantiles) - Br9902
J
Moving window: mean probe copies ± SD – Br9902
K
Proportion of Hb cells that are SLC32A1+ by threshold: All donors
Proportion of habenula cells
SLC32A1 copy threshold (>=)
Inside Hb
Outside Hb
SLC32A1+
Donor
Br3942
Br8433
Br8667
Br9090
Br9902
L
POU4F1
MBP
SLC17A6
SLC32A1
DAPI
M
POU4F1
SLC32A1
DAPI
N
GABA_LHb_C.1
GABA_LHb_C.2
De novo
LHb_C clusters
0
1
2
3
4
5
6
7
8
9
UMAP 2
UMAP 1

**Fig. S9: RNAscope validation of rare GABAergic neurons in human Hb.** (**A**) Spatial plot of segmented nuclei for Br3942, colored by detected transcripts for *POU4F1*, *SLC32A1* (VGAT) or *POU4F1/SLC32A1*. Cells are colored if detected probes are >= 5. The bounding box defines the approximate Hb area, derived using the 5th and 95th percentiles of X and Y spatial coordinates for the *POU4F1*+ nuclei. The blue box denotes the starting window for computing the sliding average scores used in (B). (**B**) Line plot depicting the sliding average of probe counts for *MBP*, *POU4F1*, and *SLC32A1* (VGAT), with the Y-position of the sliding window from (A) on the *X*-axis and the mean transcript count ± SD (shaded area) on the *Y*-axis. The dotted line depicts the lateral boundary of the Hb area annotated in (A). (**C**) Same as (A), but for Br8433. (**D**) Same as (B), but for Br8433. (**E**) Same as (A), but for Br8667. (**F**) Same as (B), but for Br8667. (**G**) Same as (A), but for Br9090. (**H**) Same as (B), but for Br9090. (**I**) Same as (A), but for Br9902. (**J**) Same as (B), but for Br9902 (**K**) Line plot depicting the proportion of cells (*Y*-axis) within the Hb boundary boxes from (A), (C), (E), (G), (I) that are *SLC32A1*+ (VGAT+) dependent on an increasing threshold of *SLC32A1* (VGAT) transcript counts (*X*-axis). The inset boxplots show the proportion of *SLC32A1*+ nuclei within and outside the Hb boundary boxes for all donors at a transcript threshold of >= 5. (**L**) Representative RNAscope fluorescent image for *POU4F1*, *MBP*, *SLC17A6*, *SLC32A1* and DAPI, highlighting *SLC32A1* (yellow) transcripts detected in the approximate Hb *POU4F1*+ (pink) region. (**M**) Same as (L), but only with *POU4F1* and *SLC32A1* and DAPI channels. (**N**) UMAP (same as **Fig. 2C-D**) of de novo LHb_C clusters with the GABA_LHb_C.1 and GABA_LHb_C.2 clusters highlighted.

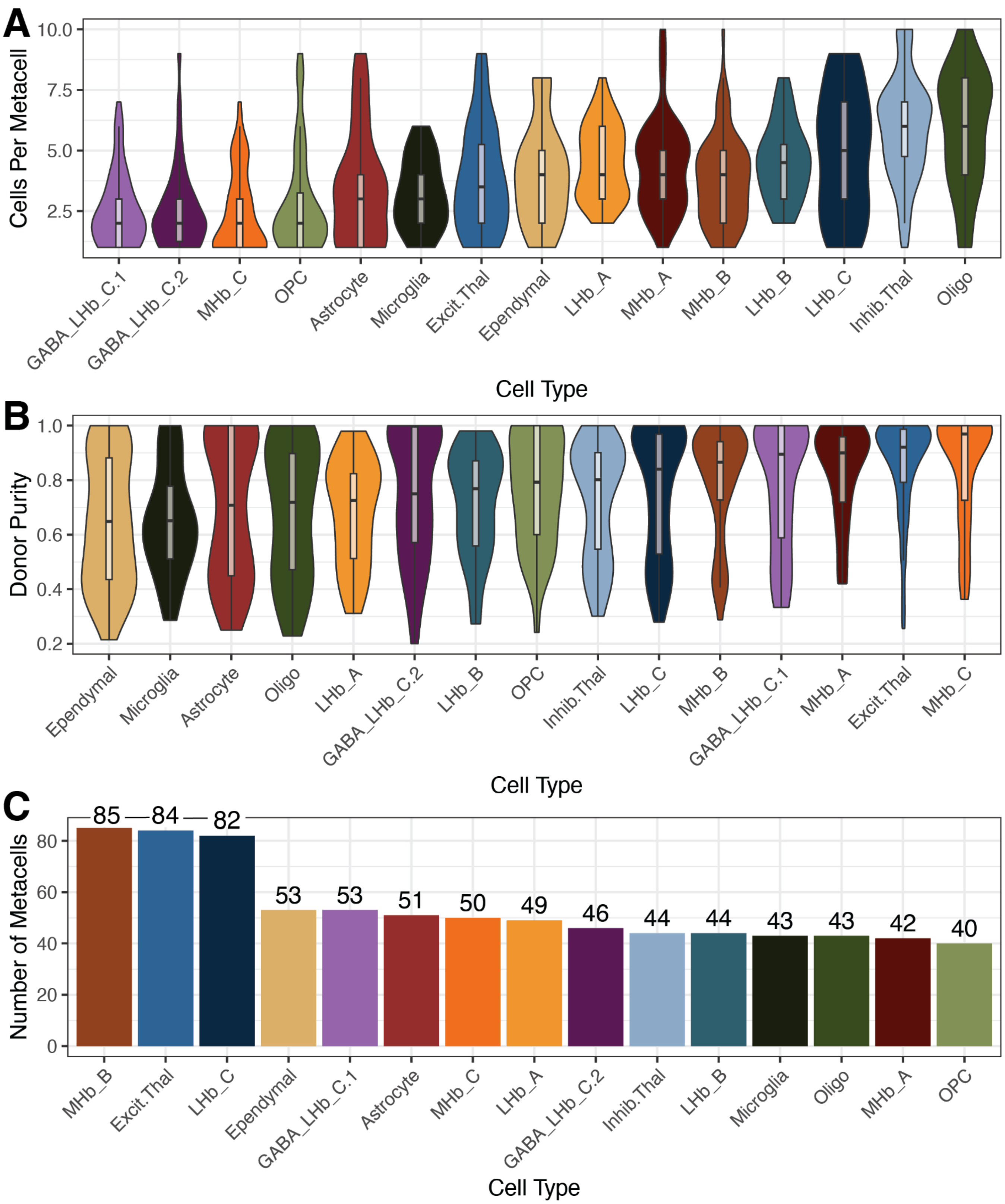


**Fig. S10: snMultiome metacell quality statistics.** (**A**) Each metacell is composed of a variable number of cells from the same cell type. (**B**) Metacells can be composed of cells from multiple donors. Donor purity (*Y*-axis) is defined as the ratio of the number of cells for the donor with the largest number against the number of cells in any other donor. (**C**) After aggregating cells into metacells, each cell type has a variable number of total metacells, between 40 and 85.

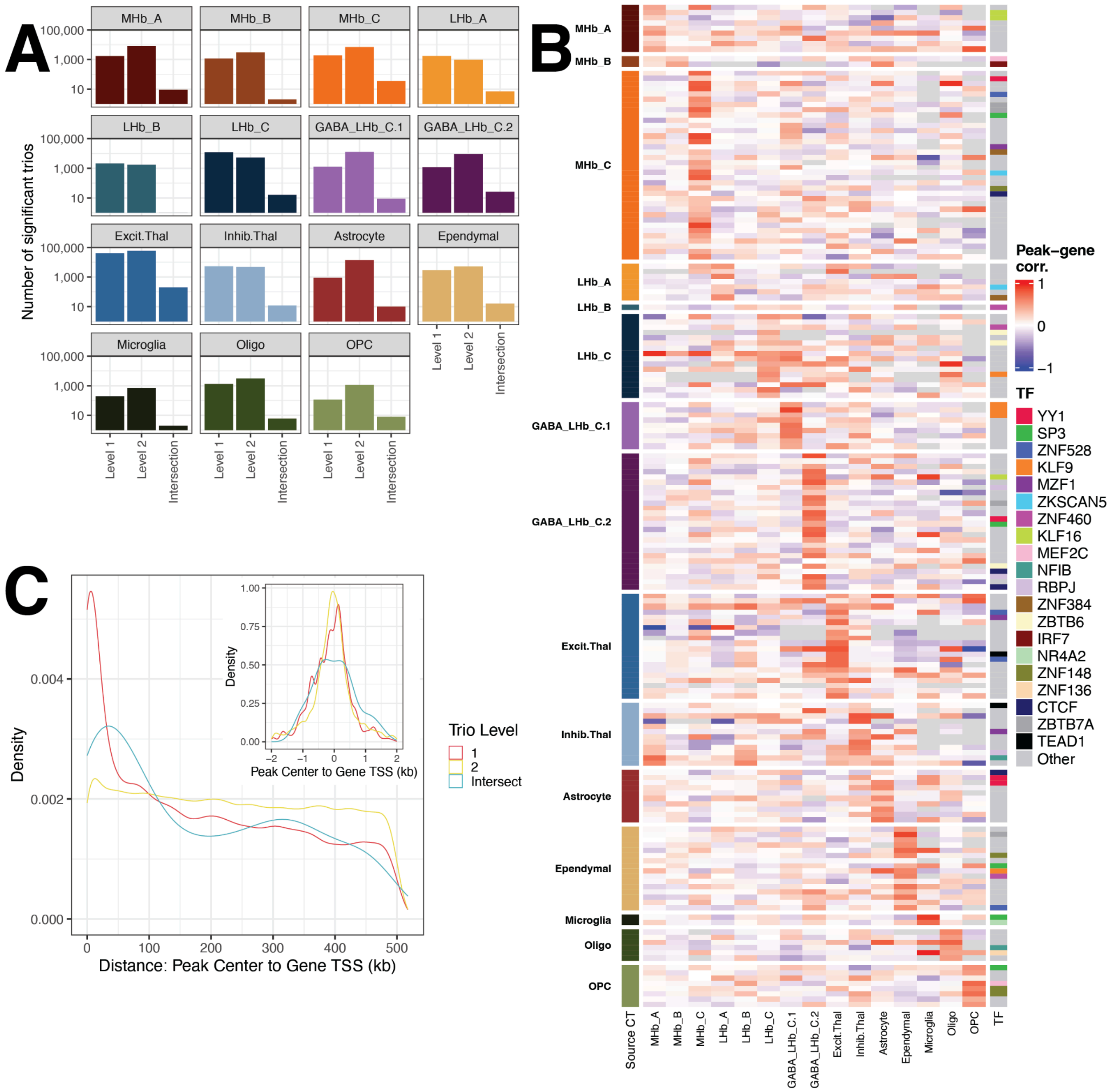


**Fig. S11: Peak-gene-transcription factor (TF) trios found for all snMultiome cell types.** (**A**) Distribution of the number of *TRIPOD* Level 1, Level 2, and intersecting Level 1 and 2 trios per cell type (**Methods 4.14**). *Y*-axis is shown in a $\log_{10}$ scale. (**B**) Correlation across each cell type's metacells between gene expression and peak accessibility is plotted for all intersection trios per cell type for all cell types (**Fig. 4D** only showed the neuronal cell types). Only the top-20 Excit.Thal trios by significance are shown; there are 198 Excit.Thal trios total. Each row is a trio, and 20 of the most common TFs color these rows. Row groups indicate the cell type where the trio was found, while columns indicate the cell type in which expression and accessibility are measured. (**C**) Frequency density of the number of *TRIPOD* trios identified relative to the transcription start site (TSS) in kb to the closest gene for each ATAC peak. The inset shows the -2 to +2 kb window around the TSS.

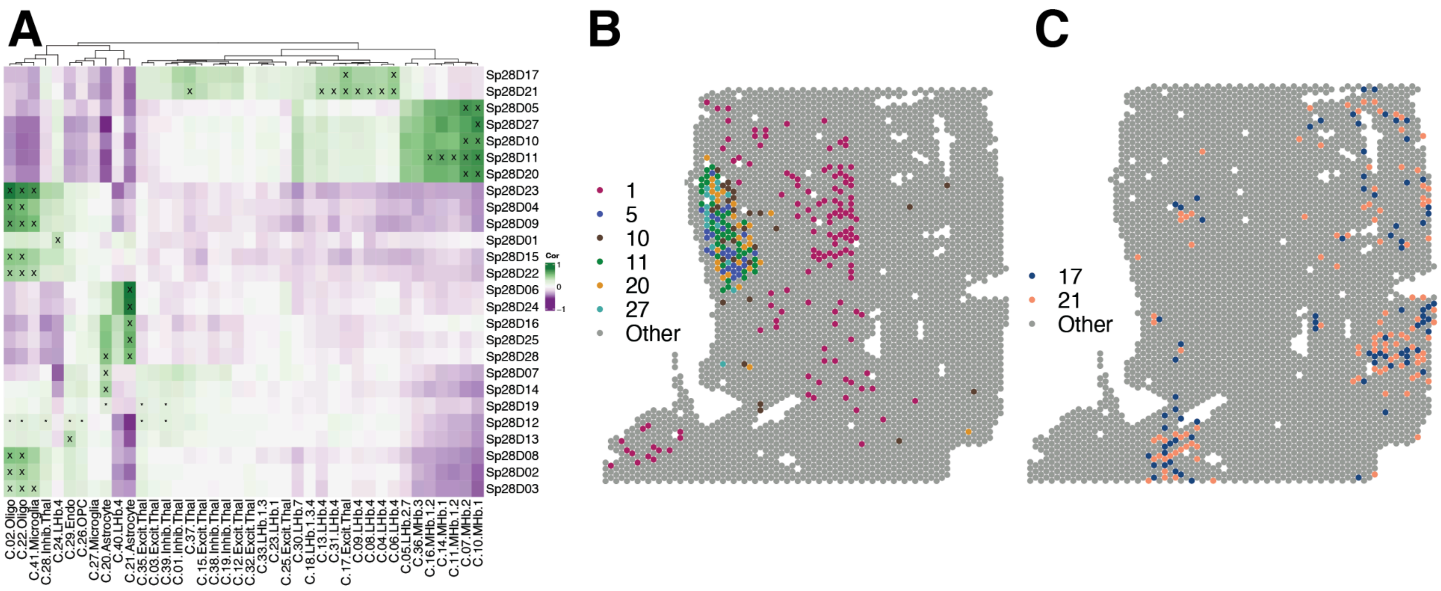


**Fig. S12: Visium was unable to separate LHb and MHb spatially-aware clusters.** (**A**) Spatial registration plot of the Visium *BayesSpace* *k* = 28 clusters (Y-axis) against the snMultiome fine resolution data (*X*-axis). A total of 6 Visium H&E *BayesSpace* spatially-aware clusters registered to LHb and MHb, with none of them separating the two from each other. (**B**) Spotplots showing the location of the 6 *BayesSpace* spatially-aware clusters registered to LHb and MHb. (**C**) Spotplots showing the location of the 2 *BayesSpace* spatially-aware clusters registered to excitatory and inhibitory thalamus.

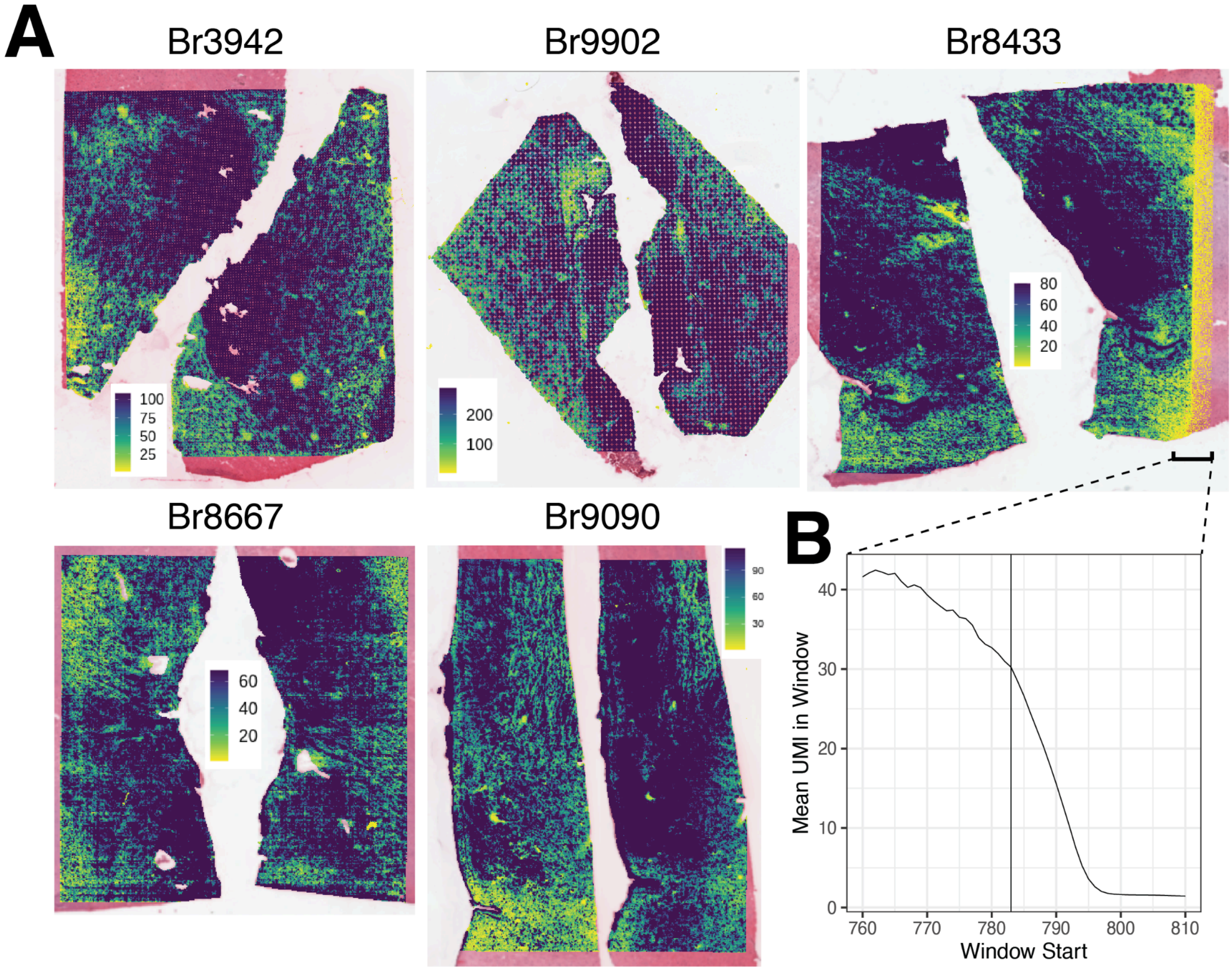


**Fig. S13: VisiumHD quality control and removal of technical artifacts.** (**A**) Total unique molecular identifier counts (UMI) plotted spatially across the 8μm-bin-level data by donor, with values limited to the donor-specific median or less to highlight areas of unusually low UMI. (**B**) For Br8433, average UMI (*Y*-axis) within a sliding 10-unit window in `array_col` is plotted against the window start coordinate (*X*-axis) for windows close to the visible boundary of the technical artifact. At `array_col = 783`, a smooth, precipitous decline begins, marking the boundary of the artifact.

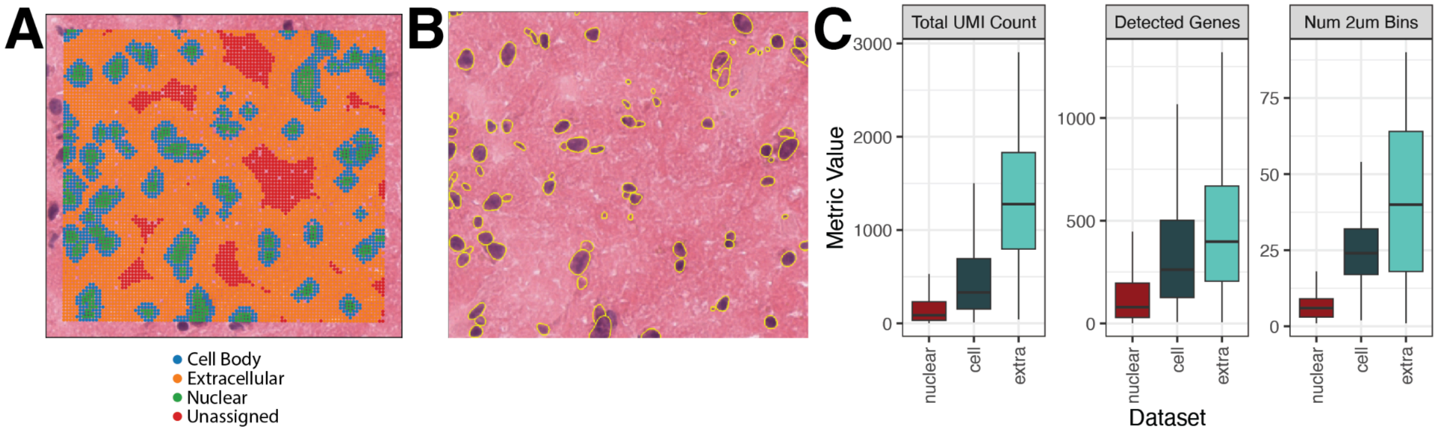


**Fig. S14: Categorization of nuclear, cellular, and extracellular bins**. (**A**) In a small subregion of Br8667, 2 µm bins are labeled. Nuclear bins are determined by segmentation of the H&E image, and cell bodies are estimated by slightly expanding around nuclei (**Methods 4.23**). A larger expansion from each cell body is used to encapsulate bins that are deemed "extracellular" (**Methods 4.22**). Red bins do not belong to any of these categories and were only used for the "all-bin" *FICTURE* analyses (**Fig. S19A**). (**B**) Same subregion as in (A), showing the cell segmentation boundaries on the H&E image. Boxplots showing typical quality control metrics across 2 µm bins in the full dataset among bin categories in (A) for the: (**C**) total UMI, (**D**) total detected genes, and (**E**) total number of 2 µm bins.

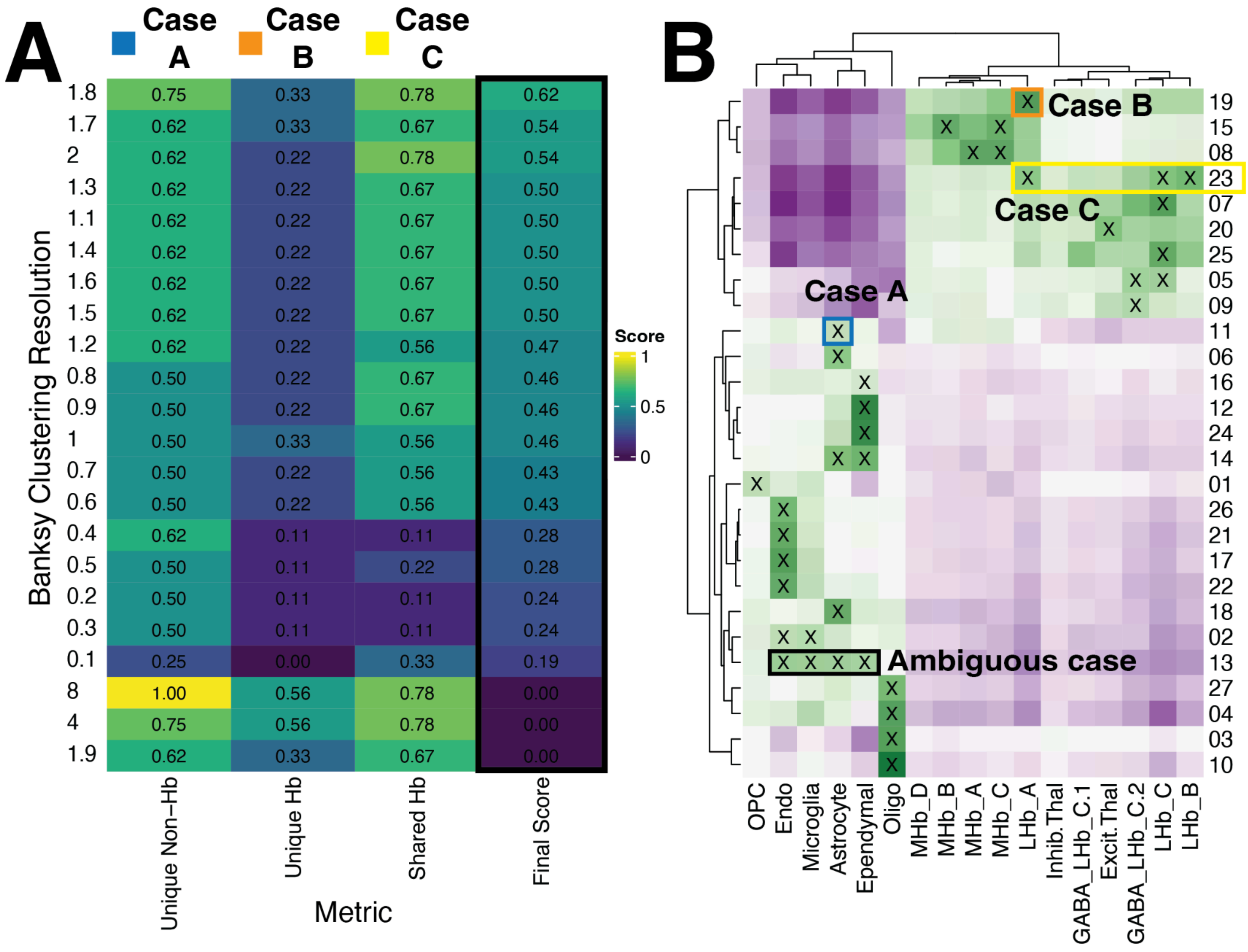


**Fig. S15: Spatial registration-based metrics to rank VisiumHD clustering resolutions**. (**A**) Heatmap showing the 3 scores and their average for the proportion-based metrics measured for candidate *BANKSY* clustering resolutions. The 3 metrics have corresponding cases A, B, and C, respectively, showing examples of clusters positively contributing to each metric. The *unique non-Hb*, *unique Hb*, *shared Hb*, and *final score* metrics are described in detail in **Methods 4.23**. For each metric, higher values were desired with some additional filters applied to remove ambiguous clusters (**Methods 4.23**), resulting in resolution 1.8 as the set of clusters achieving the best balance across all metrics. (**B**) Spatial registration results for *BANKSY* resolution 1.8, using the snMultiome cell types as a reference. In case A, cluster 11 contributes to the *unique non-hb* metric for resolution 1.8 by uniquely mapping to astrocytes. Similarly, cluster 19 maps uniquely to the Hb cell type LHb_A, boosting the *unique hb* metric in case B. In case C, cluster 23 contributes to the *shared hb* metric by mapping to several LHb subtypes. Finally, in an ambiguous case, cluster 13 maps to several distinct non-Hb cell types; clustering resolutions with more than 5 such clusters receive a score of 0 in (A).

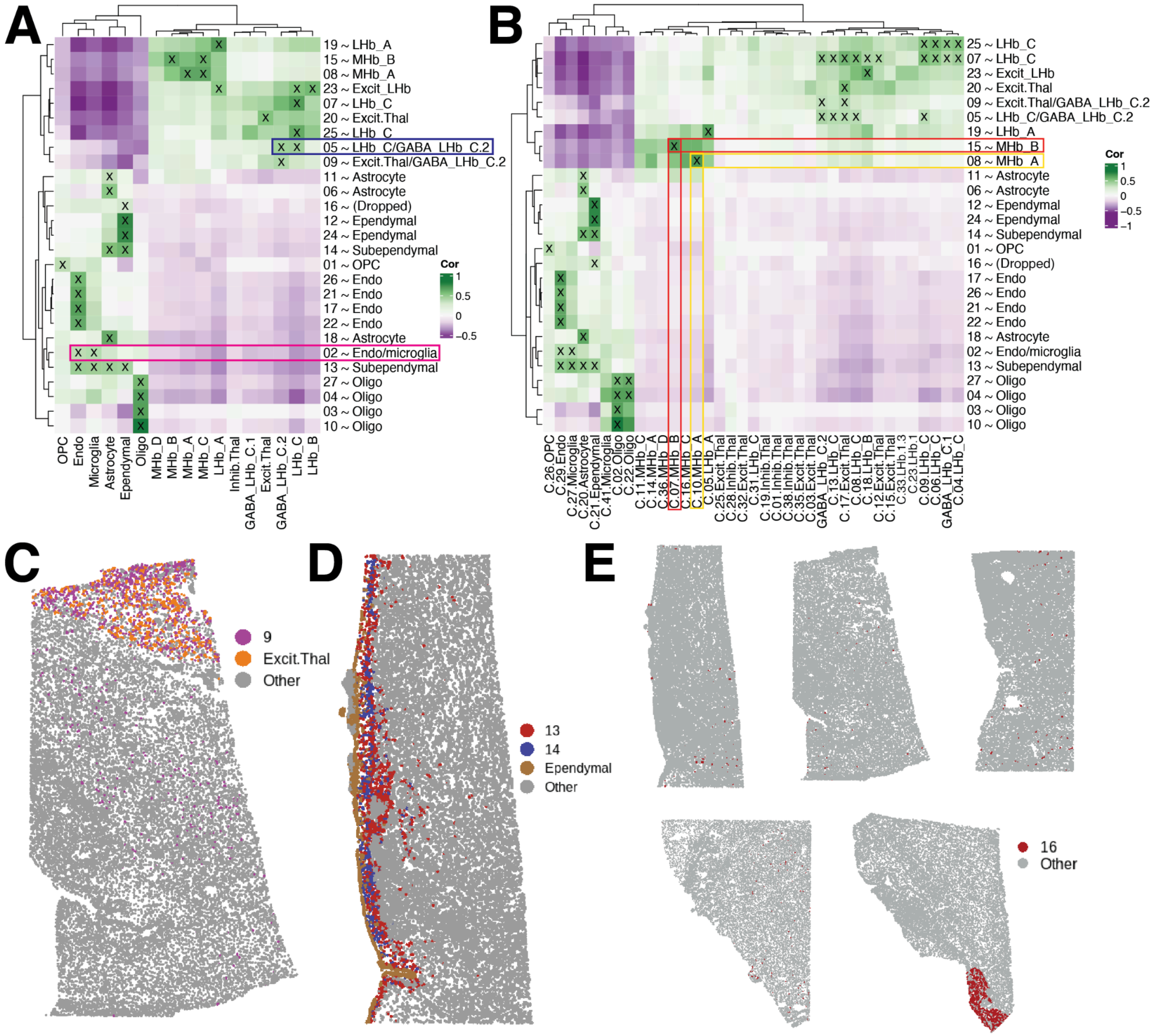


**Fig. S16: Annotation of VisiumHD *BANKSY* clusters**. (**A**) Spatial registration of *BANKSY* clusters to mid-resolution snMultiome clusters provides the main reference for annotating the VisiumHD clusters. Clusters 2 and 5 are highlighted as being assigned a dual name due to registration to two snMultiome clusters. (**B**) Spatial registration of *BANKSY* clusters to the fine-resolution snMultiome clusters help disambiguate clusters 8 and 15 as MHb_A and MHb_B, respectively, despite ambiguous mapping to the mid-resolution clusters. (**C**) Cluster 9, despite clean registration to the GABA_LHb_C.2 cluster, was assigned the dual name "Excit.Thal/GABA_LHb_C.2" due to its notable spatial location nearly exclusively in the thalamus. (**D**) Clusters 13 and 14 were annotated as subependymal due to spatial adjacency with the annotated ependymal cluster. (**E**) Cluster 16 was present in a small portion of primarily Br9902, shown in section Br9902_1.

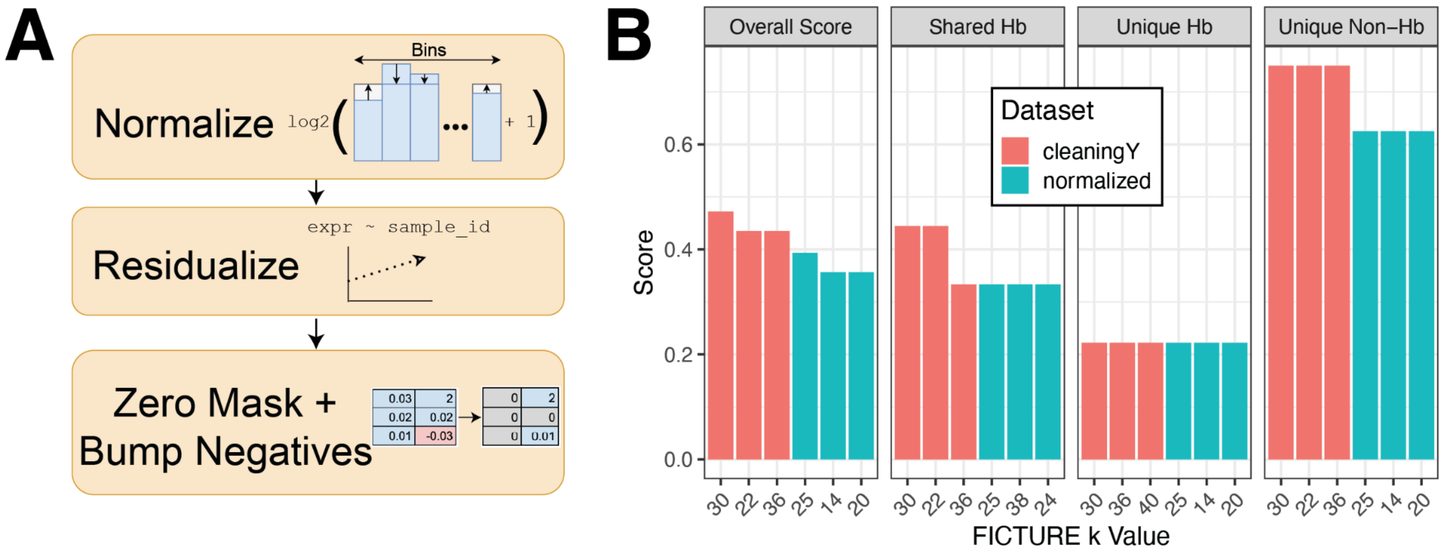


**Fig. S17: Batch correction improves interpretability of *FICTURE* results**. (**A**) Batch correction consists of log-normalization, residualization using a linear model of expression against sample ID, and an adjustment step that preserves initial zeros and prevents introduction of negative counts. (**B**) Summary scores for the top-3-scoring numbers of factors in batch-corrected ("cleaningY") and non-batch-corrected ("normalized") *FICTURE* results. The three metrics and their average are described in **Methods 4.23**.

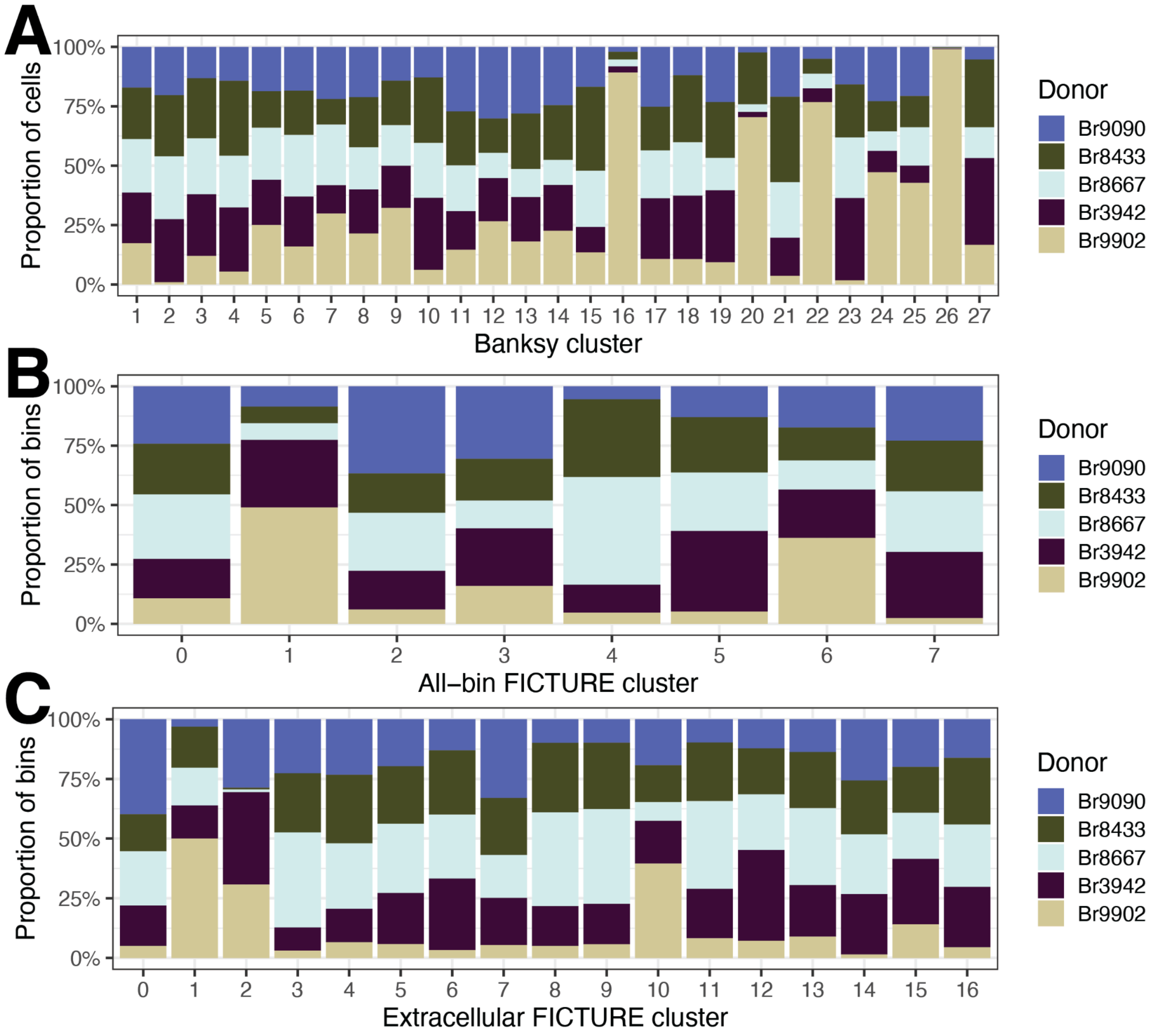


**Fig. S18: Donor composition across VisiumHD *BANKSY* and *FICTURE* clusters**. (**A**) Proportion of cells from each *BANKSY* cluster, prior to cell-type annotation, belonging to each donor. (**B**) Proportion of 2 μm bins from each $k$ = 8 all-bin *FICTURE* cluster belonging to each donor. (**C**) Proportion of 2 μm bins from each $k$ = 17 extracellular *FICTURE* cluster belonging to each donor.

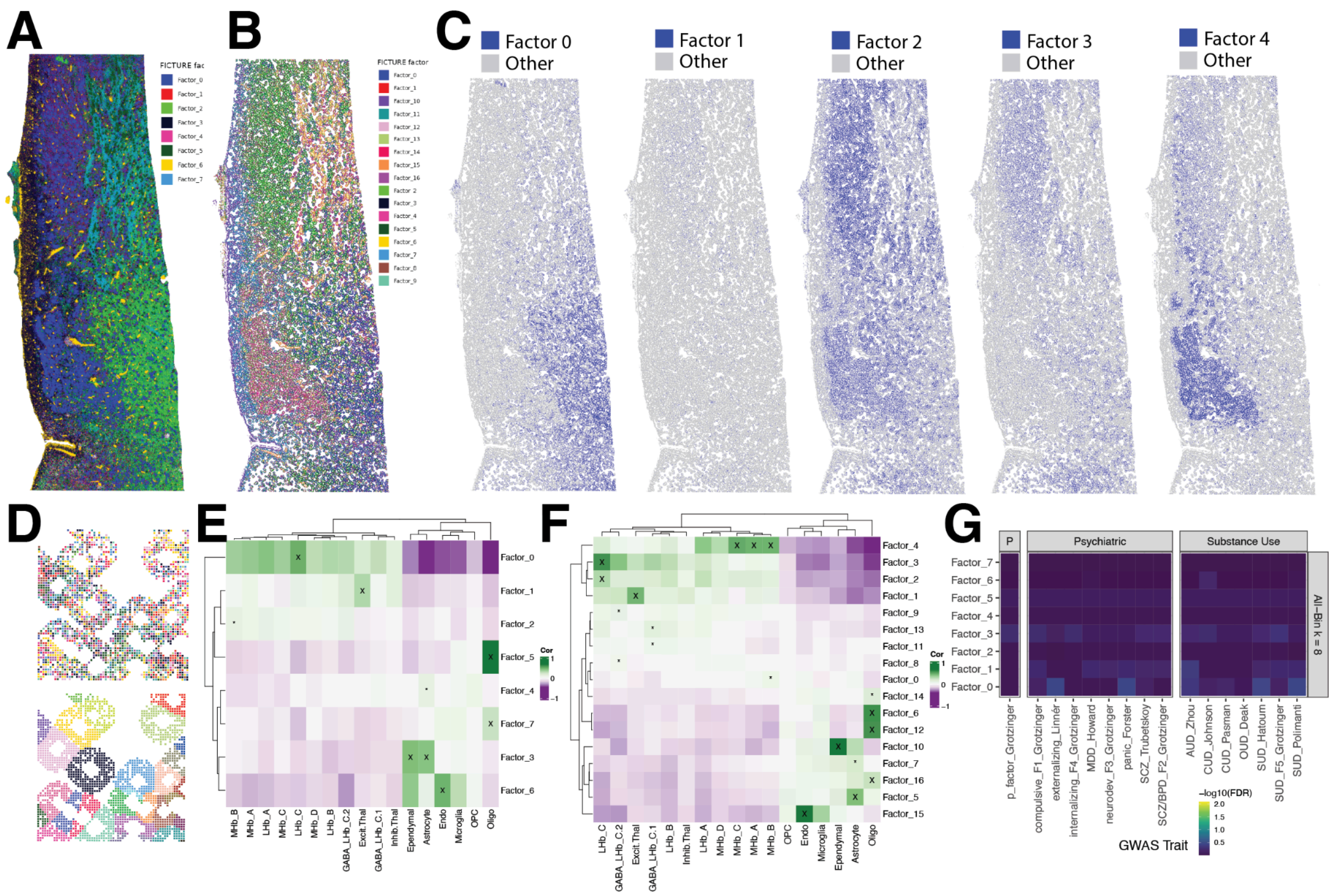


**Fig. S19: Characterization of VisiumHD all-bins and extracellular *FICTURE* clusters.** (**A**) Br9090_1 spatial plot with 2 µm bins colored by *FICTURE* factors 0 to 7 identified using all the bins (all bins shown in **Fig. S14A** and described in **Methods 4.22**). Factors 5 and 7 show white matter tracts. (**B**) Similar to (A), but for *FICTURE* analysis using only extracellular bins (orange bins shown in **Fig. S14A**) that identified 17 factors. (**C**) Similar to (B), but showing *FICTURE* factors 0 to 4 individually with all other factors in grey. (**D**) A zoomed-in region of Br9090_1 with 2 µm bins colored by the extracellular-only *FICTURE* factors (top) or the cell they are associated with (bottom). This shows that the extracellular-only *FICTURE* analysis (top, **Methods 4.22**) allows more refined assignments of the 2 µm extracellular bins than the *BANKSY* extracellular bin analysis (bottom). The latter *BANKSY* analysis was performed by aggregating expression from extracellular bins by their corresponding cells and assigning the cells labels determined in the cellular clustering and annotation (**Methods 4.23**). (**E**) Transcriptional cluster registration between Hb snMultiome clusters (columns) and all-bin *FICTURE* factors from (A) (rows). Correlations are computed across transcriptome-wide 1-vs-All DE *t*-statistics to identify transcriptionally similar clusters, with the ‘x’ representing high confidence matches (**Methods 4.22**). (**F**) Similar to (E), but with the extracellular-only *FICTURE* factors from (B) (rows). (**G**) Heatmap of -$\log_{10}$ FDR-adjusted p-values showing whether top 200 MeanRatio marker genes per all-bin *FICTURE* factors (A) are enriched in risk genes from different GWASes. ‘*’ denotes FDR < 0.05 with FDR computed by cluster across GWASes. GWASes were grouped by broad categories: psychiatric disorders, substance use disorders, or P factor (Grotzinger et al. 2026).

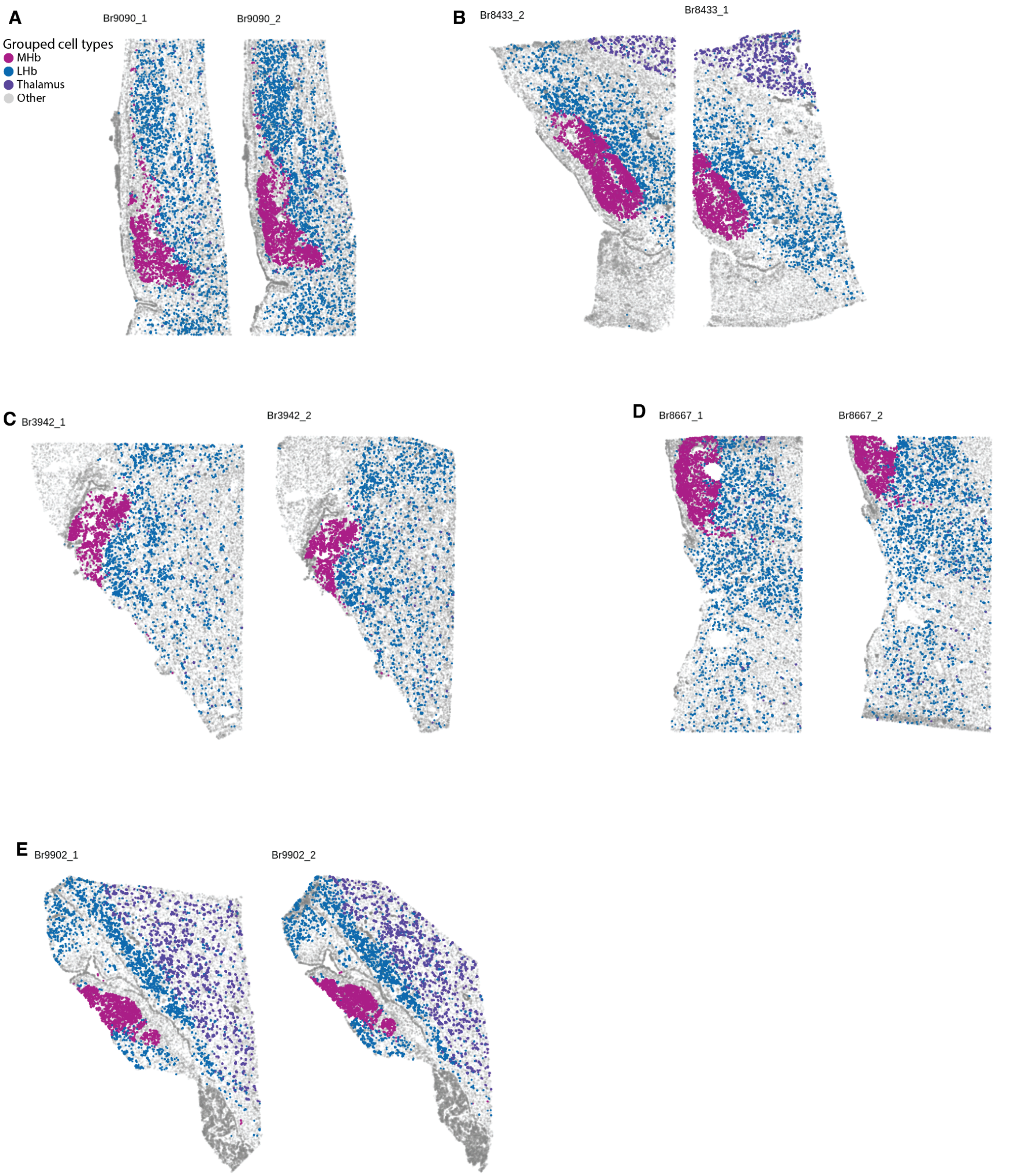


**Fig. S20: Broad MHb, LHb, and thalamus spatial clusters.** (**A**) Spatial plot of segmented VisiumHD cells for donor Br9090. Cells are labeled by collapsed *BANKSY* clusters (**Fig. 5A**) for the MHb, LHb, and thalamus. (**B**) Same as (A), for donor Br8433. (**C**) Same as (A), for donor Br3942. (**D**) Same as (A), for donor Br8667. (**E**) Same as (A), for donor Br9902.

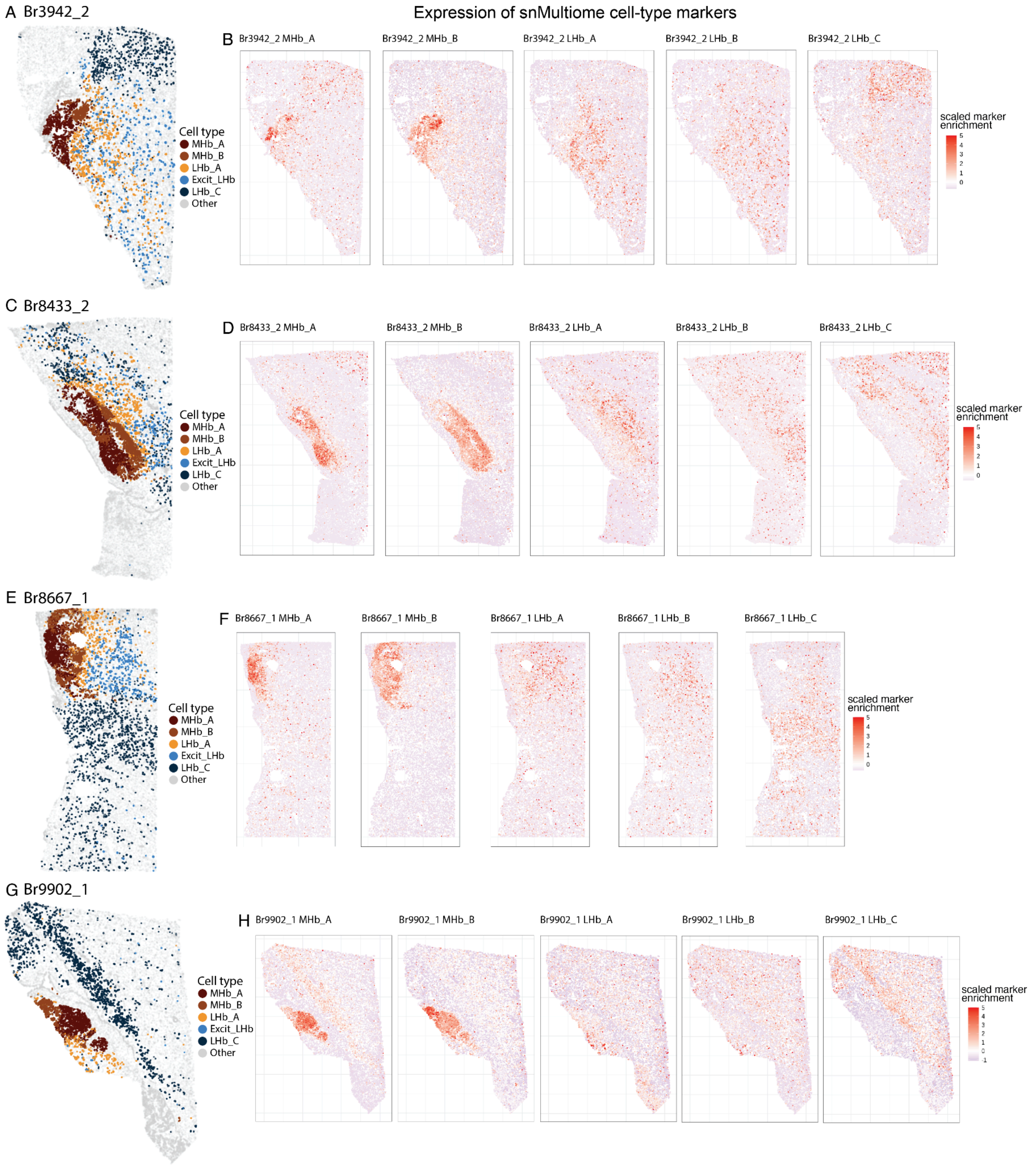


**Fig. S21: Spatial cluster plots of LHb and MHb neurons across all donors.** (**A**) Spatial plot of segmented VisiumHD cells for Br3942. *BANKSY* clusters annotated as Hb cell types are labeled in color. (**B**) Spatial plot of segmented VisiumHD cells (Br3942 donor) colored by aggregate marker gene expression of MHb_A, MHb_B, LHb_A, LHb_B, and LHb_C (left to right) snMultiome clusters. (**C**) Same as (A), but for Br8433. (**D**) Same as (B), but for Br8433. (**E**) Same as (A), but for Br8667. (**F**) Same as (B), but for Br8667. (**G**) Same as (A), but for Br9902. (**H**) Same as (B), but for Br9902.

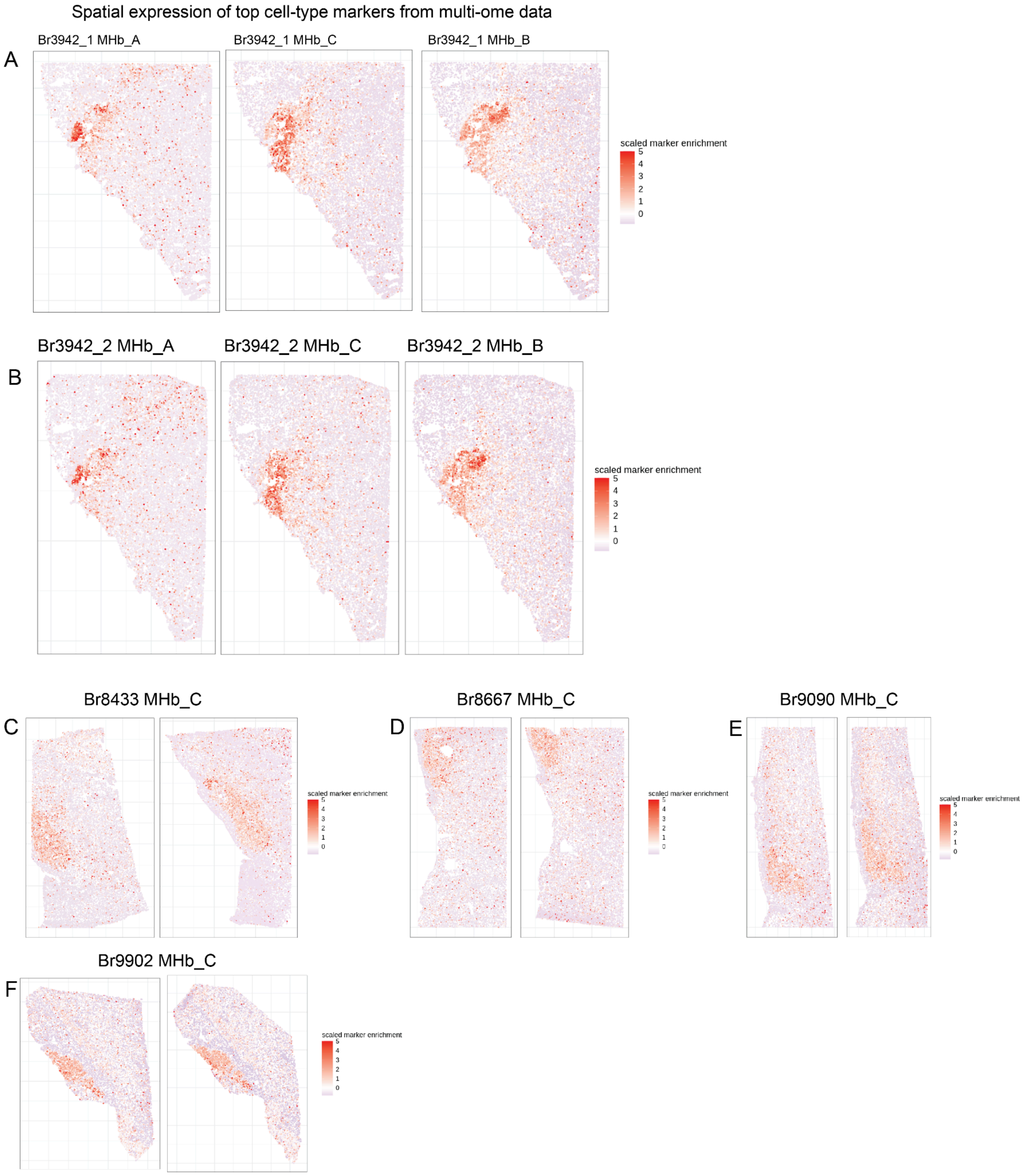


**Fig. S22: MHb_C snRNA-seq gene markers projected into VisiumHD data.** (**A**) Spatial plot of segmented VisiumHD cells colored by aggregate marker gene expression of MHb_A, MHb_C, and MHb_B (left to right) snMultiome cell types for Br3942. Aggregate expression of MHb_C is localized between MHb_A and MHb_B domains. (**B**) Same as (A), but for the second VisiumHD section for Br3942 (**C**) Same as A, but for Br8433 and just for MHb_C aggregate marker gene expression, depicting no clear MHb_C domain as in Br3942. (**D**) Same as (C), but for Br8667 . (**E**) Same as (B), but for Br9090. (**F**) Same as (B), but for Br9902.

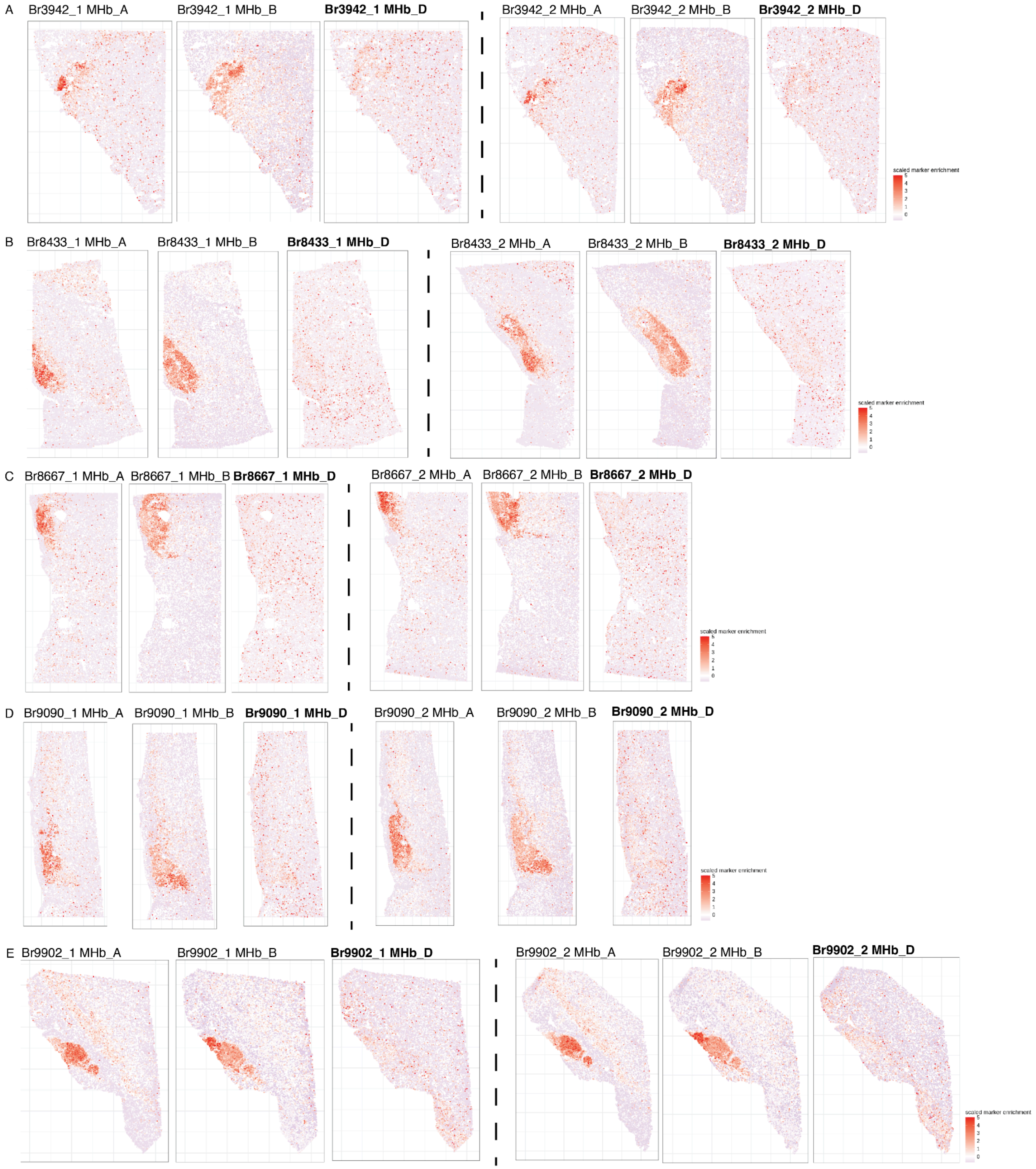


**Fig. S23: No clear spatial organization for MHb_D snMultiome gene marker expression.** (**A**) Spatial plot of segmented VisiumHD cells inBr3942 colored by aggregate marker gene expression of MHb_A, MHb_B, and MHb_D (left to right) snMultiome clusters. Similar plots are shown for (**B**) Br8433, (**C**) Br8667, (**D**) Br9090, and (**E**) Br9902.

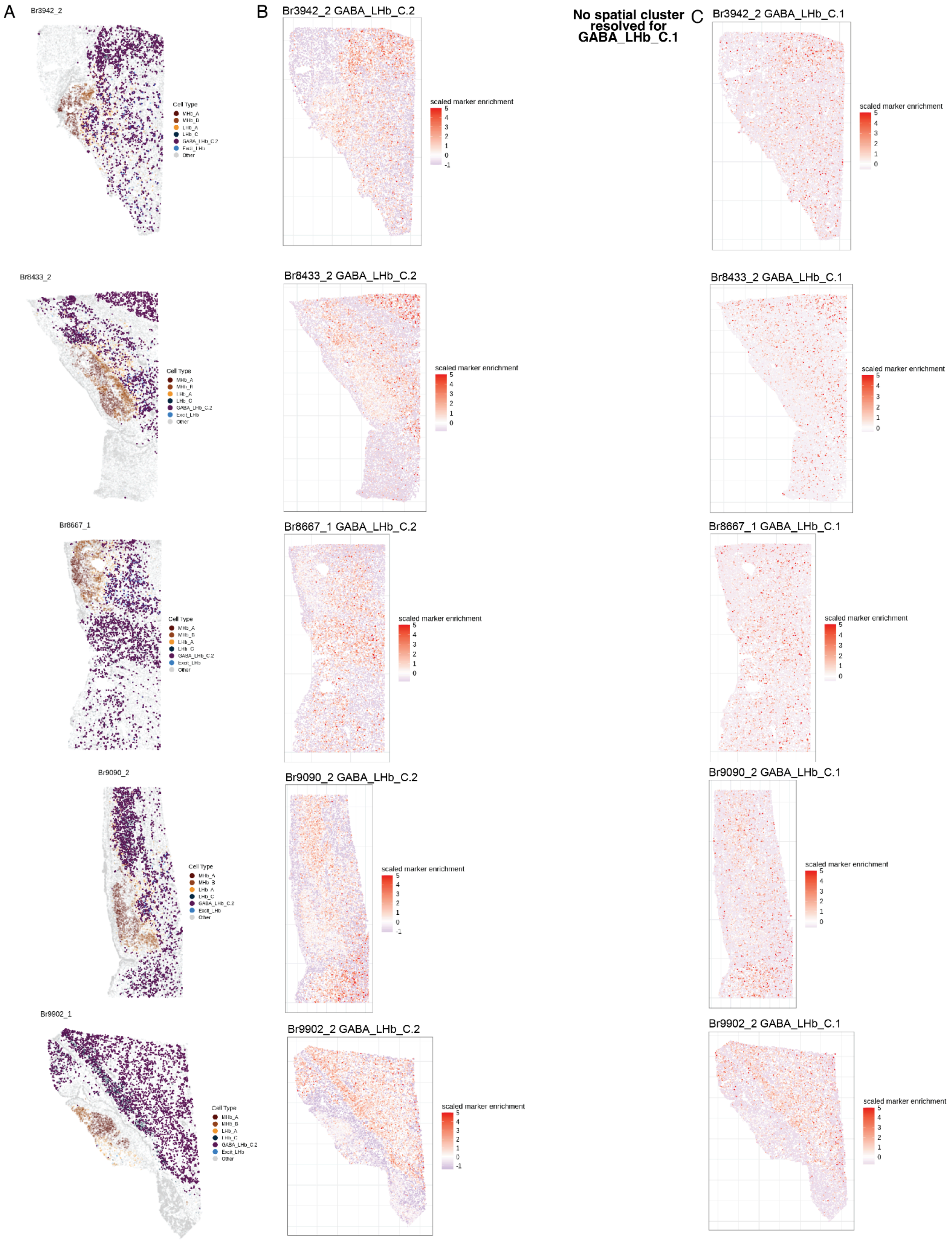
A
Br3942_2
Cell Type
MHb_A
MHb_B
LHb_A
LHb_C
GABA_LHb_C.2
Excit_LHb
Other
Br8433_2
Br8667_1
Br9090_2
Br9902_1
B
Br3942_2 GABA_LHb_C.2
scaled marker enrichment
5
4
3
2
1
0
-1
Br8433_2 GABA_LHb_C.2
Br8667_1 GABA_LHb_C.2
Br9090_2 GABA_LHb_C.2
Br9902_2 GABA_LHb_C.2
No spatial cluster resolved for GABA_LHb_C.1
C
Br3942_2 GABA_LHb_C.1
Br8433_2 GABA_LHb_C.1
Br8667_1 GABA_LHb_C.1
Br9090_2 GABA_LHb_C.1
Br9902_2 GABA_LHb_C.1

**Fig. S24: Gene marker projection of GABAergic LHb clusters onto VisiumHD data.** (**A**) Spatial plot of segmented VisiumHD cells for Br3942_2, Br8433_2, Br8667_1, Br9090_2, and Br9902_1. *BANKSY* clusters annotated as Hb cell types are labeled in color. (**B**) Spatial plot of segmented VisiumHD cells for the same donors in (A), colored by aggregate marker gene expression of the GABA_LHb_C.2 snMultiome cluster. (**C**) Similar to (B), but for GABA_LHb_C.1 expression.

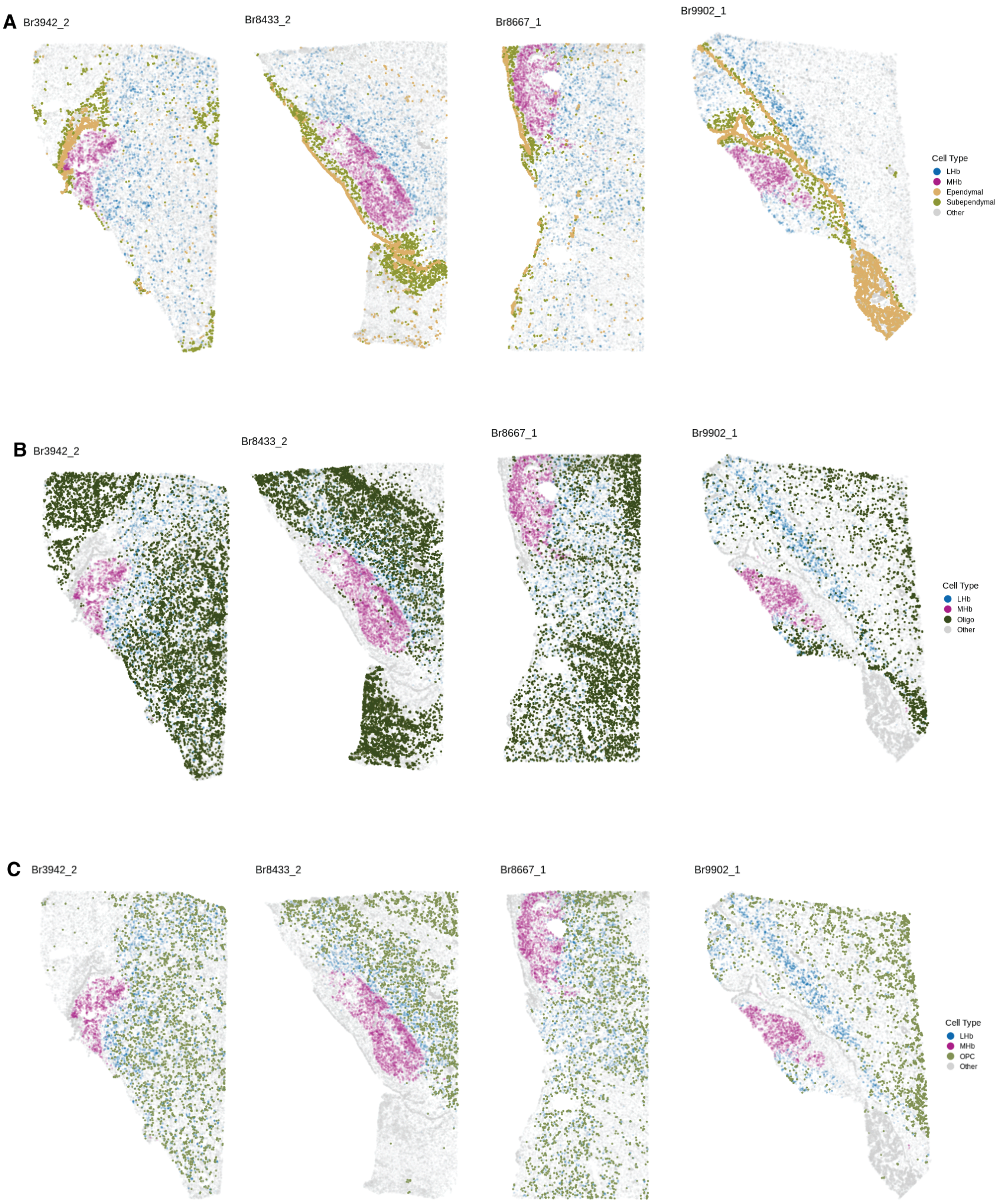


**Fig. S25: Spatial plots of non-neuronal Hb cell types.** (**A**) Spatial plot of segmented VisiumHD cells for Br3942_2, Br8433_2, Br8667_1, Br9090_2, and Br9902_1. Annotated ependymal and subependymal *BANKSY* clusters are in full color, with collapsed *BANKSY* clusters for MHb and LHb cell types in transparent color. (**B**) Same as in (A), but for the Oligo *BANKSY* cluster. (**C**) Same as in (A), but for the OPC *BANKSY* cluster.

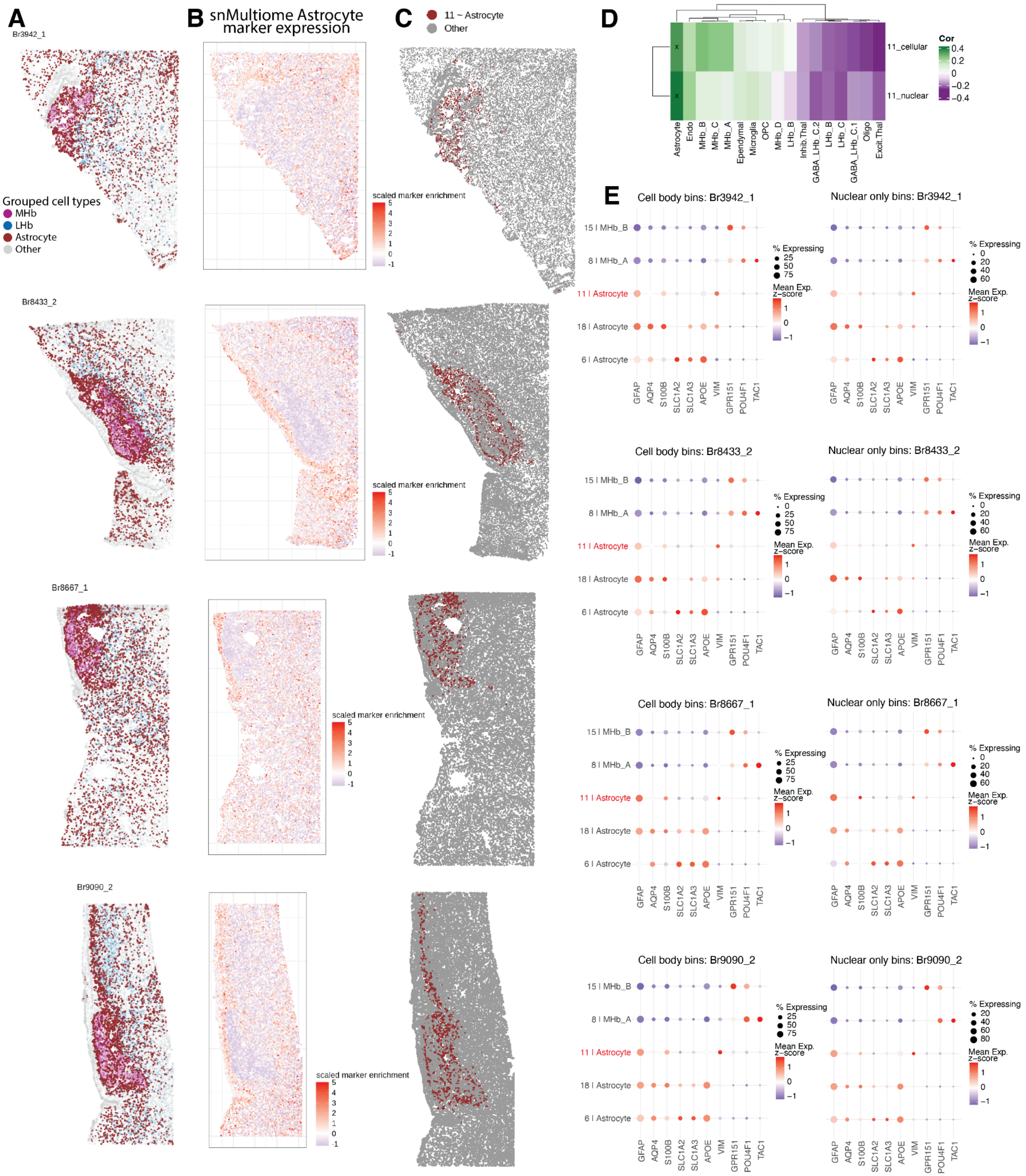
A
B snMultiome Astrocyte marker expression
C
D
E
Br3942_1
Br8433_2
Br8667_1
Br9090_2
Grouped cell types
MHb
LHb
Astrocyte
Other
scaled marker enrichment
11 ~ Astrocyte
Other
11_cellular
11_nuclear
Cor
Astrocyte
Endo
MHb_B
MHb_C
MHb_A
Ependymal
Microglia
OPC
MHb_D
LHb_B
Inhib.Thal
GABA_LHb_C.2
LHb_B
LHb_C
GABA_LHb_C.1
Oligo
Excit.Thal
Cell body bins: Br3942_1
Nuclear only bins: Br3942_1
Cell body bins: Br8433_2
Nuclear only bins: Br8433_2
Cell body bins: Br8667_1
Nuclear only bins: Br8667_1
Cell body bins: Br9090_2
Nuclear only bins: Br9090_2
15 | MHb_B
8 | MHb_A
11 | Astrocyte
18 | Astrocyte
6 | Astrocyte
% Expressing
Mean Exp. z-score
GFAP
AQP4
S100B
SLC1A2
SLC1A3
APOE
VIM
GPR151
POU4F1
TAC1

**Fig. S26: Discrepancy in spatial expression of snMultiome Astrocyte gene markers.** (**A**) Spatial plot of segmented VisiumHD cells for Br3942_2, Br8433_2, Br8667_1, Br9090_2, and Br9902_1. Annotated astrocyte *BANKSY* clusters are in full color, with the collapsed *BANKSY* clusters for MHb and LHb cell types in transparent color. (**B**) Spatial plot of segmented VisiumHD cells for the same donors in (A), colored by aggregate marker gene expression of the Astrocyte snMultiome cluster. (**C**) Spatial plot of segmented VisiumHD cells for the same donors in (A), but only the MHb localized fine-resolution astrocyte *BANKSY* cluster (cluster 11) in color. (**D**) Transcriptional cluster registration between the Hb snMultiome clusters (columns) and *BANKSY* cluster 11 (rows), either with cellular- or nuclear-segmented VisiumHD bins. Restricting to just the nuclear-segmented bins increases the transcriptional correlation of *BANKSY* cluster 11 to the snMultiome astrocyte cluster. (**E**) Bubble plots of astrocyte and MHb marker gene expression across the 3 fine-resolution *BANKSY* astrocyte clusters and the MHb_A and MHb_B *BANKSY* MHb clusters for the same donors as in (A). The left column of bubble plots is using the cellular-segmented VisiumHD bins and the right column is using the nuclear-segmented VisiumHD bins.

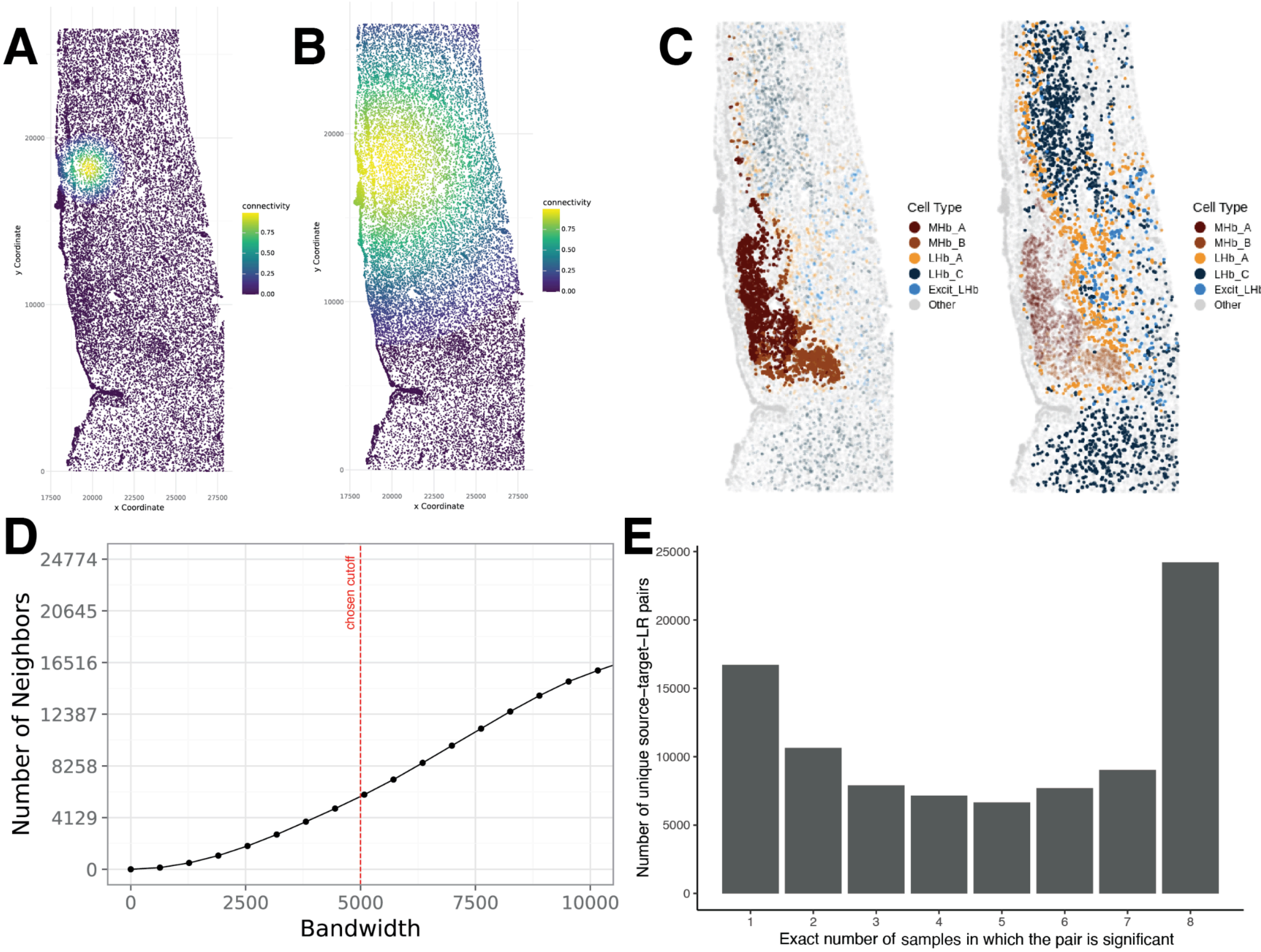


**Fig. S27: Cell to cell spatially-informed communication threshold determination for *LIANA+ InFlow* analyses.** (**A**) Connectivity gradient spatial map for *LIANA+ InFlow* bandwidth set at 1,000 pixels (~250 μm) for Br9090_2 with center at (19756, 23382). (**B**) Similar to (A), but for bandwidth set at 5,000 pixels (~1250 μm). (**C**) Cells in (A) colored by MHb and LHb cell types (identical to **Fig. 5B**). At bandwidth 5,000 pixels (B), the connectivity gradient allows connections across the whole Hb. (**D**) Average number of cell neighbors considered for Br9090_1 across *InFlow* bandwidth values. (**E**) Distribution of the number of unique significant (FDR < 0.05) source-target Ligand-Receptor (L-R) pairs across different thresholds for the number of samples in which the pair is significant. A threshold of 4 samples was chosen to identify replicable source-target L-R pairs.

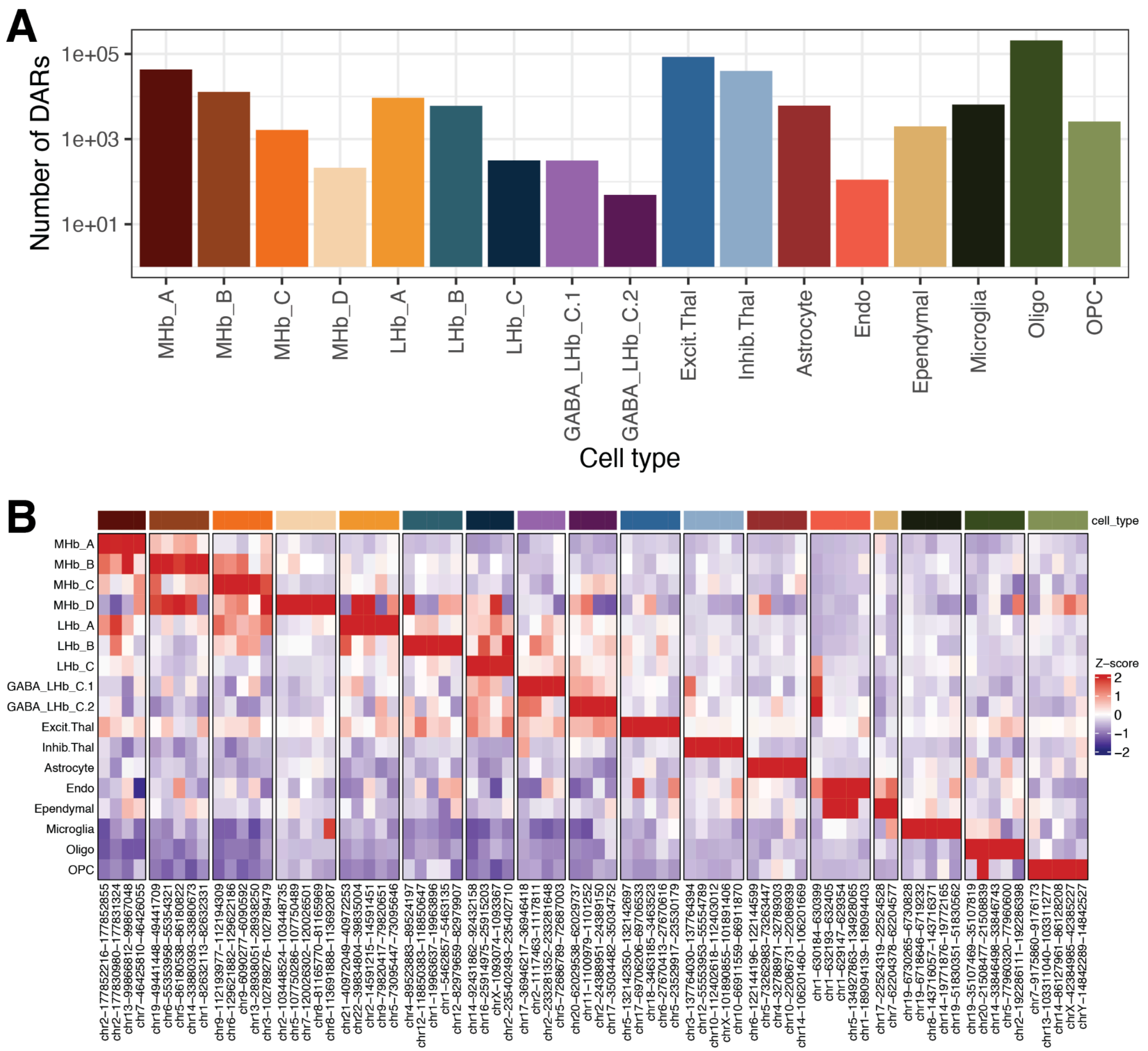


**Fig. S28: Differentially accessible regions (DARs) by snMultiome cell type.** (**A**) Distribution of the number of DARs by cell type. The *Y*-axis is shown in a $\log_{10}$ scale. (**B**) Heatmap showing up to the top 5 open DARs by cell type (higher counts in target cell type compared to all others). Hg38 genomic coordinates are shown in the *X*-axis. The ATAC normalized counts were *Z*-scored by row.

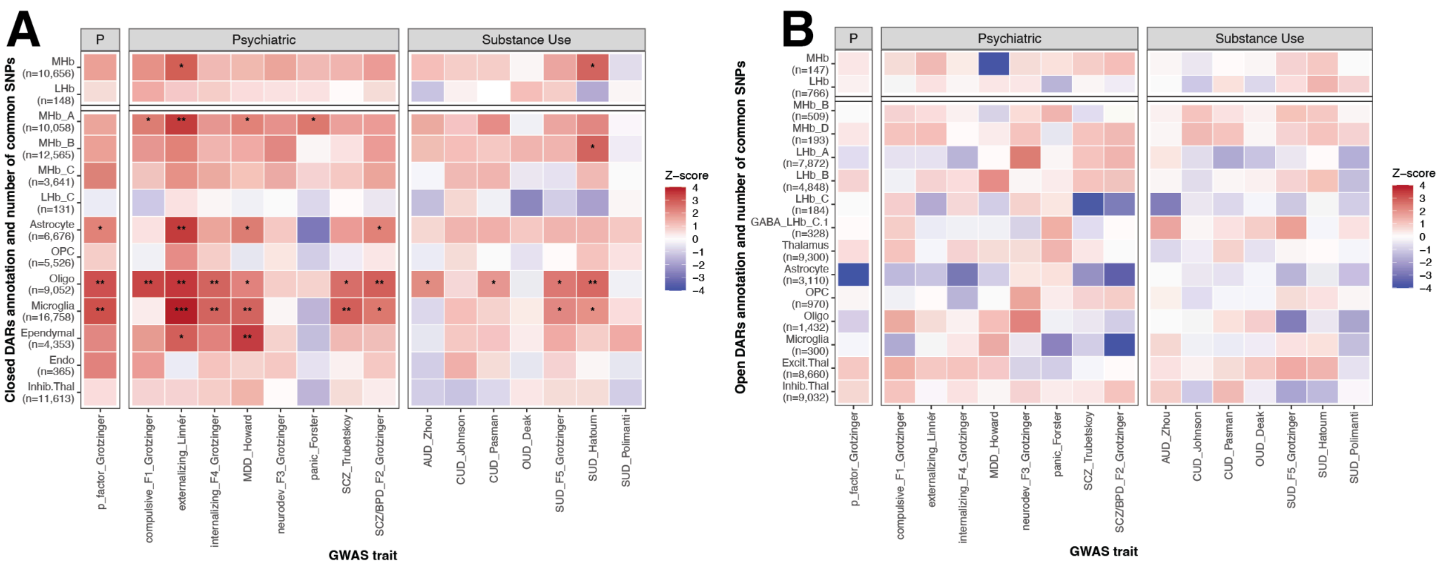


**Fig. S29: Association between closed (less accessible) and open (more accessible) cell type DARs with disorder-associated genetic variants.** Heatmaps of stratified linkage disequilibrium score regression (*S-LDSC*) associations between cell type DARs and disorder-associated genetic variants from GWAS datasets: (**A**) closed DARs (less accessible in target cell type compared to all other ones), (**B**) open DARs (more accessible in target cell type). '*' denotes FDR < 0.05 with FDR computed by cell type across GWASes. Related to **Fig. 7D**.

### Supplementary Tables

**Table S1: Donor demographics.** Columns include donor identifier (donor), whether the donor is analyzed in Visium H&E, Visium HD, and snMultiome (in_visium_HE, in_visium_HD, and in_multiome, respectively), age of death (age), ancestry (ancestry), biological sex (sex), and post-mortem interval (PMI).

**Table S2: snMultiome *Cell Ranger ARC* summary information, experimental metrics, and post-QC number of cells.** num_neuronal_nuclei_sorted, num_glial_nuclei_sorted, num_neurons_loaded, num_glia_loaded, concentration_post_nuclei_permeabilization_nuclei_per_uL, and total_nuclei_loaded are experimental metrics measuring number of sorted neuronal nuclei, number of sorted glial nuclei, number of loaded neurons, number of loaded glia, concentration of nuclei following permeabilization (in nuclei/µL), and number of loaded nuclei total, respectively. num_cells_post_QC is the number of cells after cellranger-arc reanalyze and quality control. All other columns are output from *Cell Ranger ARC* summaries. **Related to Methods 4.4**.

**Table S3: Habenula marker genes.** Hb marker genes were either "Literature-Based" (LB) or "Data-Driven" (DD) (Yalcinbas et al. 2025). Related to **Methods 4.5**. Columns include the gene symbol gene, cell type for which the gene is a marker cell_type, and marker_category, the categorization as either literature-based or data-driven.

**Table S4: Wallace mouse habenula cell-type annotations.** Author provided annotations are in the original_annot and original_subHab_annot columns (Wallace et al. 2020). The de novo sample-specific clusters, *MetaNeighbor* (Crow et al. 2018; Fischer et al. 2021) identified meta-clusters, annotated meta-clusters, and consensus annotations are provided in the de_novo_cluster, meta_cluster, annotated_meta_cluster, and consensus_annot columns, respectively. Related to **Methods 4.8**.

**Table S5: Hashikawa mouse habenula cell-type annotations.** Author provided annotations are in the original_annot column (Hashikawa et al. 2020). The annotated meta-clusters matched to the Wallace dataset meta-clusters are in the annotated_meta_cluster column, and the consensus annotations are in the consensus_annot column. Related to **Methods 4.8**.

**Table S6: Pandey zebrafish habenula cell-type annotations.** Column titles are similar to **Table S4** (Pandey et al. 2018). Related to **Methods 4.8**.

**Table S7: Yalcinbas human habenula cell-type annotations.** Column titles are similar to **Table S4** (Yalcinbas et al. 2025). Related to **Methods 4.8**.

**Table S8: Multiome human habenula cell-type annotations.** The initial fine-resolution annotations (**Fig. S5**) are in the initial_fine_res column. The adjusted fine-resolution annotations (**Fig. S5**) are in the refined_fine_res column. The mid-resolution annotations (**Fig. 1**) before cross-species alignment are in the mid_res column. And the consensus annotations (**Fig. 2**) are in the consensus_annot column. Related to **Methods 4.7,4.8**.

**Table S9: Data driven marker genes for Hb cell types.** The top ~50 1-vs-All markers (Huuki-Myers, Montgomery, et al. 2025) used for the various marker projections to mouse and spatial data. Further described in **Results 2.4**. Duplicate markers were filtered as described in **Methods 4.8**.

**Table S10: All level 1, level 2, and intersecting trios for the snMultiome cell types.** Trios were computed using *TRIPOD* (Jiang et al. 2022). Columns include the ATAC peak coordinates (`peak`), gene symbol (`gene`), transcription factor gene symbol (`TF`), snMultiome cluster (`mid_cluster`), level 1 and level 2 coefficients and p-values (`trio_coef_lvl1`, `trio_coef_lvl2`, `trio_p_adj_lvl1`, and `trio_p_adj_lvl2`), and the average log-normalized expression across metacells of the same cluster for the gene (`gene_avg_expr`) and TF (`TF_avg_expr`). Related to **Methods 4.14**.

**Table S11: Genes and gene ontology enrichment terms for astrocyte DE.** Significant (FDR < 0.05) GO terms enriched using *clusterProfiler* (Xu et al. 2024) among MHb vs. LHb astrocyte differentially expressed genes are enumerated, along with each term's constituent genes. Columns include the gene ontology term ID (`term_id`), its corresponding description (`term_description`), gene ENSEMBL ID (`gene_id`), Hb region where astrocytes had higher expression of the gene (`DE_higher_in`), and FDR-adjusted enrichment p-value (`p.adjust`). Related to **Methods 4.27**.

**Table S12: MHb vs. LHb astrocyte DE statistics for all tested genes.** Fold changes are oriented such that positive values indicate higher expression in MHb. Columns include the gene ENSEMBL ID and symbol (`gene_id` and `gene_name` respectively), DE *t*-statistic (`t`), DE log fold change (`logFC`), nominal and FDR-adjusted p-values (`p` and `fdr`, respectively), and nominal and FDR-adjusted empirical p-values (`p_empirical` and `fdr_empirical`, respectively, **Methods: 4.27**) computed with *edgeR* (Chen et al. 2025) and *limma* (Ritchie et al. 2015).

**Table S13: Up to top 200 *MeanRatio* markers for each VisiumHD and snMultiome dataset in the study.** Markers are included for all-bin `k = 8` *FICTURE* (Si et al. 2024), extracellular *FICTURE* `k = 17`, VisiumHD cell types, VisiumHD extracellular cell types, and snMultiome cell types, which are described in the `dataset` column. Other columns include `set_id`, the cluster identity, `gene_id` and `gene_name` for the gene ENSEMBL ID and symbol respectively, and the *MeanRatio* (Huuki-Myers, Montgomery, et al. 2025) of that cluster against the next-highest-expressing cluster (`mean_ratio`). Marker computation is further described in **Methods 4.11**.

**Table S14: Visium H&E *SpaceRanger* summary information.** All columns are produced by *SpaceRanger* except `num_spots_post_QC`, which is the number of spots present per sample after quality control. Related to **Methods 4.17**.

**Table S15: Visium HD *SpaceRanger* summary information.** All columns are produced by *SpaceRanger* except `num_cells_post_QC`, which is the number of cells present per donor after quality control. Related to **Methods 4.20**.

**Table S16: Summary of software details for Visium HD analysis.** Columns include the name of the software tool (`software_name`), whether it natively supports operating on multiple samples (capture areas) simultaneously (`multi_sample`), its typical run time for a single capture area (`run_time`), required virtual memory in gigabytes (`required_mem_gb`), programming language (`language`), and additional free-form notes providing further details (notes). The use of each software tool is further described in **Methods 22-23, 25-26**.

**Table S17: Significant ligand-receptor interactions.** Global communication scores for ligand-receptor pairs between pairs of interacting cell types are included for interactions significant ($p < 0.05$) in at least 4 of 8 VisiumHD samples, computed with *LIANA+* (Dimitrov et al. 2024). Columns include the source and target cell types (source and target), ligand and receptor genes (ligand and receptor), ligand-receptor pair (lr_pair), source-target cell-type pair (source_target), number of samples in which the interaction was significant (n_samples_significant), identifiers of those samples (sample_ids), and the average communication score (global_communication_score). Related to **Methods 4.26**.

**Table S18: Genes and gene ontology enrichment terms for cell-cell-communication results.** Significant (FDR < 0.05) terms and their constituent ligand or receptor genes are provided for enrichment tests performed for each pair of interacting cell types using *clusterProfiler* (Xu et al. 2024). Columns include the source and target cell types (source and target), Gene Ontology category (ontology), GO term identifier and description (term_id and term_description), constituent gene (gene), its role in the cell-cell communication interaction (role; ligand, receptor, or ligand/receptor), gene ratio and background ratio used for enrichment testing (GeneRatio and BgRatio), number of genes associated with the term (Count), enrichment p-value (pvalue), and FDR-adjusted p-value (FDR). Related to **Methods 4.26**.

**Table S19: Gene-level p-values for enrichment of each cluster for each GWAS trait.** Only genes comprising significantly enriched (FDR < 0.05) marker sets from **Table S13** are included. Columns dataset, set_id, gene_id, and gene_name are shared with **Table S13**. Also included are the GWAS tested for enrichment stats (gwas) and the gene-level enrichment p-value (p) from *MAGMA* (Leeuw et al. 2015). Related to **Methods 4.28**.

**Table S20: *S-LDSC* enrichment of DARs.** Columns included the GWAS phenotype (GWAS_label), DAR set (dar_set; open, closed, or all DARs), cell type (cell_type), number and proportion of SNPs in each annotation (n_snps and Prop._SNPs), proportion of heritability and its standard error (Prop._h2 and Prop._h2_std_error), enrichment and its standard error (Enrichment and Enrichment_std_error), *S-LDSC* coefficient *z*-score (coefficient_z) (Bulik-Sullivan et al. 2015), one-sided coefficient p-value (coefficient_p_one_sided), and FDR-adjusted p-value (coefficient_FDR). Related to **Methods 4.29**.

**Table S21: All significant (FDR < 0.05) open and closed DARs for each snMultiome cell type.** Columns include the ATAC peak coordinates (peak), snMultiome cluster (cell_type), differential accessibility direction ("up" for open and "down" for closed, DA_direction), average log fold change for the cluster against all others (avg_log2FC), and the nominal and FDR-corrected p-values (p_val and p_adj, respectively) computed with *Seurat* (Stuart et al. 2019). Related to **Methods 4.30**.
